# Guidelines for Optimizing Type S Non-Ribosomal Peptide Synthetases

**DOI:** 10.1101/2023.03.21.533600

**Authors:** Nadya Abbood, Juliana Effert, Kenan A. J. Bozhueyuek, Helge B. Bode

## Abstract

Bacterial biosynthetic assembly lines, such as non-ribosomal peptide synthetases (NRPS) and polyketide synthases, are often subject of synthetic biology – because they produce a variety of natural products invaluable for modern pharmacotherapy. Acquiring the ability to engineer these biosynthetic assembly lines allows the production of artificial non-ribosomal peptides (NRP), polyketides, and hybrids thereof with new or improved properties. However, traditional bioengineering approaches have suffered for decades from their very limited applicability and, unlike combinatorial chemistry, are stigmatized as inefficient because they cannot be linked to the high-throughput screening platforms of the pharmaceutical industry. Although combinatorial chemistry can generate new molecules cheaper, faster, and in greater numbers than traditional natural product discovery and bioengineering approaches, it does not meet current medical needs because it covers only a limited biologically relevant chemical space. Hence, methods for high-throughput generation of new natural product-like compound libraries could provide a new avenue towards the identification of new lead compounds. To this end, prior to this work, we introduced an artificial synthetic NRPS type, referred to as type S NRPS, to provide a first-of-its-kind bicombinatorial approach to parallelized high-throughput NRP library generation. However, a bottleneck of these first two generations of type S NRPS was a significant drop in production yields. To address this issue, we applied an iterative optimization process that enabled titer increases of up to 55-fold compared to the non-optimized equivalents, restoring them to wild-type levels and beyond.

## Introduction

Synthetic biology (SynBio) seeks to create or redesign systems and cells with molecular biology tools and genome editing technologies to produce novel biological parts, such as molecules, materials, or cells. For example, synthetic biology holds the potential to bio-synthesize drugs, chemicals, and bio-fuels more sustainably and cost-effectively than with traditional methods.^1, 2^ Multimodular enzyme complexes, such as polyketide synthases and non-ribosomal peptide synthetases (NRPSs), are often subject of SynBio – because they produce a variety of valuable chemicals or pharmaceutically relevant peptides. Engineering these biosynthetic assembly lines can produce artificial polyketides, non-ribosomal peptides (NRPs), or hybrids thereof with new or improved properties. ^3^

Recently, we have focused in particular on developing more efficient SynBio methods for engineering modular NRPSs (type I NRPS),^4, 5^ which has been in its infancy for decades.^6, 7^ By introducing the exchange Unit (XU) and the XU Condensation Domain (XUC) concepts, we provided the necessary means to enable more reproducible, predictable, and robust engineering than before.^4, 5^ The XU concept, for example, leverages a fusion point located between the NRPS’s condensation (C) and adenylation (A) domains within the structurally flexible region of the C-A interdomain linker.^4^ A domains in NRPSs are responsible for ATP-dependent selection and activation of specific amino acids (AAs), while C-domains catalyze peptide-bond formation between two AA residues. Together with the T domain, which transports the activated AA from the A-domain to the C-domain, they form the core domains of a canonical minimal NRPS module (C-A-T).^8^ NRPSs usually consist of not just one but several sequentially repeating modules, each responsible for the incorporation and optional functional group modification of a specific AA. Commonly, the number of modules in a NRPS corresponds directly to the number of AAs incorporated into the associated NRP.^8, 9^ Of note, Dell et al., re-categorized ribosome-independent peptide assembly systems into five groups (type I, II, III, IV, and V) and assigned NRPSs to type I and type II. Type I and II are distinguished by their modular or non-modular/freestanding character, with modular NRPSs classified as type I and freestanding NRPSs as type II NRPSs.^10^ Iterative systems, such as the enniatin^11^ or rhabdopeptide^12^ producing NRPSs that reuse individual modules, no longer represent a separate group but are assigned to type I NRPSs.^9, 10^

Despite the progress made, assembly line engineering remained difficult and time consuming due to their mere size and repetitive nature.^7^ Traditional NRPS cloning and engineering often requires elaborated cloning strategies such as yeast cloning, LLHR^13^ (Linear plus linear homologous recombination-mediated recombineering), or ExoCET^14^ (Exonuclease Combined with RecET recombination) which are frequently accompanied by technical limitations.^7^ Therefore, we established a new synthetic type of NRPS (type S) that allows easier and faster cloning by splitting large single-protein multi-modular NRPSs into two or three smaller and independently expressible subunits that are reconstituted to full-length in the living cell via the interaction of high-affinity leucine zippers (SYNZIPs).^15, 16^ Further studies have recently used the same or similar high-affinity tags, e.g., SpyTag/SpyCatcher^17^, zinc fingers^18^ and SYNZIPSs^19^ (SZs), to mediate proteinprotein interaction of split NRPS or polyketide synthases (PKSs). Introduction of these protein tags has increased productions titers (in vanlinomycin synthesis)^17^, mediated DNA-protein recognition (of DNA-templated NRPS)^18^, or enabled generation of chimeric PKS (of 6-deoxyerythronolide B synthase)^19^, emphasizing the diverse applicability of the various interaction toolboxes for SynBio applications.

The ability to separate NRPS-encoding biosynthetic gene clusters into smaller DNA fragments that encode partial NRPS proteins (subunits) and then distribute the respective gene fragments onto different expression plasmids naturally simplifies cloning – making ‘standard’ *in vitro* cloning strategies such as Gibson,^20^ HiFi and Hot Fusion^21^ (Isothermal-) assembly sufficient. To reconstitute the communication of generated NRPS protein subunits, we attached SZs that post-translationally restore the full-length biosynthetic capacity of the modular NRPS system in the living cell.^15, 16, 22, 23^ However, type S NRPSs not only simplify NRPS engineering, but also offer, for the first time, the possibility of true bio-combinatorial approaches to the design of natural product-like NRP libraries.^15^ Type S NRPSs can thus be created much faster and in unprecedented numbers compared to conventional bio-engineering approaches (Fig. S1). Previously, we demonstrated the vast biocombinatorial potential of type S NRPSs by creating bi- and tri-partite NRPS libraries, wherein each library member consists of two or three type S subunits, respectively, which are post-translationally assembled to full length *in vivo* via the interaction of SZs, bio-synthesizing about 50 NRPs, NRP derivatives, and new to nature artificial NRPs.^15^ In this study, compared to our original proof of concept study on type S NRPSs, wherein we functionally introduced SZs within C-A linker regions at the previously defined splicing position in between individual XUs (A-T-C tri-domain units), we further broadened the applicability of SZs by successfully introducing them in between A-T and T-C interdomain linker regions.^15^ However, one bottleneck, particularly for type S NRPSs with attached SZs within the C-A linker region and type S NRPSs with certain SZ pairs (*i.e*., SZ1:2 in tri-partite type S NRPSs), was the significant drop in production yield observed.^15, 16^ Herein, we will describe the iterative optimization process (Fig. 1) of SZs and type S assembly lines, respectively, yielding up to >50-fold increased titers compared to their non-optimized equivalents. To achieve the desired optimization of NRP biosynthesis, we applied two strategies: (I) Truncation of SZs from the *N*- and/or *C*-terminus (Fig. 1, I); and (II) introduction of structurally flexible GS (Glycin-Serine) linkers into type S subunits – in between the NRPS protein subunit and the attached SZ(s) (Fig. 1, II).

**Figure 1.**
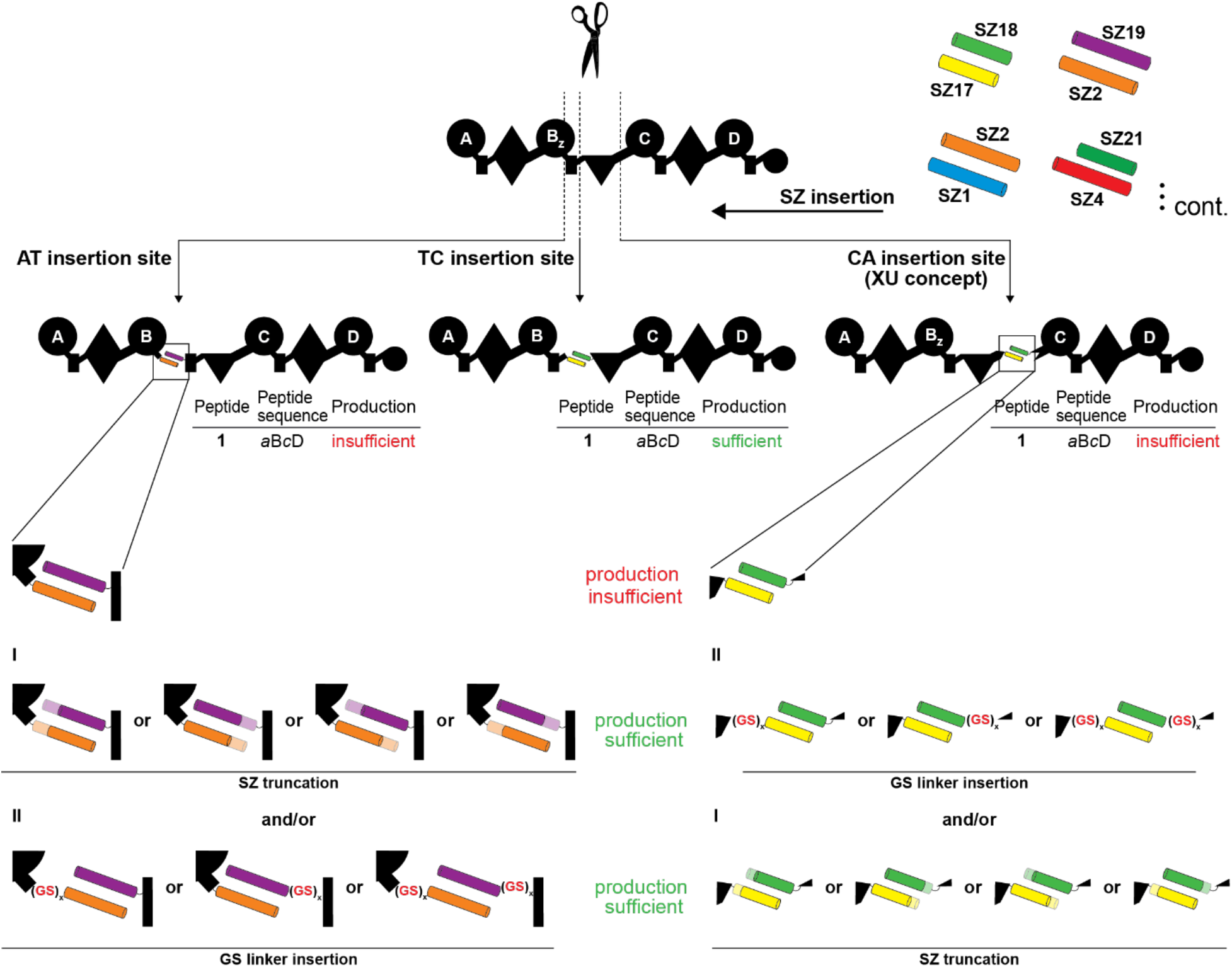
Optimization of type S NRPSs. From a pool of 27 biophysically characterized pairs,^23^ SZs can be introduced in three different positions within the A-T, T-C, and C-A linker regions, respectively. For type S NRPSs yielding insufficient production titers, we explored two optimization strategies: (I) the truncation of SZs either from the *C*-terminal, *N*-terminal, or both sides; (2) the insertion of flexible GS linkers upstream of the SZ donor, downstream of the SZ acceptor or into both positions.

### I. Insertion of GS Linker Sequences

Previously, we introduced the antiparallel SZ pair 17 & 18 (SZ17:18) at three different positions within the A-T, T-C, and C-A linker regions (the exact introduction sites are highlighted in Figure S2) of the xenotetrapeptide-producing synthetase (XtpS) from *Xenorhabdus nematophila* ATCC 19061,^24^ to create three different two protein type S XtpS variants.^15^ Although all variants showed catalytic activity, synthesizing the expected cyclic xenotetrapeptide (**1**, cyclo(*v*L*v*V); D-AA with small letters and in italics throughout the paper), we also found that the type S XtpS split within the C-A linker region (*c.f*., NRPS-1, Figure 2), resulted in the lowest production of **1** with ~30% compared to WT level (Figure 2a). In contrast, both type S XtpS splits within the A-T and T-C linker regions produced **1** at ~86% compared to WT XtpS level (Figure S2).^24^ Of note, throughout the present work, mentioned recombinant type S and WT NRPSs were produced heterologously in *E. coli* DH10B::*mtaA*. The resulting peptides (Table S1) and yields were confirmed by HPLC-MS/MS and comparison of retention times with synthetic standards (c.f., Supporting Information Figures S3-24).

**Figure 2.**
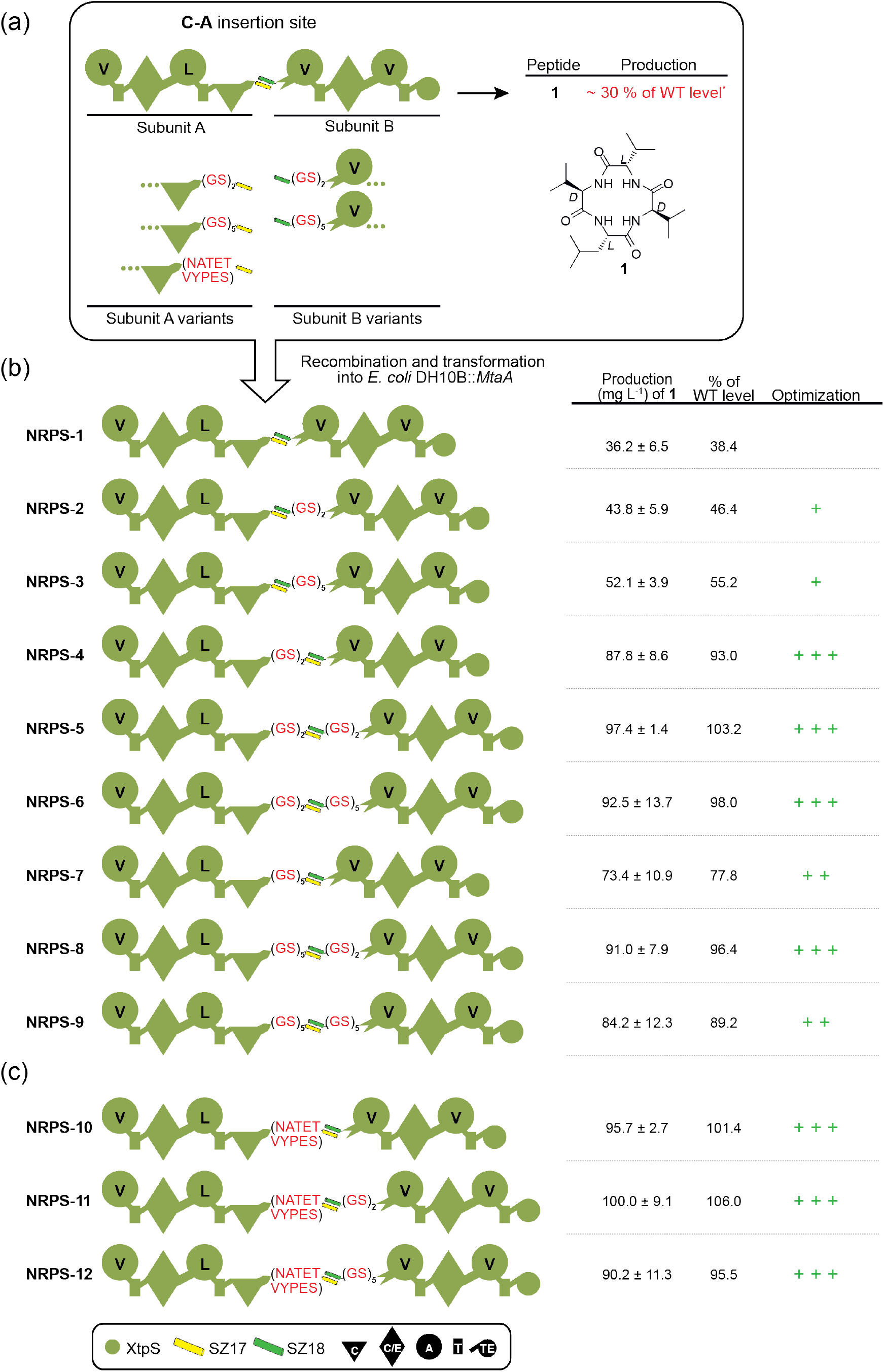
Insertion of synthetic GS-stretches of varying length into a type S XtpS system. (a) On average, the production of non-optimized NRPS-1 is at ~30% of WT level having a full-length XtpS. A set of modified subunit A and B variants were constructed by inserting GS-stretches of 4-10 AAs between the *C*-terminus of XtpS subunit A and SZ17 and SZ18 and the *N*-terminus of subunit B. (b) Generated modified subunits were re-combined with non-modified subunits and transformed into *E. coli* DH10B::*mtaA* to obtain NRPS-2 to −9. (c) The native 10 AA (NATETVYPES) were additionally inserted between the *C*-terminus of subunit A and SZ17. Production titres of NRPS-1 to −12 were compared with each other and rated +, ++, +++. Corresponding peptide yields (mg/L) and standard deviations are obtained from biological triplicate experiments. For domain assignment, the following symbols are used: (A, large circles), (T, rectangle), (C, triangle), (C/E, diamond), (TE, small circle); substrate specificities are assigned for all A domains and indicated by capital letters.

In our endeavor to further optimize type S assembly lines, we concluded from these initial results, along with available structural data^25^ of the C-A domain-domain interface and linker regions, that the introduction of SZs, due to their sheer size and rigidity could hinder the necessary structural rearrangements and thus catalytically ideal positioning of involved C, A, and T domains during the NRPS catalytic cycle. We hypothesized that more spatial flexibility would enhance dynamic domain-domain interactions of the type S XtpS (NRPS-1) variant. Hence, we introduced synthetic Gly-Ser (GS) segments of 4 and10 AAs in length between the *C*-terminus of NRPS-1 subunit A (subA_GS0) and SZ17 (subA_GS2, subA_GS5) as well as SZ18 and the *N*-terminus of NRPS-1 subunit B (sub_GS0, sub_GS2, sub_GS5), respectively (Fig. 2a).

To test these modifications, we co-transformed, produced, and analyzed all possible type S subA:subB combinations (NRPS-2 to −9) and compared peptide yields of **1** with the previously created unmodified type S NRPS-1 (subA_GS0:subB_GS0) and WT XtpS. In brief, all modified type S NRPSs (NRPS-2 to −9) showed increased catalytic activity compared to NRPS-1 (38 mg L^-1^), producing **1** at titers of 44 – 100 mg L^-1^. Interestingly, while insertions of GS2 (NRPS-2; 46 mg L-1) and GS5 (NRPS-3; 55 mg L^-1^) stretches downstream of SZ18 (subB_GS2 & sub_GS5), respectively, led to the least increase in peptide production, any modification upstream of SZ17 (subA_GS2, subA_GS5) significantly raised production titers of **1** close to (NRPS-7; 78 mg L^-1^) or back to (NRPS-4; 88 mg L^-1^) WT level.

Additionally, to run a control experiment, we reinserted the original 10 AAs (NATETVYPES) of the C-A linker that we removed in our original feasibility study^15^ to maintain the native spacing of the C and A domains involved. As we identified the position upstream of SZ17 as structurally more suitable, we chose this position to reinsert the original AAs into NRPS-1 subunit A (subA_NATETVYPES), shifting the original XU fusion site by 10 AAs (Fig. S2). From the three additionally generated synthetases harboring subA_NATETVYPES (NRPS-10 to −12, Fig 2c), NRPS-11 was the best-producing synthetase resulting from a combination of subA_NATETVYPES with subB_GS2 (100 mg L-1), biosynthesizing **1** at 106% compared to WT XtpS and 278% compared to NRPS-1.

Further modified type S XtpS with GS2 and GS5 linker insertions are shown in SI Fig. S25 (NRPS-49 to −56). Here, GS stretches were introduced into a tri-partite type S XtpS system (also depicted in, Fig. 4, NRPS-16) partitioned in between the A2-T2 and A3-T3 linker regions by introducing SZ17:18 and SZ1:2 pairs, respectively (SI Fig. S2, NRPS-16) – with emphasis on optimizing the latter SZ pair facilitating the specific interaction of the tripartite NRPS’s subunits B and C. Interestingly, again all created type S assembly lines (NRPS-49 to −56) showed catalytic activity with yields ranging from 30 to 52 mg L^-1^ and all but two (NRPS-51 & −49) showed slightly increased amounts of **1** compared to unmodified type S NRPS-16, but still only at 25 % compared to its bipartite counterpart (Fig. 3, NRPS-13). From these insights, we concluded that impairments caused by introducing SZ1:2 cannot be overcome by simply introducing flexible GS stretches, making a different optimization strategy necessary, which will be discussed in the following sections.

**Figure 3.**
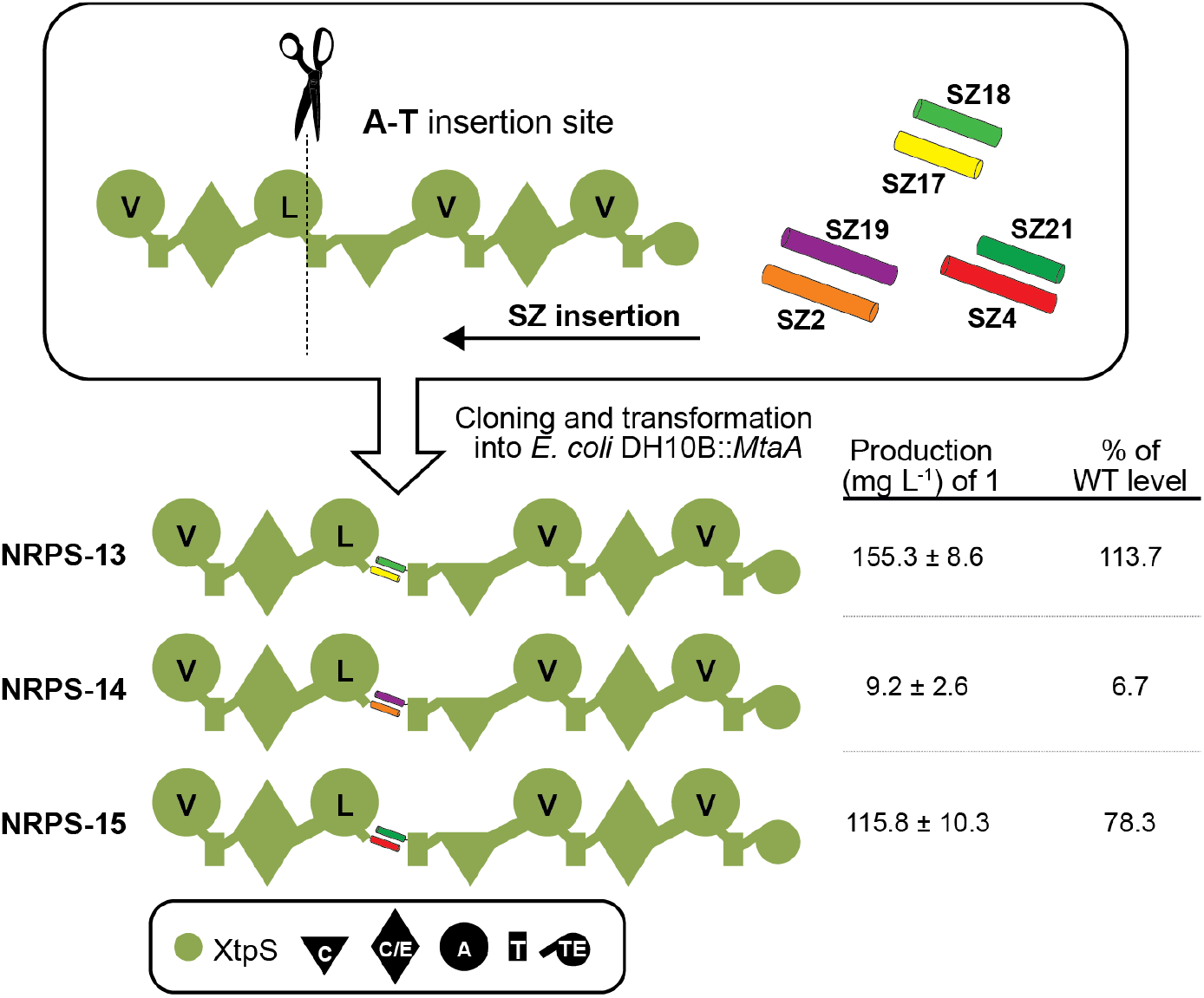
Introduction of other SZ pairs into XtpS. Two parallel interacting SZs, SZ2:19 and SZ4:21, were introduced into the A-T position of module two to generate NRPS-14 and NRPS-15. Corresponding peptide yields (mg/L) and standard deviations are obtained from biological triplicate experiments. Domain assignment is as described before.

**Figure 4.**
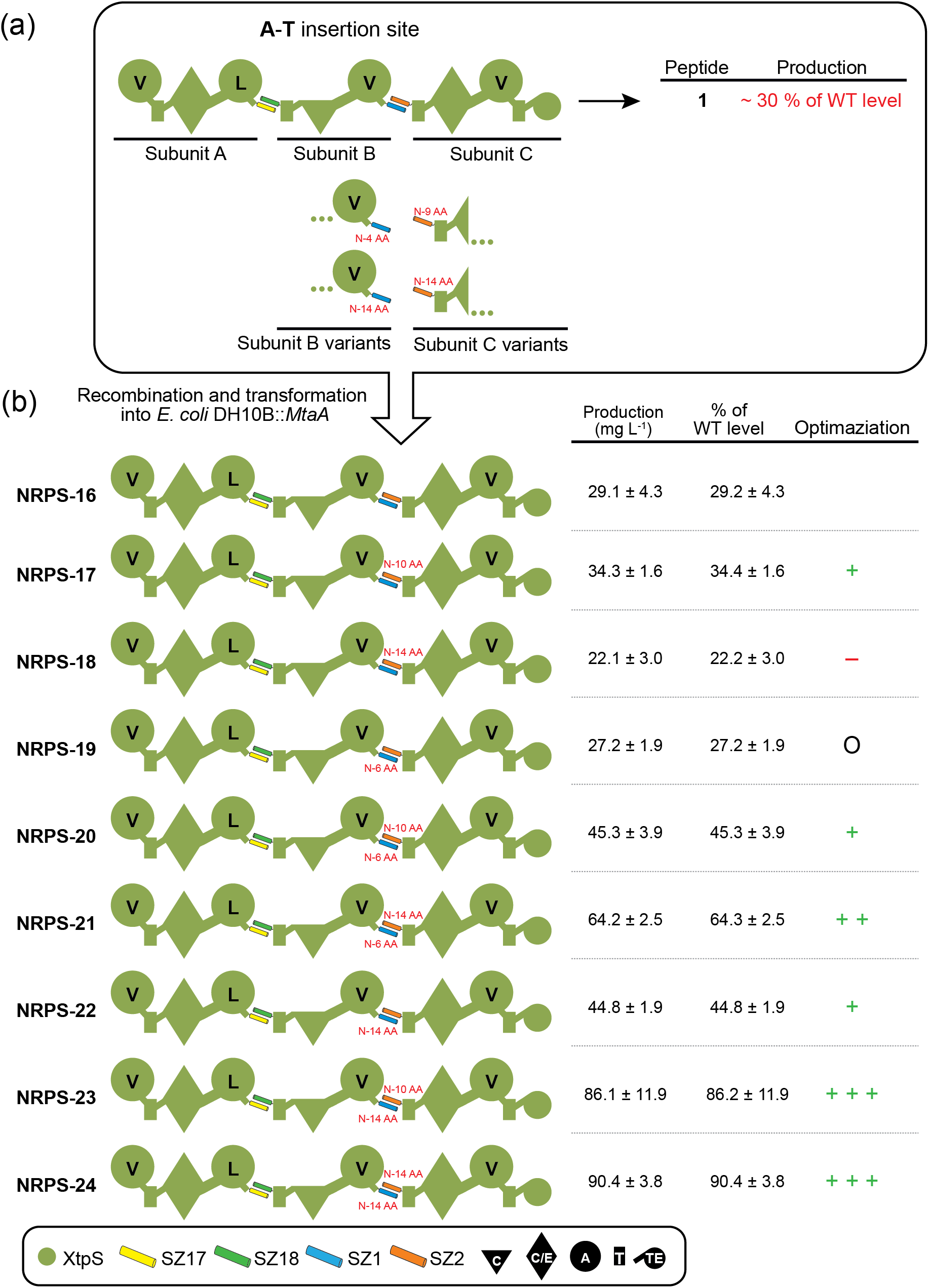
Truncation of SZ1:2 in the tri-partite XtpS. The production of non-optimized NRPS-16 is on average ~30% of WT level of a single-protein XtpS variant. A set of modified subunit B and C variants were constructed by *N*-terminally truncating SZ1 by 6 AAs and 14 AAs and SZ2 by 10 AAs and 14 AAs. Generated modified subunits were re-combined with non-modified subunits and transformed into *E. coli* DH10B::*mtaA* to obtain NRPS-16 to −24. Corresponding peptide yields (mg/L) and standard deviations are obtained from biological triplicate experiments. Production titres of NRPS-16 to −24 were compared with each other and rated -, O, +, ++, +++.

### II. Introduction of other SZ pairs

Depending on the experimental approach, it might be necessary to introduce additional or other SZ pairs than the previously applied pairs SZ17:18 and SZ1:2. Based on the observed mixed effects of the previously used SZ pairs on the productivity of type S NRPSs,^15, 16^ we decided to evaluate the effects of other SZ pairs on the functionality and productivity of NRPSs. Again targeting XtpS, we introduced two additional parallel interacting SZ pairs, SZ2:19 (NRPS-14) and SZ21:4 (NRPS-15), at the A-T position within module two (Fig. 3).

While both NRPS were functional, NRPS-14 (10 mg L^-1^) and −15 (115 mg L^-1^) resulted in reduced biosynthesis of **1** compared to WT XtpS (136 mg L^-1^) and NRPS-13 (155 mg L^-1^) (Fig. 3), the impairing effects of SZ2:19 appeared to be significantly greater. Since the experimental setup for NRPS-13 to −15 was identical except for the selected SZ pair, it is obvious that the respective selected SZ pairs are responsible for the observed effects on peptide yields. We, therefore, raised the question: What does the catalytic activity depend on, and which biophysical parameters of SZs have the most significant influence on the productivity of type S NRPS-13 to −15?

In search of an answer, we took a closer look at the biophysical data of all SZs used, compiled in a specification sheet provided by Thompson et al.^23^ We found that all three SZ pairs have similar properties, e.g., similar affinities of <10 nm and non-toxicity to the living cell (demonstrated by yeast-two-hybrid experiments with two different reporter genes), but considerably differ in length.^23^ Here, SZ17:18, used in the best-producing NRPS-13, harbors the shortest SZ pair with a length of 42 AAs (SZ17) and 41 AAs (SZ18), respectively. All other SZs available and tested so far are significantly longer, ranging from 45 AAs to 54 AAs. These insights, combined with findings of our previous work,^15, 16^, and the results of GS linker modifications of SZ1:2 (Fig. S25), we inferred a direct correlation between the length of SZs and the productivity of type S NRPSs. Consequently, we next planned to optimize chimeric type S NRPSs by truncating applied SZs.

### III. Truncation of SZs

We previously demonstrated the possibility of generating tri-partite type S assembly lines assembled from three independent type S subunits.^15^ Although type S tri-partite NRPSs further enhance the advantages associated with the insertion of SZs, i.e., increased biocombinatorial potential, they have so far suffered from low production titers (~30% of WT level, c.f. NRPS-16 Fig. 4) compared to their WT and di-partite counterparts (~86% of WT level, c.f. Fig. S2).^15^ Thus, to optimize these systems we again targeted the SZ pair 1:2 of NRPS-16 (Fig. 4) – which was already target of our GS linker insertion strategy but resulted in moderate optimization results (Fig. S25).

As a rationale to guide our optimization attempt, we took the results from Reinke et al., who investigated the truncation of SZs (exemplified for SZ4) and its effect on their stability.^22^ They found that the *N*-terminal truncation of SZ4 did not affect its stability, while the *C*-terminal truncation noticeably destabilized SZ4. Consequently, we decided to truncate SZ1 (−6 and −14 AAs) *N*-terminally and SZ2 (−10 and −14 AAs) attached to subunit B and subunit C, respectively, of NRPS-16. Co-production of all modified and unmodified subunit B and C variants together with unmodified subunit A resulted in NRPS-17 to NRPS-24 (Fig. 4). Of note, whereas the SZ pair SZ1_-6_AAs and SZ2_-10_AAs were created to simulate the length of SZ17:18, the SZ pair SZ1_-14_AAs and SZ2_-14_AAa was created because for SZ4 the two most *N*-terminal heptads were described as dispensable.

All resulting assembly lines showed catalytic activity biosynthesizing **1** in a range of 22 – 90 mg L^-1^. Of these, all but NRPS-18 and −19 resulted in a 17% to 210% increase in **1** compared to NRPS-16, with NRPS-23 (86 mg L^-1^) and −24 (90 mg L^-1^) showing the highest titers almost restoring WT XtpS production, highlighting the great potential of truncating SZs for type S NRPS optimization. However, to test the effects of further, more invasive truncations, we also attempted to remove 28 AAs from the *N*-terminus of SZ1 and 2 but found that the synthesis of **1** decreased to 62% compared to WT XtpS (SI Fig. S3, NPRS-57), suggesting that the ideal truncation is probably in the range of 14 AAs. For more truncations, see SI Fig. S27, S28, and S29. A comparative overview of all truncated SZs and their impact on peptide synthesis is shown in SI Fig. S30. In brief, truncation of SZ2:19 in NRPS-14 resulted in an increased production of four constructs (NRPS-58, −61, −63, and −64) with NRPS-64 even restoring the synthesis of **1** to WT levels (Fig. S27; NRPS-58 to −65). Truncation of SZ 17:18 at the C-A position of XtpS (NRPS-1), however, resulted in strongly decreased yields of **1** (3 – 25 mg L^-1^; Fig. S5: NRPS-66 to −80), indicating that truncation of SZ17:18 is not recommendable.

Lastly, we applied the identified best-performing SZ1:2 variant to our previously published tripartite SZ library to determine whether the observed product yield-increasing changes are exclusively linked to Xtps type S assembly line variants or whether there is an observable generality in this approach. To this end, we modified all functional (12 out of 18) type S NRPSs subunit B and subunit C variants by removing 14 AAs from the *N*-terminal sites of SZ1 and SZ2, respectively. As shown in Fig. 5, all optimized type S NRPSs resulted in a strong increase in production with an optimization between 2-to 56-fold, ranging from 123.2% to 5592.4% compared to the non-optimized constructs. These results not only confirm the apparent correlation between the length of SZs and the productivity of type S NRPSs, but also infer the general applicability of this particular optimized SZ pair for the generation of high-yield artificial type S assembly lines rather than being exclusively tied to a particular NRPS. In conclusion, truncating the SZs can completely eliminate their adverse effects, allowing SZs to be used without any restrictions for the generation of reprogrammed NRPSs.

**Figure 5.**
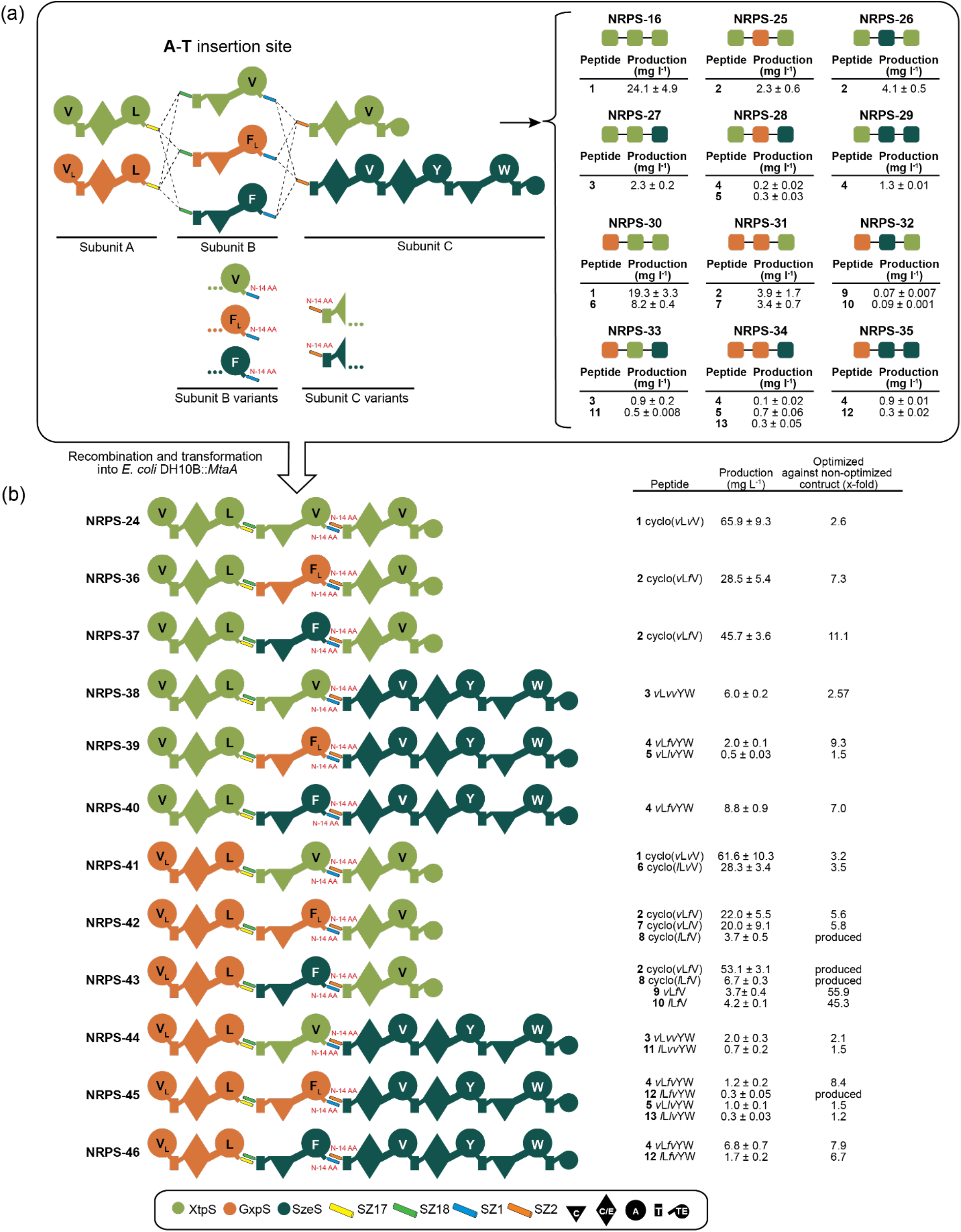
Optimized Tri-partite NRPS library. Production titres of non-optimized constructs NRPS-16 and NRPS-36 to-46 are shown on the upper right corner. Subunits B and C were modified by removing 14 AAs from the N-terminal site of SZ1 and SZ2. Generated modified subunits were re-combined with non-modified subunits A and transformed into *E. coli* DH10B::*mtaA* to obtain NRPS-24 to −46. Production titres of NRPS-24 to −46 were compared with non-optimized NRPS-15 to −35. Corresponding peptide yields (mg/L) and standard deviations are obtained from biological triplicate experiments. Domain assignment is as described before.

### IV. Other SZ networks

As SZs have several interaction partners providing access to distinct interaction networks,^22, 23^ we tried to establish the SZ mediated *in vivo* assembly of type S NRPSs beyond tri-partite assembly lines by establishing a ring network. Initially, when we created the tri-partite library (Fig. 5), we decided to choose the orthogonal network in which applied SZs 17:18 and 1:2 cannot communicate with each other. To demonstrate the applicability of other SZ networks for NRPS engineering, we recreated a previously constructed type S NRPS^15^ (NRPS-47), assembled from building blocks of XtpS and the GameXPeptide synthetase (GxpS) from *Photorhabdus luminescens* TT01^26^, by replacing SZ1:2 by SZ17:18 to generate NRPS-48 (Fig. 6). With two SZ17:18 pairs, cross-talk between both type S NRPSs should be possible, theoretically leading to none or multiple incorporations of subunit B in NRPS-48, mimicking an iterative-like NRPS system as in the rhabdopeptide biosynthesis^27^. In general, naturally occurring iterative NRPSs reuse certain modules or even entire assembly lines, resulting in peptide products that differ in composition from the number (and order) of modules in an assembly line.^9^ HPLC-MS analysis of extracts from NRPS-48 producing cultures indeed suggested none or multiple, up to three times, use of subunit B (NRPS-48a to −48d), resulting in the production of peptides **15** - **18,** which are not synthesized by NRPS-47 (Fig. 6). Additionally, with these results we were able to demonstrate that we can build not only functional di- or tri-partite type S NRPSs, but also functional penta-partite systems that theoretically allow for almost inconceivably large combinatorics. With 16 building blocks per subunit already more than a million new NRP combinations can be generated. With just a few more building blocks, this number can be driven exponentially into previously unimaginable new dimensions.

**Figure 6.**
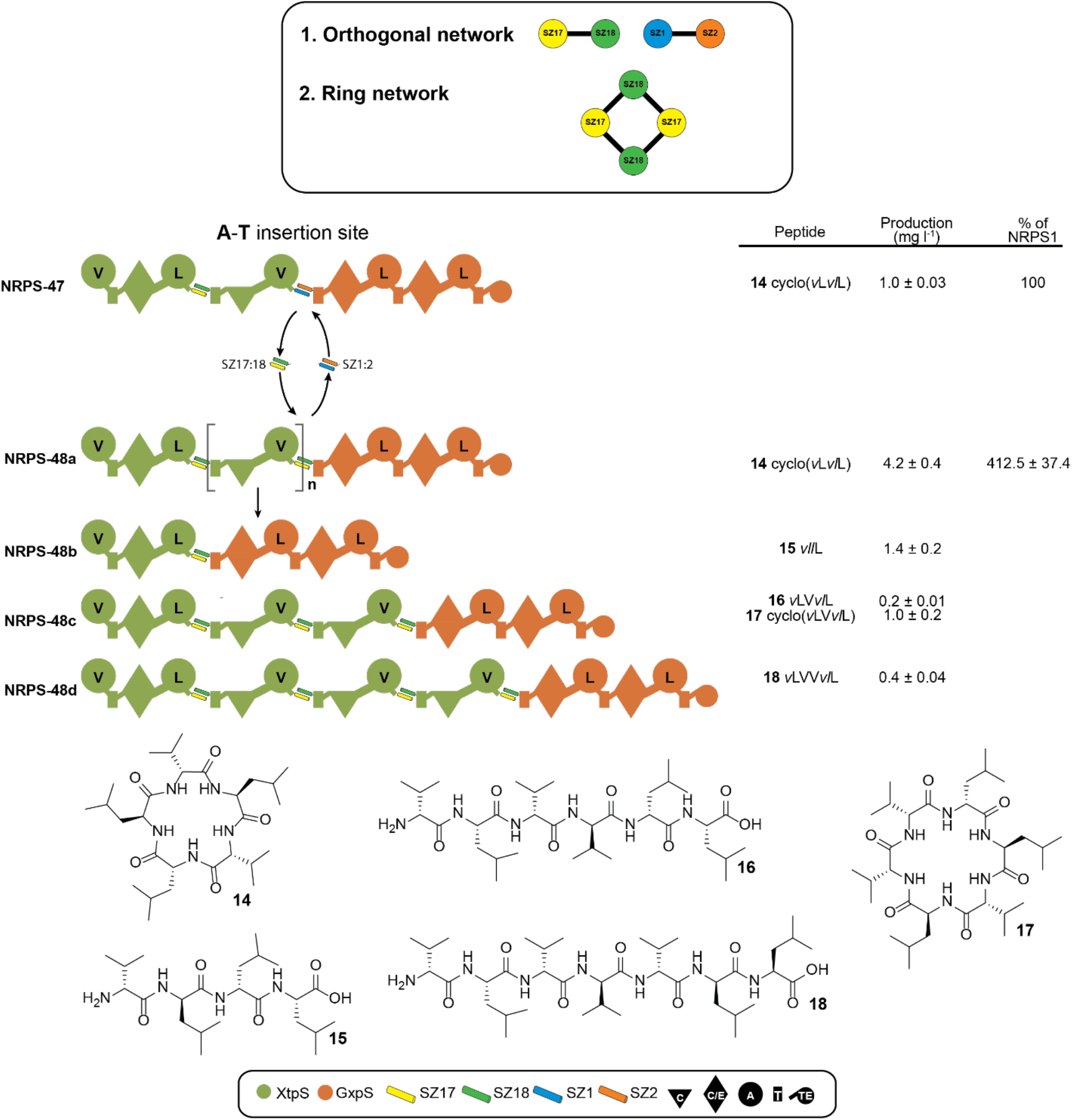
Introduction of a ring network. (A) Applied orthogonal and ring networks. SZ17:18 and SZ1:2 form an orthogonal network, meaning both SZs do not interact with each other. Introducing two SZ17:18 pairs result in a ring network whereby applied SZs cross-talk. (B) For the construction of a ring interaction network, SZ1:2 in NRPS-47 was changed against SZ17:18, resulting in NRPS-48. NRPS-48 was capable to incorporate subunit B not at all or up to three times (NRPS-18a to 18c), leading to the production of peptides **15** - **18**. Corresponding peptide yields (mg/L) and standard deviations are obtained from biological triplicate experiments. Domain assignment is as described before and chemical structure of produced peptides **15** - **18** shown at the bottom.

## Conclusion

SynBio – the design and engineering of biological systems to create and improve processes and products – is becoming a disruptive force transforming the economy into a bio-based Bio-Economy. Biology is usually defined as the study of living things and life itself, but SynBio has turned science into the manufacturing paradigm of the future. Genetic engineering can, in theory, turn microorganisms into bio-factories producing almost everything that human beings consume, from fuels, flavors, fabrics to foods and drugs. But besides all the future potential of SynBio, most innovations are still in their infancy. Especially in the field of natural products research, there are only a few examples of successfully applied SynBio for the development of novel drugs or their manufacturing, such as SynBio’s first malaria drug Artemisinin.^28^

Large-scale bioengineering of NRPSs using SZs is thus a promising strategy for obtaining a variety of new valuable natural products, but as with many new technologies, there are limitations, namely low yields for some type S NRPS constructs. To overcome this bottleneck turning type S NRPSs into a valuable tool for the production of NRPs with high yields and the development of novel bioactive molecular scaffolds with high confidence, we pursued two strategies: firstly, the insertion of flexible and unstructured GS stretches (Fig. 2); and secondly, the targeted truncation of available SZ pairs (Fig. 4 and 5). With those approaches, we aimed to reduce the presumably introduced rigidity of type S NRPSs – probably caused by the insertion of the structurally stable alpha-helical SZs that can have a size of up to 54 AAs – while simultaneously enhancing the highly dynamic domain-domain interactions of NRPSs during the catalytic biosynthesis cycle. In particular, the possibility of introducing a plethora of distinct SZ pairs into three interdomain linker regions increases the likelihood of creating impaired type S NRPSs, making efficient, rational optimization strategies necessary. With the iterative optimization strategy outlined in this paper, we have not only presented two extremely efficient SZ pairs (SZ17:18 & SZ1:2) for the generation of di- and tri-partite type S NRPSs, but also paved the way for the rapid optimization of other SZ pairs of interest.

Currently, SZ 17:18 with 42 and 41 AAs not only it is the shortest readily available pair but also the most efficient to generate unimpaired high-yielding type S NRPSs that even outcompete WT NRPSs (c.f., NRPS-13, Fig. 3). These extraordinary capabilities of SZ17:18, if introduced correctly (c.f. NRPS-1 vs. NRPS-10, Fig. 2) also appeared in our proof of concept study, in which we compared covalently fused recombinant NRPSs with analogues SZ linked type S variants to examine the impact of SZ17:18. Even with the introduction of the respective unoptimized SZ pair (c.f., Fig. 2 NRPS-1) into recombinant NRPSs, observed peptide yields did not decrease compared to the recombinant covalent counterparts. Moreover, truncating SZ17:18 in NRPS-1 (Fig. S5) led to drastically reduced production of **1**, indicating its ideally suited biophysical character for NRPS bio-engineering purposes. We therefore recommend using the length of SZ17:18 as a guide for the optimization of other SZ pairs.

Nevertheless, to optimize further SZ pairs, the ideal length, and composition should still be determined experimentally for every unique SZ pair analogous to the workflow presented here. Noteworthy, in prior work, we observed decreasing peptide production to ~30% compared to WT levels at the C-A position upon the insertion of SZ17:18 in XtpS (NRPS-1), which, however, was not due to the length of SZ 17:18 but rather caused by the deletion of the native 10 AAs of the particular C-A linker region. This AAs stretch has been deleted because we assumed that maintaining the native distance of the C- and A-domains is essential. Apparently, we underestimated the structural flexibility of NRPSs and noticed that once the native AAs were reinserted, WT-level peptide production could be restored (c.f. NRPS-10, Fig. 2). Therefore, we would like to revise our initial design and recommend to keep the native C-A linker AAs in type S NRPSs and choose the fusion site for SZ insertion as depicted in Fig. S2.

In contrast to SZ17:18, unmodified SZ1:2 is significantly longer, resulting in reduced production of peptides in several constructs. The detrimental impact of SZ1:2 is also apparent in NRPS-17 (Fig. 4). Replacing SZ1:2 with SZ17:18 in NRPS-18 resulted in a four-fold increase in production. Truncating SZ1:2 by 14 AAs restored production to WT levels (NRPS-24) and increased the productivity ~100-fold in SZ1:2 optimized chimeric type S NRPS (NRPS-24 to −46) compared to non-optimized assembly lines (NRPS-16).

In case the truncation of any other inserted SZ pair does not lead to an increased or restored peptide production, or if the truncated SZs lose their stability and thus affinity, we strongly recommend to use GS stretches to increase the enzyme’s spatial flexibility (Fig. 1). The reduced productivity does not exclusively depend on the length of applied SZs but also on the insertion point itself and the targeted NRPS system, which might not provide enough spatial flexibility to allow the efficient progressing of the catalytic cycle.

Last but not least, we were capable to draw most interesting conclusions from the applied ring network. Besides bi- and tri-partite systems, we could also generate tetra- and penta-partite systems, raising theoretical (bio-)combinatorial possibilities to yet unprecedented dimensions.

