## Supplementary Information for "Guidelines for Optimizing Type S Non-Ribosomal Peptide Synthetases"

* Corresponding author

### 1. Material and Methods

#### 1.1 Cultivation of Strains

Cultivation was done as described before: All *E. coli*, *Photorhabdus* and *Xenorhabdus* strains were cultivated in LB (10 g/L Trypton, 5 g/L yeast extract, 10 g/L NaCl, pH 7,5) or TB liquid medium (12 g/L Tryton, 24 g/L yeast extract, 0.4% (v/v) Glycerin, 10% (v/v), 17 mM $KH_2PO_4$, 72 mM $K_2HPO_4$, pH 6.5) at 37°C (*E. coli*) or 30°C (*Photorhabdus*, *Xenorhabdus*) for 16-18 h at 160-200 rpm. 1% (w/v) agar was added for growth on solid LB. If necessary, medium was supplemented 1:1000 with kanamycin- (50 µg/mL in sterile $ddH_2O$), chloramphenicol (34 µg/mL in ethanol) and/or spectinomycin stock solution (50 mg/mL in sterile $ddH_2O$). For short-time storage LB agar plates were stored either at 4°C (*E. coli*) or 18°C (*Photorhabdus*, *Xenorhabdus*). For permanent storage, liquid cultures were supplemented with 20% (v/v) glycerol and frozen at -80°C.[1]

#### 1.2 Plasmid assembly

Genomic DNA from *Xenorhabdus* and *Photorhabdus* were isolated using the Gentra Puregene Yeast/Bact Kit (Qiagen) accordingly to the manufacturers' instruction for Gram negative bacteria. Plasmid DNA was isolated using PureYield Plasmid Miniprep System (Promega). PCRs were performed with oligonucleotides obtained from Eurofins Genomics (Table S4) containing homology arms of ~20 bp in a one or two step PCR program. Phusion Hot Start Flex (New England Biolabs) was applied as High Fidelity DNA Polymerase and used accordingly to the manufacturers' instruction. PCR fragments were digested with DpnI (Thermo Fisher Scientific). Purification of all fragments was performed with Monarch PCR & DNA Cleanup Kit or from 1% TAE agarose gel using Monarch Gel Extraction Kit. Plasmid assembly was done by HiFi (New

England Biolabs) or Hot Fusion cloning and DNA mix was transformed into *E. coli* DH10B via electroporation. Cells were regenerated in LB for 1 h at 37°C and plated on LB agar plates containing appropriate antibiotics. Plasmids were isolated an verified by plasmid digest and DNA sequencing using sanger sequencing (Eurofins Genomics).[1]

#### 1.3 Heterologous expression of NRPS templates and HPLC-MS analysis

Constructed plasmids were transformed into *E. coli DH10B*::mtaA, and cells from one colony were grown overnight in LB medium containing all necessary antibiotics (50 µg/ml kanamycin, 34 µg/ml chloramphenicol, 50 µg/ml spectinomycin). 100 µl of the overnight culture were used to inoculate 10 ml LB medium containing antibiotics, 0.002 mg/mL L-arabinose and 2 % (v/v) XAD-16. After 72 h at 22 °C, XAD-16 beads were harvested and incubated with one culture volume methanol. for 60 min at 180 rpm. The organic phase was filtrated and extracts were evaporated to dryness. With 1 mL MeOH, extracts were re-solved, centrifuged for 20 min and diluted 1:10 for HPLC-MS analysis. Liquid chromatography was performed on an UltiMate 3000 LC system (Dionex) with an installed C18 column (ACQUITY UPLCTM BEH, 130 Å, 2.1 mm x 100 mm, Waters). Separation was conducted at a flow rate of 0.4 mL/min using acetonitrile (ANC) and water containing 0.1% formic acid (v/v) in a 5-95% gradient over 16 min. Mass spectrometric analyses were performed using an ESI ion-trap mass spectrometer (AmaZon X, Bruker) or ESI. ESI-MS spectra were recorded in positive-ion-mode with the mass range from 100-1200 *m/z* and ultraviolet (UV) at 200-600 nm. Evaluation was performed using DataAnalysis 4.3 software (Bruker).[1]

#### 1.4 Peptide quantification

Absolute production titers were calculated as previously described.[2] Synthetic standard **1** (for the quantification of **1**, **2**, **6**, **7** and **8**), **3** (for the quantification of **4**, **5**, **9**, **10** and **11**) and **12** (for the quantification of **12**) were obtained from Synpeptide. Synthetic standard **15**, **16**, **17** and **18** were synthesized as described below.

#### 1.5 Chemical synthesis

Peptide synthesis was done automatically with the Syro Wave™ peptide synthesizer (Biotage, Sweden) using standard Fmoc/t-Bu chemistry on a 25 or 50 µM scale. Fmoc amine-protected AAs in dimethylformamide (DMF) was added to preloaded H-AAn-2-CT resin and the coupling reaction was perfomred by adding HCTU (O-(6-chlorobenzotriazol-1-yl)-N,N,N′,N′-

tetramethyluronium hexafluorophosphate) in DMF (25 µmol: 250 µL, 0.54 mol/L, 5.4 eq.; 50 µmol: 500 µL, 0.27 mol/L, 2.7 eq.) and DIPEA (N,N-diisopropylethylamine) in NMP for 50 min alternating between shaking (15 s) and pausing (2 min). Washing the resin with 800 µL NMP was followed by adding the capping solution (0.45 mL DIPEA, 0.95 mL $Ac_2O$, 40 mg HOBt auf 20 mL NMP; 25 µmol: 500 µL; 50 µmol: 1000 µL) and incubating for 5 min (15 s shaking, 1 min pausing). Fmoc protecting group was cleaved off by incubation with 40% piperidine in NMP (25 µmol: 300 µL; 50 µmol: 600 µL) for 3 min (shaking 10 s and pausing 1 min) and 20% piperidine in NMP for 10-min (shaking 10 s and pausing 2 min). Between each reaction step, resin was washed with 800 µL NMP. After synthesis, the resin was washed 5 times each with NMP, DMF and DCM and dried.

The peptide was cleaved off from the solid phase by adding the cleavage cocktail (1:4 HFIP (hexafluoroisopropanol)/DCM) for one hour and rinsed twice with the cleavage cocktail afterwards. The resin was removed by filtration and the cleavage cocktail was evaporated. For intramolecular cyclization, the peptide was dissolved in DMF/DCM (25 µmol, 25 mL, 1 mM) and mixed with HATU (38 mg, 100 µmol, 4 eq.) and DIPEA (13 mg, 17 µL, 100 µmol, 4 eq.) followed by incubating for 20 min at 60 °C. The cyclized or linear peptide was dissolved in DMSO, DMF and MeOH and purified by preparative HPLC (Pure chromatography system, Büchi). The purity was determined by HPLC-MS.

### 2. Supplementary tables

**Table S1.** ESI-MS data of all produced peptides.

| Peptide (#) | mass-to-charge ratio (*m/z*) | Molecular formula | AA sequence | Reference |
|---|---|---|---|---|
| **1** | 411.29 | $C_{21}H_{38}O_4N_4$ | cyclo(*v*L*v*V) | [3] |
| **2** | 459,30 | $C_{25}H_{38}N_4O_4$ | cylco(*v*L*f*V) | [1] |
| **3** | 778,45 | $C_{41}H_{59}N_7O_8$ | *v*L*vv*YW | [1] |
| **4** | 826.45 | $C_{45}H_{59}N_7O_8$ | *v*L*fv*YW | [1] |
| **5** | 792.47 | $C_{42}H_{61}N_7O_8$ | *v*L*lv*YW | [1] |
| **6** | 425.31 | $C_{22}H_{40}N_4O_4$ | cyclo(*l*L*v*V) | [1] |
| **7** | 425,31 | $C_{22}H_{40}N_4O_4$ | cyclo(*v*L*l*V) | [1] |
| **8** | 472.31 | $C_{26}H_{40}N_4O_4$ | cyclo(*l*L*f*V) | [1] |
| **9** | 476.62 | $C_{25}H_{40}N_4O_5$ | *v*L*f*V | [1] |
| **10** | 490.65 | $C_{26}H_{42}N_4O_5$ | *l*L*f*V | [1] |
| **11** | 792.47 | $C_{42}H_{61}N_7O_8$ | *l*L*vv*YW | [1] |
| **12** | 840.47 | $C_{46}H_{61}N_7O_8$ | *l*L*fv*YW | [1] |
| **13** | 806.48 | $C_{43}H_{63}N_7O_8$ | *l*L*lv*YW | [1] |
| **14** | 538.40 | $C_{28}H_{51}O_5N_5$ | cyclo(*v*L*vl*L) | [1] |
| **15** | 470.35 | $C_{24}H_{46}O_5N_4$ | *vl*L | this study |
| **16** | 655.47 | $C_{33}H_{62}O_7N_6$ | *v*LV*vl*L | this study |

| | | | | |
|---|---|---|---|---|
| **17** | 637.46 | $C_{33}H_{60}O_6N_6$ | cyclo(*v*LV*vl*L) | this study |
| **18** | 754.54 | $C_{38}H_{71}O_8N_7$ | *v*LVV*vl*L | this study |

**Table S2.** Strains used in this work.

| Strain | Genotype/ NRPS | Reference |
|---|---|---|
| *E. coli* DH10B | F_mcrA (*mrr-hsd*RMS-*mcr*BC), 80*lac*ZΔ, M15, Δ*lac*X74 *rec*A1 *end*A1 *ara*D 139Δ(*ara*, *leu*)7697 *gal*U *gal*K λ *rps*L (*Strr*) *nup*G / - | [4] |
| *E. coli* DH10B::*mta*A | DH10B with *mtaA* from pCK_*mtaA*Δ*entD* / - | [5] |
| *P. luminescens* TTO1 | - / *gxpS* | DSMZ |
| *X. nematophila* ATCC 19061 | - / *xtpS* | ATCC |
| *X. szentirmaii* DSM 16338 | - / *szeS* | DSMZ |

**Table S3.** Plasmids used in this work.

| Plasmids | Genotype | Reference |
|---|---|---|
| pCOLA_ara/tacI | ori ColA, $kan^R$, *araC-$P_{BAD}$* and *tac*I | unpublished |
| pCK_0402 | ori p15A, $cm^R$, *araC-$P_{BAD}$* and *tac*I-*araE* | [6] |
| pCOLA_ara_xtpS_tacI_JW | ori ColA, $kan^R$, *araC-$P_{BAD}$ xtpS* and *tac*I | [6] |
| pCOLA_ara_gxpS_tacI_JW | ori ColA, $kan^R$, *araC-$P_{BAD}$ gxpS* and *tac*I | [6] |
| pNA2 | ori p15A, $cm^R$, *araC-$P_{BAD}$ xtpS*_ $A_1T_1C/E_2A_2T_2$-SYNZIP17 und *tac*I-*araE* | [1] |
| pNA3 | ori ColA, $kan^R$, *araC-$P_{BAD}$* SYNZIP18-*xtpS*_ $C_3A_3T_3C/E_4A_4T_4$TE und *tac*I | [1] |
| pNA4 | ori p15A, $cm^R$, *araC-$P_{BAD}$ xtpS*_ $A_1T_1C/E_2A_2$-SYNZIP17 und *tac*I-*araE* | [1] |
| pNA5 | ori ColA, $kan^R$, *araC-$P_{BAD}$* SYNZIP18-*xtpS*_ $T_2C_3A_3T_3C/E_4A_4T_4$TE und *tac*I | [1] |
| pNA8 | ori p15A, $cm^R$, *araC-$P_{BAD}$ xtpS*_$A_1T_1C/E_2A_2T_2C_3$-$(GS)_5$-SYNZIP17 and *tac*I-*araE* | this study |
| pNA9 | ori p15A, $cm^R$, *araC-$P_{BAD}$ xtpS*_$A_1T_1C/E_2A_2T_2C_3$-$(GS)_4$-SYNZIP17 and *tac*I-*araE* | this study |
| pNA10 | ori p15A, $cm^R$, *araC-$P_{BAD}$ xtpS*_$A_1T_1C/E_2A_2T_2C_3$-$(GS)_2$-SYNZIP17 and *tac*I-*araE* | this study |
| pNA15 | ori ColA, $kan^R$, *araC-$P_{BAD}$* SYNZIP18-*xtpS*_$T_2C_3A_3$-SYNZIP1 and *tac*I | [1] |
| pNA16 | ori CloDF13, $spec^R$, *araC-$P_{BAD}$* SYNZIP2-*xtpS*_$T_3C/E_4A_4T_4$TE and *tac*I | [1] |
| pNA17 | ori ColA, $kan^R$, *araC-$P_{BAD}$* SYNZIP18-*xtpS*_$C_3A_3T_4$-SYNZIP1 and *tac*I | [1] |
| pNA18 | ori CloDF13, $spec^R$, *araC-$P_{BAD}$* SYNZIP2-*xtpS*_C/$E_4A_4T_4$TE and *tac*I | [1] |

| | | |
|---|---|---|
| pNA26 | ori p15A, $cm^R$, *araC-$P_{BAD}$ gxpS*_$A_1T_1C/E_2A_2$-SYNZIP17 and *tac*I-*araE* | [1] |
| pNA27 | ori ColA, $kan^R$, *araC-$P_{BAD}$* SYNZIP18-*gxpS*_$T_2C_3A_3$-SYNZIP1 and *tac*I | [1] |
| pNA30 | ori ColA, $kan^R$, *araC-$P_{BAD}$* SYNZIP18-*szeS*_$T_2C_3A_3$-SYNZIP1 and *tac*I | [1] |
| pNA31 | ori CloDF13, $spec^R$, *araC-$P_{BAD}$* SYNZIP2-*szeS*_$T_3C/E_4A_4T_4$ $C/E_5A_5T_5C_6A_6T_6$TE and *tac*I | [1] |
| pNA40 | ori p15A, $cm^R$, *araC-$P_{BAD}$ xtpS*_ $A_1T_1C/E_2A_2$-SYNZIP2 und *tac*I-*araE* | this study |
| pNA41 | ori ColA, $kan^R$, *araC-$P_{BAD}$* SYNZIP19-*xtpS*_ $T_2C_3A_3T_3C/E_4A_4T_4$TE und *tac*I | this study |
| pNA42 | ori p15A, $cm^R$, *araC-$P_{BAD}$ xtpS*_ $A_1T_1C/E_2A_2$-SYNZIP21 und *tac*I-*araE* | this study |
| pNA43 | ori ColA, $kan^R$, *araC-$P_{BAD}$* SYNZIP4-*xtpS*_ $T_2C_3A_3T_3C/E_4A_4T_4$TE und *tac*I | this study |
| pNA72 | ori p15A, $cm^R$, *araC-$P_{BAD}$ xtpS*_ $A_1T_1C/E_2A_2$-N-terminally truncated SYNZIP2 (-9 AA) und *tac*I-*araE* | this study |
| pNA73 | ori ColA, $kan^R$, *araC-$P_{BAD}$* N-terminally truncated SYNZIP19 (-2 AA)-*xtpS*_ $T_2C_3A_3T_3C/E_4A_4T_4$TE und *tac*I | this study |
| pNA145 | ori p15A, $cm^R$, *araC-$P_{BAD}$ xtpS*_$A_1T_1C/E_2A_2T_2C_3$-N-terminally truncated SYNZIP17 (-7 AA) and *tac*I-*araE* | this study |
| pNA146 | ori ColA, $kan^R$, *araC-$P_{BAD}$* N-terminally truncated SYNZIP18 (-7 AA)-*xtpS*_$A_3T_3C/E_4A_4T_4$TE and *tac*I | this study |
| pNA147 | ori p15A, $cm^R$, *araC-$P_{BAD}$ xtpS*_$A_1T_1C/E_2A_2T_2C_3$- N-terminally truncated SYNZIP17 (-14 AA) and *tac*I-*araE* | this study |
| pNA148 | ori ColA, $kan^R$, *araC-$P_{BAD}$* N-terminally truncated SYNZIP18 (-14 AA)-*xtpS*_$A_3T_3C/E_4A_4T_4$TE and *tac*I | this study |
| pNA149 | ori ColA, $kan^R$, *araC-$P_{BAD}$* SYNZIP18-*xtpS*_$T_2C_3A_3$- N-terminally truncated SYNZIP1 (-14 AA) and *tac*I | this study |
| pNA150 | ori CloDF13, $spec^R$, *araC-$P_{BAD}$* N-terminally truncated SYNZIP2 (-14 AA)-*xtpS*_$T_3C/E_4A_4T_4$TE and *tac*I | this study |
| pNA151 | ori ColA, $kan^R$, *araC-$P_{BAD}$* N-terminally truncated SYNZIP4 (-14 AA)-*xtpS*_ $T_2C_3A_3T_3C/E_4A_4T_4$TE und *tac*I | this study |
| pNA152 | ori ColA, $kan^R$, *araC-$P_{BAD}$* SYNZIP18-*xtpS*_$T_2C_3A_3$-SYNZIP17 and *tac*I | this study |
| pNA153 | ori CloDF13, $spec^R$, *araC-$P_{BAD}$* SYNZIP18-*xtpS*_$T_3C/E_4A_4T_4$TE and *tac*I | this study |
| pNA154 | ori ColA, $kan^R$, *araC-$P_{BAD}$* SYNZIP18-$(GS)_5$-*xtpS*_$A_3T_3C/E_4A_4T_4$TE and *tac*I | this study |
| pNA155 | ori ColA, $kan^R$, *araC-$P_{BAD}$* SYNZIP18-$(GS)_2$-*xtpS*_$A_3T_3C/E_4A_4T_4$TE and *tac*I | this study |
| pNA156 | ori p15A, $cm^R$, *araC-$P_{BAD}$ xtpS*_$A_1T_1C/E_2A_2T_2C_3$-SYNZIP17-(NATETVYPES) and *tac*I-*araE* | this study |
| pNA157 | ori ColA, $kan^R$, *araC-$P_{BAD}$* SYNZIP18-*xtpS*_$T_2C_3A_3$-$(GS)_5$-SYNZIP1 and *tac*I | this study |
| pNA158 | ori ColA, $kan^R$, *araC-$P_{BAD}$* SYNZIP18-*xtpS*_$T_2C_3A_3$-$(GS)_2$-SYNZIP1 and *tac*I | this study |
| pNA159 | ori CloDF13, $spec^R$, *araC-$P_{BAD}$* SYNZIP2-$(GS)_5$-*xtpS*_$T_3C/E_4A_4T_4$TE and *tac*I | this study |

| | | |
|---|---|---|
| pNA160 | ori CloDF13, spec$^R$, *araC*-$P_{BAD}$ SYNZIP2-(GS)$_2$-*xtpS*_$T_3C/E_4A_4T_4TE$ and *tac*I | this study |
| pNA161 | ori p15A, cm$^R$, *araC*-$P_{BAD}$ *xtpS*_ $A_1T_1C/E_2A_2$-N-terminally truncated SYNZIP2 (-14 AA) und *tac*I-*araE* | this study |
| pNA162 | ori ColA, kan$^R$, *araC*-$P_{BAD}$ N-terminally truncated SYNZIP19 (-7 AA)-*xtpS*_ $T_2C_3A_3T_3C/E_4A_4T_4TE$ und *tac*I | this study |
| pNA163 | ori p15A, cm$^R$, *araC*-$P_{BAD}$ *xtpS*_$A_1T_1C/E_2A_2T_2C_3$- C-terminally truncated SYNZIP17 (-7 AA) and *tac*I-*araE* | this study |
| pNA164 | ori ColA, kan$^R$, *araC*-$P_{BAD}$ C-terminally truncated SYNZIP18 (-7 AA)-*xtpS*_$A_3T_3C/E_4A_4T_4TE$ and *tac*I | this study |
| pNA165 | ori ColA, kan$^R$, *araC*-$P_{BAD}$ SYNZIP18-*xtpS*_$T_2C_3A_3$-N-terminally truncated SYNZIP1(-28 AA) and *tac*I | this study |
| pNA166 | ori CloDF13, spec$^R$, *araC*-$P_{BAD}$ N-terminally truncated SYNZIP2 (-28 AA)-*xtpS*_$T_3C/E_4A_4T_4TE$ and *tac*I | this study |
| pNA167 | ori ColA, kan$^R$, *araC*-$P_{BAD}$ SYNZIP18-*gxpS*_$T_2C_3A_3$-N-terminally truncated SYNZIP1 (-14 AA) and *tac*I | this study |
| pNA168 | ori ColA, kan$^R$, *araC*-$P_{BAD}$ SYNZIP18-*szeS*_$T_2C_3A_3$-N-terminally truncated SYNZIP1 (-14 AA) and *tac*I | this study |
| pNA169 | ori CloDF13, spec$^R$, *araC*-$P_{BAD}$ N-terminally truncated SYNZIP2 (-14 AA)-*szeS*_$T_3C/E_4A_4T_4$ $C/E_5A_5T_5C_6A_6T_6TE$ and *tac*I | this study |
| pNA170 | ori pUC19, kan$^R$, *araC*-$P_{BAD}$ SYNZIP18-*xtpS*_$T_2C_3A_3$-SYNZIP17 and *tac*I | this study |
| pJW61 | ori p15A, cm$^R$, *araC*-$P_{BAD}$ *xtpS*_$A_1T_1C/E_2A_2T_2C_3$-SYNZIP17 and *tac*I-*araE* | [6] |
| pJW62 | ori ColA, kan$^R$, *araC*-$P_{BAD}$ SYNZIP18-*xtpS*_$A_3T_3C/E_4A_4T_4TE$ and *tac*I | [6] |

**Table S4.** Oligonucleotides used in this work.

| Plasmids | Oligo-nucleotides | Sequence (5' →3'; overlapping ends) | Template |
|---|---|---|---|
| pNA8 | KB-pACYC-II-FW | AACGAGAAGGAGGAATTAAAATCG | pJW61 |
| | na17_RV | CGATTTTAATTCCTCCTTCTCGTTTGATCCCGAACCTGAGCCGGATCCAGACCCCCAGGTTTTTAACAACAATGTGC | pJW61 |
| pNA9 | KB-pACYC-II-FW | AACGAGAAGGAGGAATTAAAATCG | pJW61 |
| | na19_RV | CGATTTTAATTCCTCCTTCTCGTTCGAACCTGAGCCGGATCCAGACCCCCAGGTTTTTAACAACAATGTGC | pJW61 |
| pNA10 | KB-pACYC-II-FW | AACGAGAAGGAGGAATTAAAATCG | pJW61 |
| | na20_RV | CGATTTTAATTCCTCCTTCTCGTTGGATCCAGACCCCCAGGTTTTTAACAACAATGTG | pJW61 |
| pNA40 | na3 | TGGGCTAACAGGAGGAATTCCATGAAAGATAGCATGGCTAAAAGGG | *X. nematophila* ATCC 19061 |
| | na87 | CTTACGCAGATACGCGTTACGCGCATAAATCTGGCGGGCGAA | *X. nematophila* ATCC 19061 |
| | na85 | GCTGGAACGTGATGAACAGAACCTGGAAAAAATCATCGCGAACCTGCGTGACGAAATCGCGCGTCTCGAAAACGAAGTTGCGTCTCACGAACAGTGACAATTAATCATCGGCTCG | pCK_0402 |

| | | | |
|---|---|---|---|
| | na43 | CATGGAATTCCTCCTGTTAGCC | pCK_0402 |
| | na86 | GCGCGTAACGCGTATCTGCGTAAGAAAATCGCACGTCTGAAAAAGACAACCTGCAGCTGGAACGTGATGAACAGAAC | pNA40_BB1_SZ2half |
| | na43 | CATGGAATTCCTCCTGTTAGCC | pNA40_BB1_SZ2half |
| pNA41 | na90 | GACGCGTACAAAAACCGTCTGGTTGCGCCACAAGGAGAA | *X. nematophila* ATCC 19061 |
| | na7 | CGAGCCGATGATTAATTGTCACAGCGCCTCCACTTCG | *X. nematophila* ATCC 19061 |
| | jw61 | TGACAATTAATCATCGGCTCG | pCOLA_ara/tacI |
| | na88 | GTTTCTGTTTCAGCTGTTCACGTTTCTGCTTCAGCTCTTCGTTACGGTTCTTCAGTTCTTCTTTTTTGTTCTCCAGAGATTCCAGTTCGTTCATGGAATTCCTCCTGTTAGCC | pCOLA_ara/tacI |
| | jw61 | TGACAATTAATCATCGGCTCG | pNA41_BB1_SZ19half |
| | na89 | CAGACGGTTTTTGTACGCGTCCAGTTTGTTACGCAGAGCCGCCAGTTTCTGTTTCAGCTGTTCACG | pNA41_BB1_SZ19half |
| pNA42 | na3 | TGGGCTAACAGGAGGAATTCCATGAAAGATAGCATGGCTAAAAAGGG | *X. nematophila* ATCC 19061 |
| | na93 | TTCCAGCTGCGCAACTTCGTTATAAATCTGGCGGGCGAA | *X. nematophila* ATCC 19061 |
| | na91 | GCGTACCTGGAGAAGGAGATCGCGCGTCTGCGTAAAGAAATTGCGGCGCTGCGTGACCGTCTGGCGCACAAAAAATGACAATTAATCATCGGCTCG | pCK_0402 |
| | na43 | CATGGAATTCCTCCTGTTAGCC | pCK_0402 |
| | na92 | AACGAAGTTGCGCAGCTGGAAAACGACGTTGCGGTTATCGAAAATGAAAACGCGTACCTGGAGAAGGAGATC | pNA42_BB1_SZ21half |
| | na43 | CATGGAATTCCTCCTGTTAGCC | pNA42_BB1_SZ21half |
| pNA43 | na96 | TGGAAAACGACGTTGCAGAAGTTGCGCCACAAGGAGAA | *X. nematophila* ATCC 19061 |
| | na7 | CGAGCCGATGATTAATTGTCACAGCGCCTCCACTTCG | *X. nematophila* ATCC 19061 |
| | jw61 | TGACAATTAATCATCGGCTCG | pCOLA_ara/tacI |
| | na102 | CGTTACGATTCAGTTTAACCGCAACACGGTTTTTGAGTTCCGCAACTTTCTGCATGGAATTCCTCCTGTTAGCC | pCOLA_ara/tacI |
| | jw61 | TGACAATTAATCATCGGCTCG | pNA41_BB1_SZ4half |
| | na103 | GCGTTACGGTTCTTCAGCTCTTCAACTTTGTTTTTCAGCTGTTCGTTACGATTCAGTTTAACCGC | pNA41_BB1_SZ4half |
| | jw61 | TGACAATTAATCATCGGCTCG | pNA41_BB2_SZ4half |
| | na95 | TTCTGCAACGTCGTTTTCCAGACGCGCAACCTCGTTCTCCAGGGTCGCCAGTTCGTTCTTGAGGTAAGCGTTACGGTTCTTCAGCTC | pNA41_BB2_SZ4half |
| pNA72 | na141 | ATCGCACGTCTGAAAAAAGAC | pNA40 |
| | na144 | GTCTTTTTTCAGACGTGCGATATAAATCTGGCGGGCGAA | pNA40 |
| pNA73 | na143 | CTGGAATCTCTGGAGAACAAAAAAG | pNA41 |
| | Na145 | CTTTTTTGTTCTCCAGAGATTCCAGCATGGAATTCCTCCTGTTAGCC | pNA41 |
| pNA145 | na304 | CATTGTTGTTAAAAACCTGGTCGAAAAAGGCTGAATTGC | pJW61 |
| | jw62 | CCAGGTTTTTAACAACAATGTGC | pJW61 |
| pNA146 | na305 | CTAACAGGAGGAATTCCATGCTGAAAGCCTTGGACCGC | pJW62 |
| | jw64 | CATGGAATTCCTCCTGTTAGCC | pJW62 |
| pNA147 | na306 | CATTGTTGTTAAAAACCTGGAATCGCATCGAACAGTTAAAACAG | pJW61 |
| | jw62 | CCAGGTTTTTAACAACAATGTGC | pJW61 |
| | na307 | CTAACAGGAGGAATTCCATGTTAAATGCCATTGACAAAGAGCTG | pJW62 |

| | | | |
|---|---|---|---|
| pNA148 | jw64 | CATGGAATTCCTCCTGTTAGCC | pJW62 |
| pNA149 | na308<br>na286 | GTTTGCCCGGCAGGTCTATAATGAGAACGAAACCCTGAAGAAAAAG<br>ATAGACCTGCCGGGCAAAC | pNA125<br>pNA125 |
| pNA150 | na309<br>na43 | CTAACAGGAGGAATTCCATGAAAGACAACCTGCAGCTGGAAC<br>CATGGAATTCCTCCTGTTAGCC | pNA126<br>pNA126 |
| pNA151 | na310<br>jw64 | CTAACAGGAGGAATTCCATGAATCGTAACGAACAGCTGAAAAAC<br>CATGGAATTCCTCCTGTTAGCC | pNA43<br>pNA43 |
| pNA152 | jw61<br>na286<br>na311<br>na312 | TGACAATTAATCATCGGCTCG<br>ATAGACCTGCCGGGCAAAC<br>GTTTGCCCGGCAGGTCTATAACGAGAAGGAGGAATTAAAATCG<br>CGAGCCGATGATTAATTGTCACTTGTAGGCTTCGATCTCCTTACG | pNA15<br>pNA15<br>pJW61<br>pJW61 |
| pNA153 | na315<br>na43<br>na313<br>na314 | CAAGCGCCACAAGGGGA<br>CATGGAATTCCTCCTGTTAGCC<br>GGCTAACAGGAGGAATTCCATGTTCTATGCTGAAGAGCGTGAACTG<br>TTCCCCTTGTGGCGCTTGTGAGATAGCTGCAGTCAGCTCG | pNA28<br>pNA28<br>pJW62<br>pJW62 |
| pNA154 | na316<br>na317 | AACGAGCTGACTGCAGCTATCTCAGGGTCTGGATCCGGCTCAGGTT CGGGATCATTATGTATTCATCAACTTTTTGAACAGC<br>TGAGATAGCTGCAGTCAGCTC | pJW62<br>pJW62 |
| pNA155 | na318<br>na317 | AACGAGCTGACTGCAGCTATCTCAGGTTCGGGATCATTATGTATTCA TCAACTTTTTGAACAGC<br>TGAGATAGCTGCAGTCAGCTC | pJW62<br>pJW62 |
| pNA156 | KB-pACYC-II-FW<br>na319 | AACGAGAAGGAGGAATTAAAATCG<br>CGATTTTAATTCCTCCTTCTCGTTCGATTCAGGATACACGGTTTCAG TGGCATTCCAGGTTTTTAACAACAATGTGC | pJW61<br>pJW61 |
| pNA157 | na320<br>na286 | GTTTGCCCGGCAGGTCTATGGGTCTGGATCCGGCTCAGGTTCGGG ATCAAACCTGGTTGCGCAGCTC<br>ATAGACCTGCCGGGCAAAC | pNA15<br>pNA15 |
| pNA158 | na321<br>na286 | GTTTGCCCGGCAGGTCTATGGTTCGGGATCAAACCTGGTTGCGCAG CTC<br>ATAGACCTGCCGGGCAAAC | pNA15<br>pNA15 |
| pNA159 | na322<br>na290 | GAAGTTGCGTCTCACGAACAGGGGTCTGGATCCGGCTCAGGTTCG GGATCAGCGGCTCCGCAGGG<br>CTGTTCGTGAGACGCAACTTC | pNA16<br>pNA16 |
| pNA160 | na323<br>na290 | GAAGTTGCGTCTCACGAACAGGGTTCGGGATCAGCGGCTCCGCAG GG<br>CTGTTCGTGAGACGCAACTTC | pNA16<br>pNA16 |
| pNA161 | na324<br>na142 | TTCGCCCGCCAGATTTATAAAGACAACCTGCAGCTGGAAC<br>ATAAATCTGGCGGGCGAA | pNA44<br>pNA44 |
| pNA162 | na325<br>na43 | GGCTAACAGGAGGAATTCCATGAACAAAAAAGAAGAACTGAAGAAC CG<br>CATGGAATTCCTCCTGTTAGCC | pNA45<br>pNA45 |
| pNA163 | jw61<br>na326 | TGACAATTAATCATCGGCTCG<br>CGAGCCGATGATTAATTGTCAACGCAGATTGGCGATCTTTTG | pJW61<br>pJW61 |

| | | | |
|---|---|---|---|
| pNA164 | jw63 | TTATGTATTCATCAACTTTTTGAACAGC | pJW62 |
| | na327 | GCTGTTCAAAAAGTTGATGAATACATAAGTTATCAAGGGCGCGAAGTT | pJW62 |
| pNA165 | na329 | GTTTGCCCGGCAGGTCTATGACCTGATCGCGTACCTGG | pNA15 |
| | na286 | ATAGACCTGCCGGGCAAAC | pNA15 |
| pNA166 | na330 | GGCTAACAGGAGGAATTCCATGAAAATCATCGCGAACCTGC | pNA16 |
| | na43 | CATGGAATTCCTCCTGTTAGCC | pNA16 |
| pNA167 | na331 | AATGAGAACGAAACCCTGAAGAAAAAG | pNA27 |
| | na333 | CTTTTTCTTCAGGGTTTCGTTCTCATTGTAAGCTTGGCGAGCAAAGG | pNA27 |
| pNA168 | na331 | AATGAGAACGAAACCCTGAAGAAAAAG | pNA30 |
| | na332 | CTTTTTCTTCAGGGTTTCGTTCTCATTATAATGCTGACGGGCAAACG | pNA30 |
| pNA169 | na309 | CTAACAGGAGGAATTCCATGAAAGACAACCTGCAGCTGGAAC | pNA31 |
| | na43 | CATGGAATTCCTCCTGTTAGCC | pNA31 |

### 3. Supplementary Figures

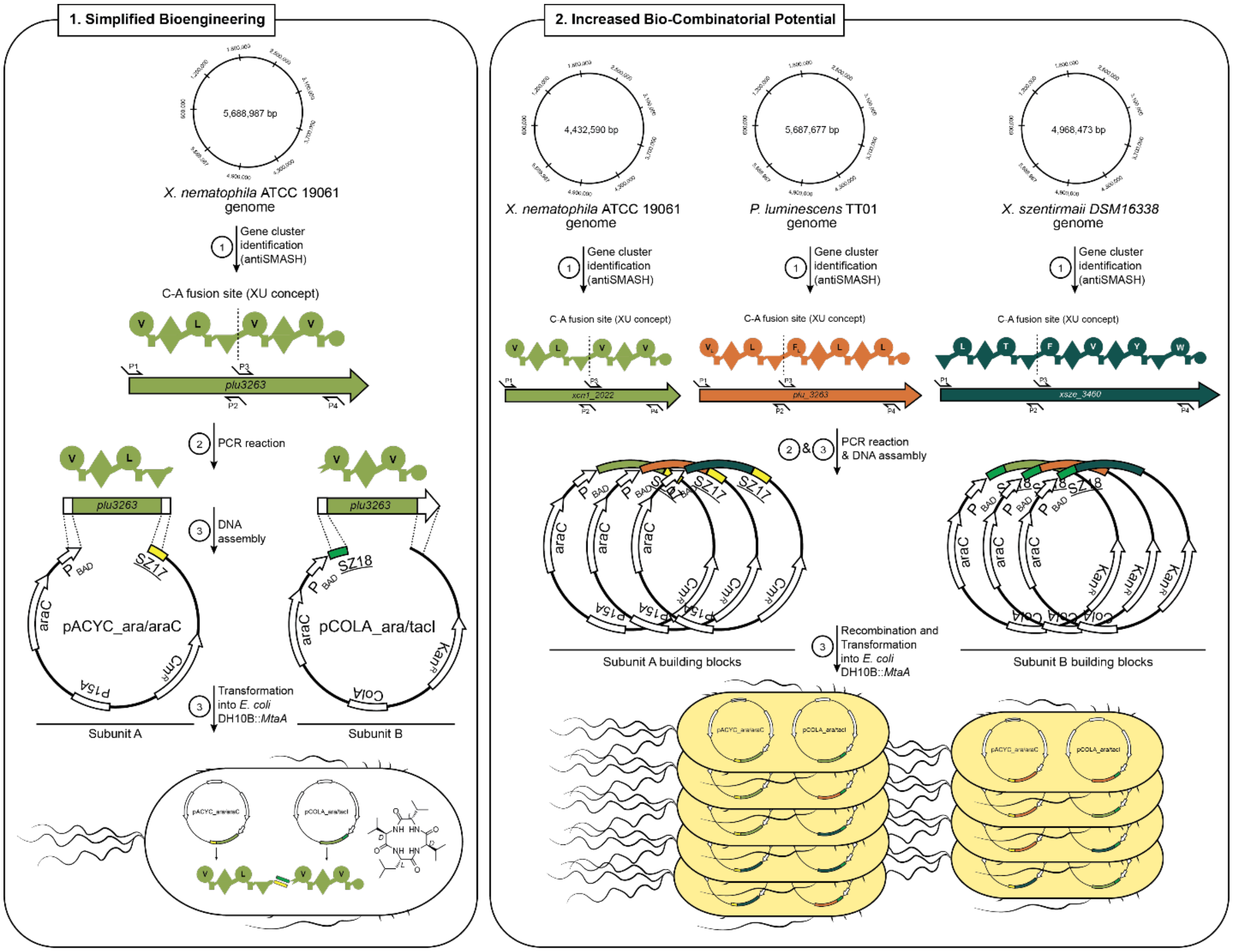

**Figure S1. Advantages of Type S NRPS. 1) Simplified bioengineering:** Splitting NRPS into two or three independently expressed SZ linked subunits enables easier and faster cloning. Traditional NRPS engineering often requires elaborated cloning strategies (yeast cloning,[7] LLHR[8] or ExoCET[9]) which are frequently accompanied with technical problems and limitation. By breaking NRPSs into smaller subunits, cloning can be simplified, making standard strategies such as Gibson[10], HiFi and Hot Fusion[11] assembly sufficient. **2) Increased bio-combinatorial potential:** With SZs, type S NRPSs can be created faster and to a greater extent than before, as the number of artificial NRPSs increases exponentially with the number of subunits. Once generated, subunits can be reused at any time and for any experimental approaches without any additional cloning efforts.

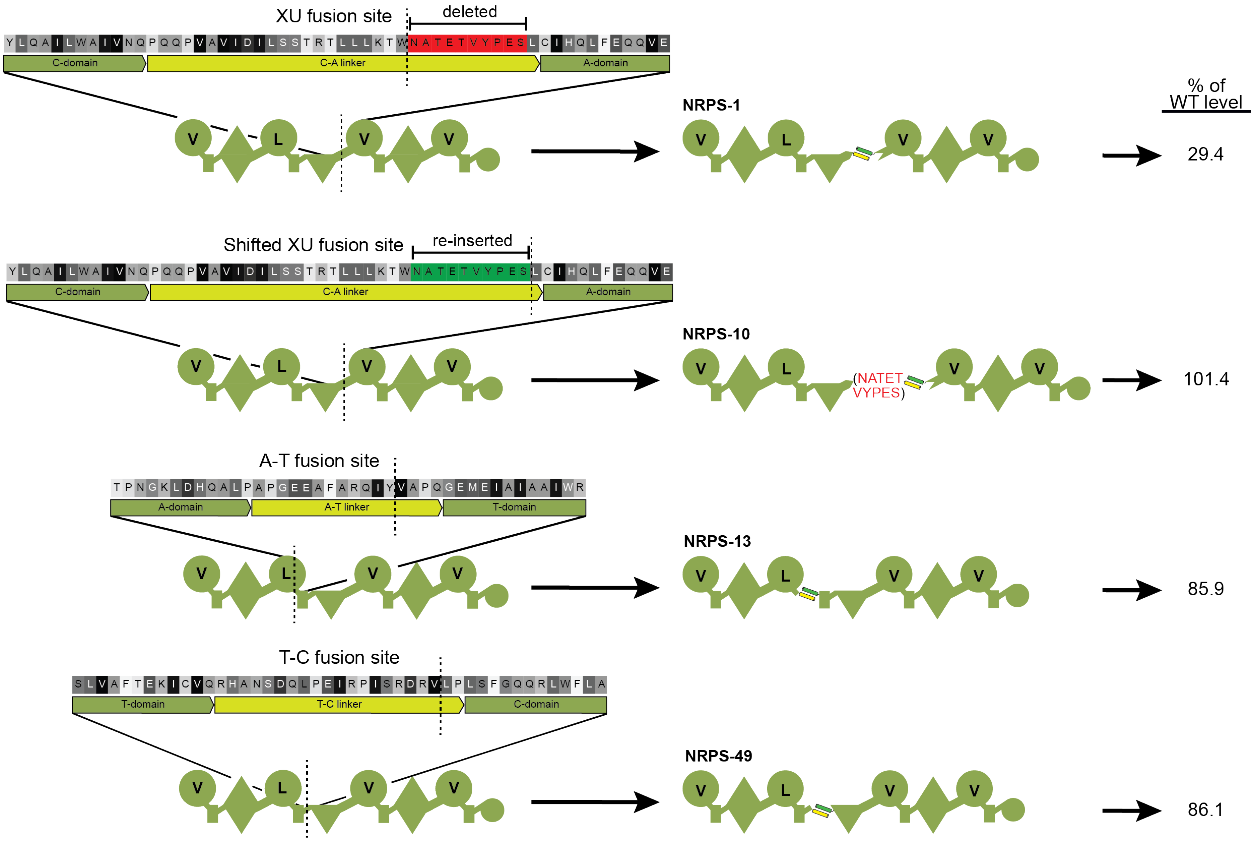

**Figure S2. Other splicing positions**. SZ17:18 Introduction at three different positions within the C-A, T-C and A-T linker region to create two protein type S XtpS variants. AS sequences and exact SZ17:18 introduction sites are highlighted with a vertical dashed line. Initially, for the introduction of SZs into the C-A position, 10 AAs were deleted (highlighted in red) to meet the distance between the C- and A-domain. Re-insertion of the 10 AAs (highlighted in green) and shifted fusion site restored peptide production. Production of 1 (cyclo (vLvV)) relative to WT level are indicated on the right hand site.

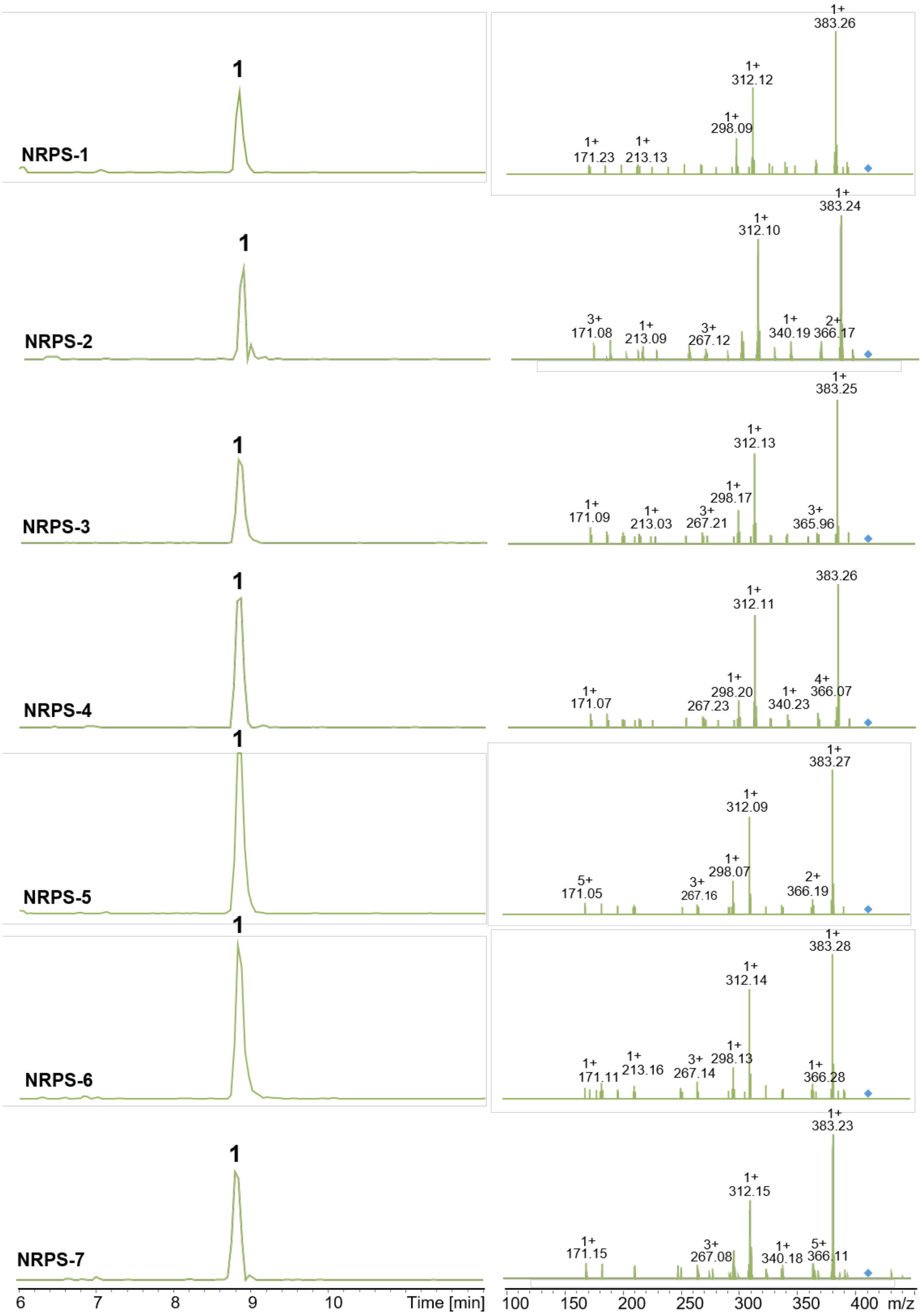

NRPS-1
1
1+ 383.26
1+ 312.12
1+ 298.09
1+ 171.23
1+ 213.13
NRPS-2
1
1+ 383.24
1+ 312.10
3+ 171.08
1+ 213.09
3+ 267.12
1+ 340.19
2+ 366.17
NRPS-3
1
1+ 383.25
1+ 312.13
1+ 298.17
1+ 171.09
1+ 213.03
3+ 267.21
3+ 365.96
NRPS-4
1
383.26
1+ 312.11
1+ 298.20
1+ 171.07
267.23
1+ 340.23
4+ 366.07
NRPS-5
1
1+ 383.27
1+ 312.09
1+ 298.07
5+ 171.05
3+ 267.16
2+ 366.19
NRPS-6
1
1+ 383.28
1+ 312.14
1+ 298.13
1+ 171.11
1+ 213.16
3+ 267.14
1+ 366.28
NRPS-7
1
1+ 383.23
1+ 312.15
1+ 171.15
3+ 267.08
1+ 340.18
5+ 366.11
6
7
8
9
10
Time [min]
100
150
200
250
300
350
400
m/z

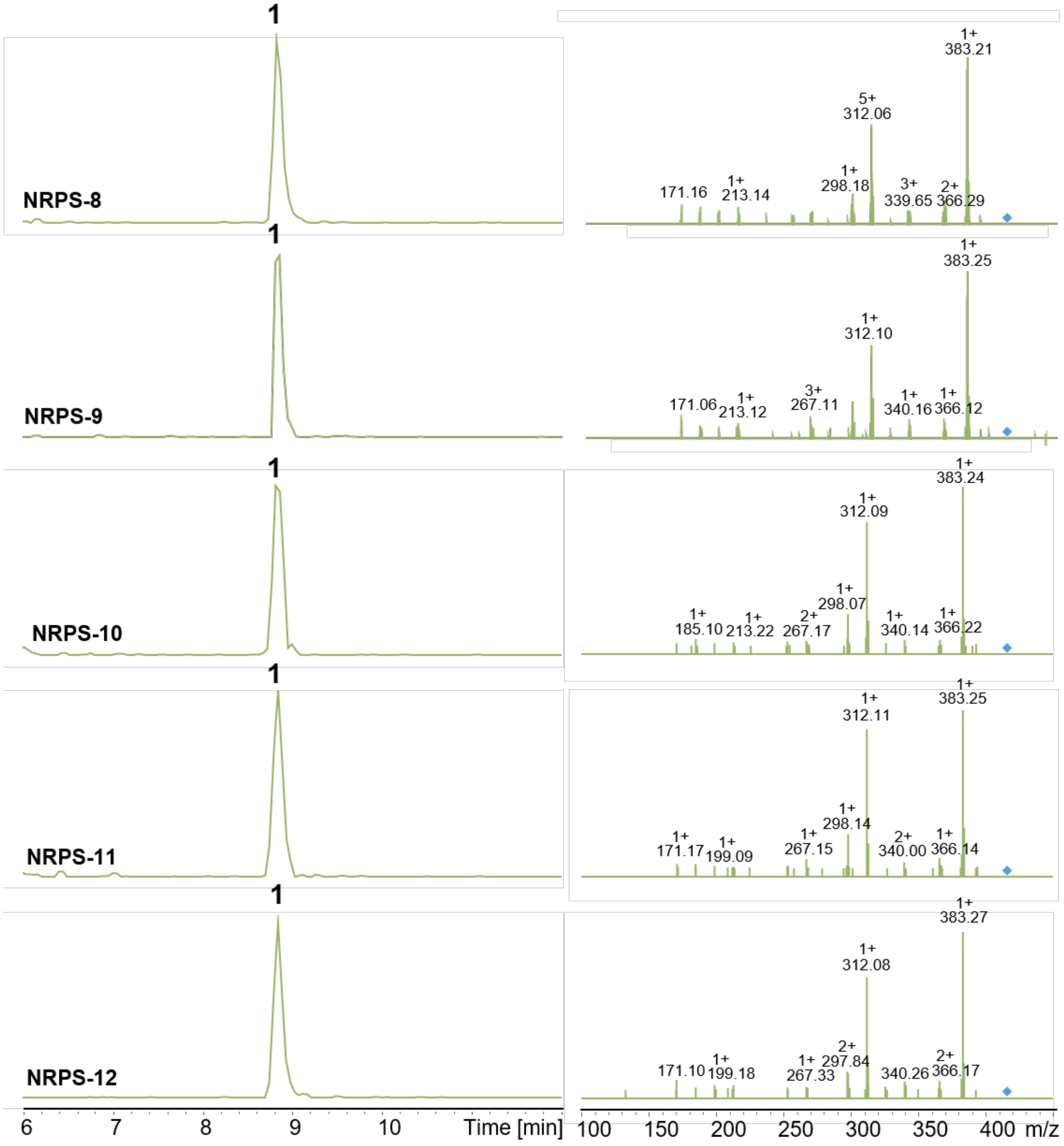

**Figure S3. HPLC/MS data (Figure 2) of compound 1 produced in *E. coli* DH10B::*mtaA*.** EIC/MS$^2$ of 1 (m/z $[M+H]^+$ = 411.29) produced by NRPS-1 to -12.

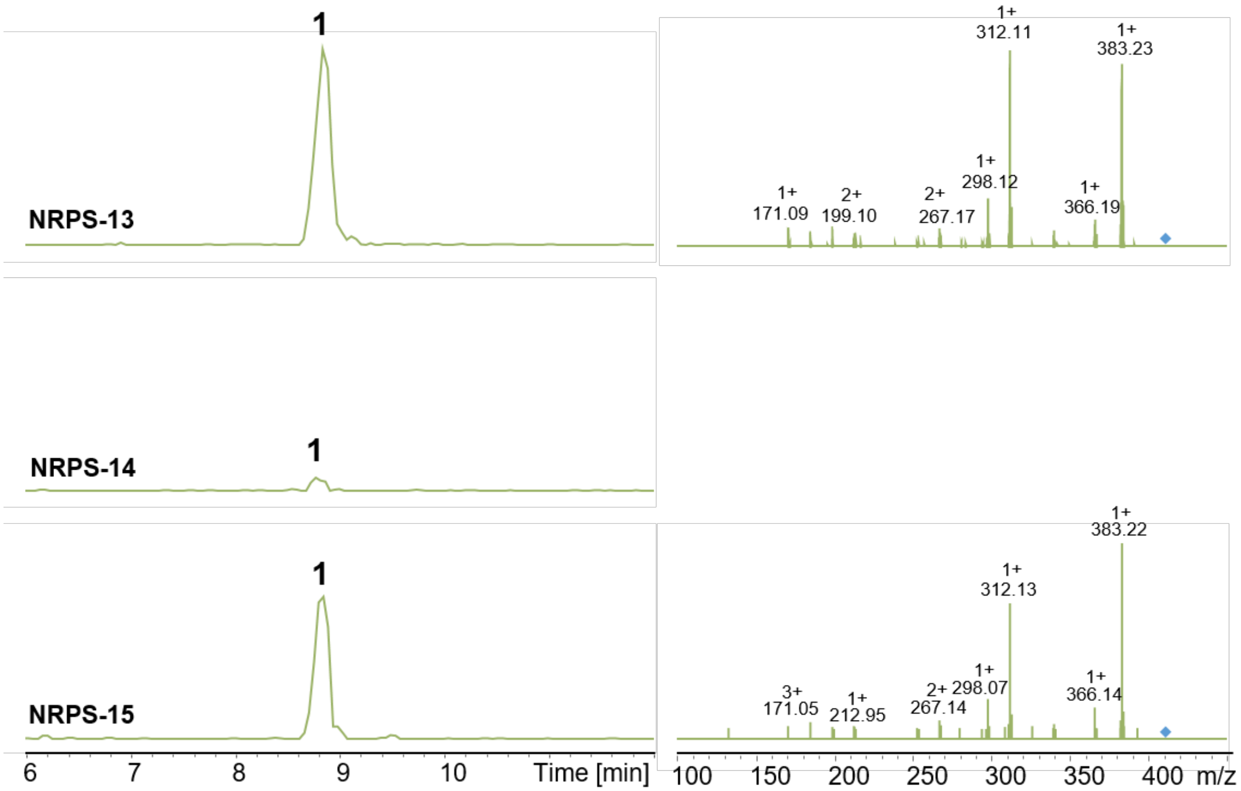

**Figure S4. HPLC/MS data (Figure 3) of compounds 1 produced in *E. coli* DH10B::*taA*.** EIC/MS[2] of 1 (m/z $[M+H]^+$ = 411.29) produced by NRPS-13 to -15.

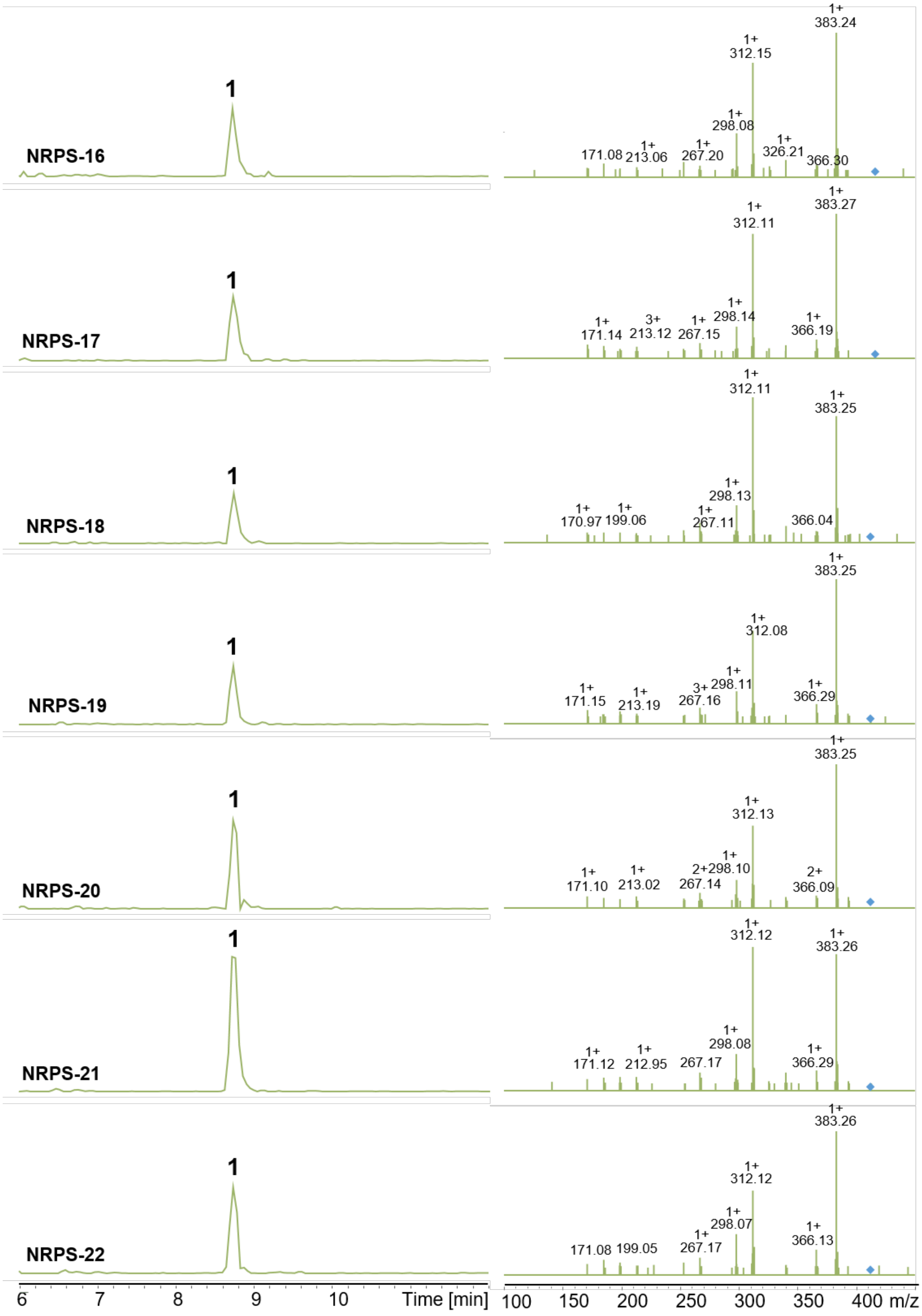
NRPS-16
NRPS-17
NRPS-18
NRPS-19
NRPS-20
NRPS-21
NRPS-22
1
Time [min]
m/z
383.24
312.15
298.08
326.21
366.30
171.08
213.06
267.20
383.27
312.11
298.14
171.14
213.12
267.15
366.19
383.25
298.13
170.97
199.06
267.11
366.04
312.08
298.11
171.15
213.19
267.16
366.29
312.13
298.10
171.10
213.02
267.14
366.09
312.12
383.26
298.08
171.12
212.95
267.17
298.07
171.08
199.05
366.13

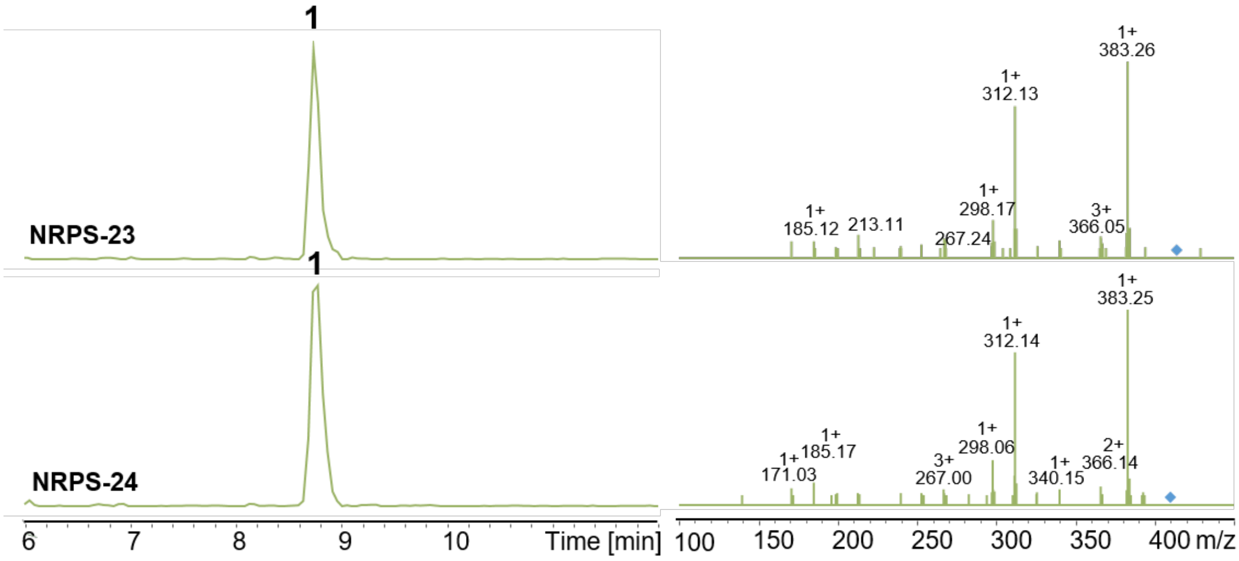

**Figure S5. HPLC/MS data (Figure 4) of compound 1 produced in *E. coli* DH10B::*mtaA*.** EIC/MS$^2$ of 1 (m/z $[M+H]^+$ = 411.29) produced by NRPS-16 to -24.

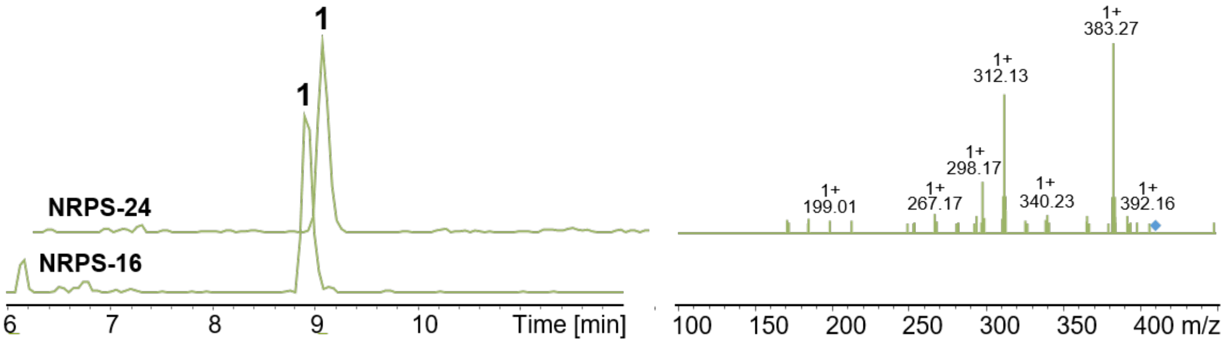

**Figure S6. HPLC/MS data (Figure 5) of compound 1 produced in *E. coli* DH10B::*mtaA*.** EIC/MS2 of 1 (m/z [M+H]+ = 411.29) produced by NRPS-24. EIC of 1 produced by NRPS-16**.**

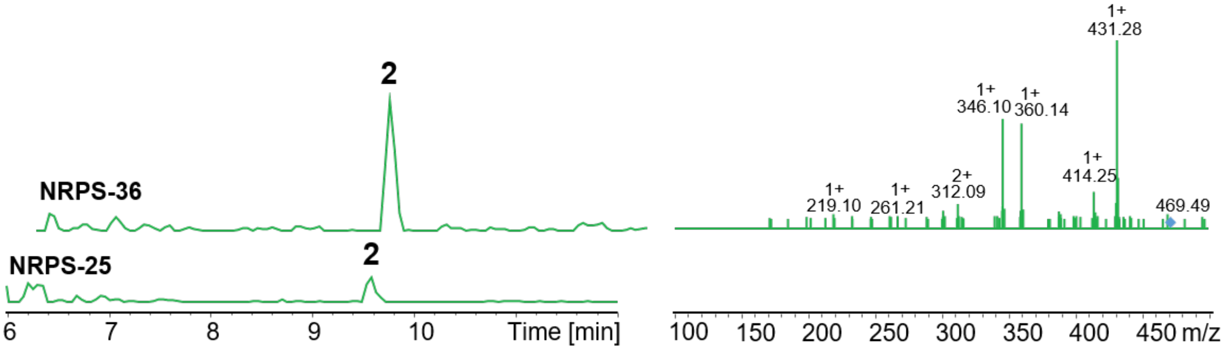

**Figure S7. HPLC/MS data (Figure 5) of compound 2 produced in E. coli DH10B::mtaA.** EIC/MS2 of 2 (m/z [M+H]+ = 459.30) produced by NRPS-36. EIC of 1 produced by NRPS-25.

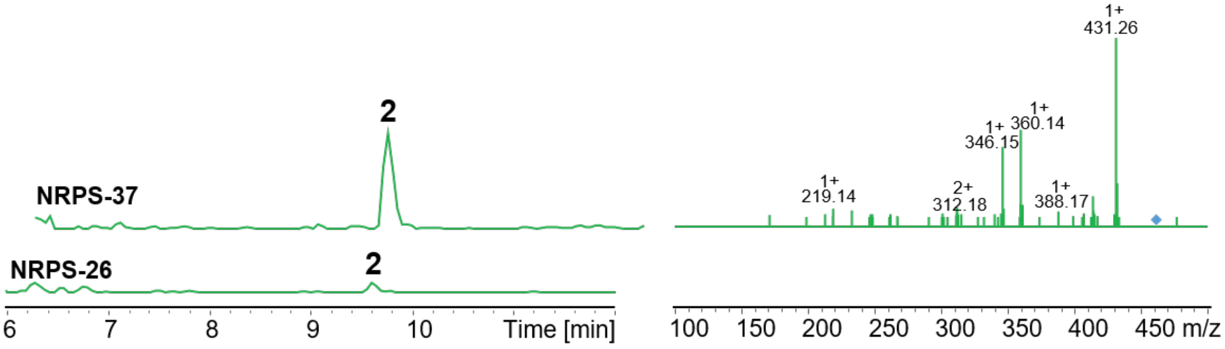

**Figure S8. HPLC/MS data (Figure 5) of compound 2 produced in *E. coli* DH10B::*mtaA*.** EIC/MS$^2$ of 2 (m/z [M+H]$^+$ = 459.30) produced by NRPS-37. EIC of 1 produced by NRPS-26.

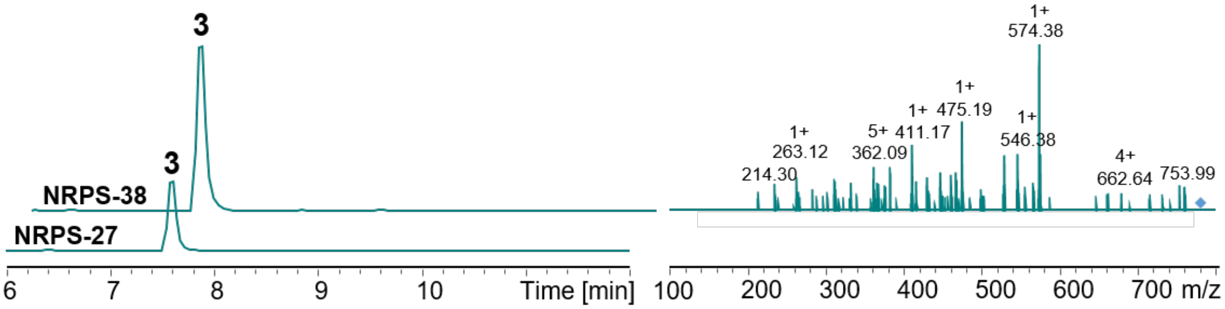

**Figure S9. HPLC/MS data (Figure 5) of compound 3 produced in *E. coli* DH10B::*mtaA*.** EIC/MS$^2$ of 3 (m/z [M+H]$^+$ = 778,45) produced by NRPS-38. EIC of 3 produced by NRPS-27.

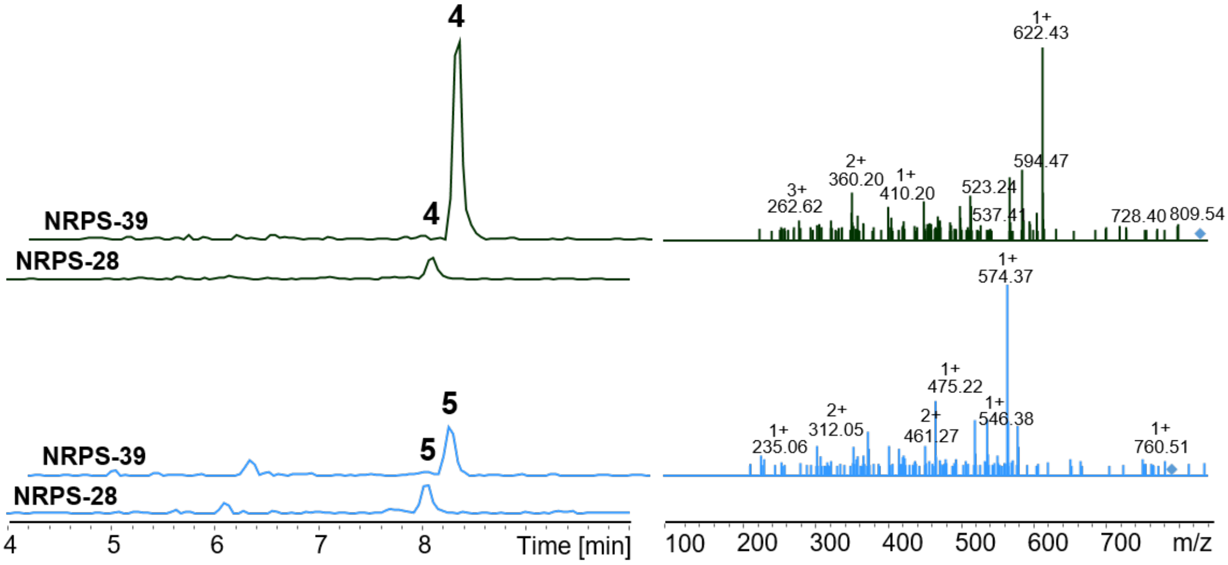

**Figure S10. HPLC/MS data (Figure 5) of compounds 4 and 5 produced in *E. coli* DH10B::*mtaA*.** EIC/MS$^2$ of 4 (m/z $[M+H]^+$ = 826.45) and 5 (m/z $[M+H]^+$ = 792.47) produced by NRPS-38. EIC of 4 and 5 produced by NRPS-28.

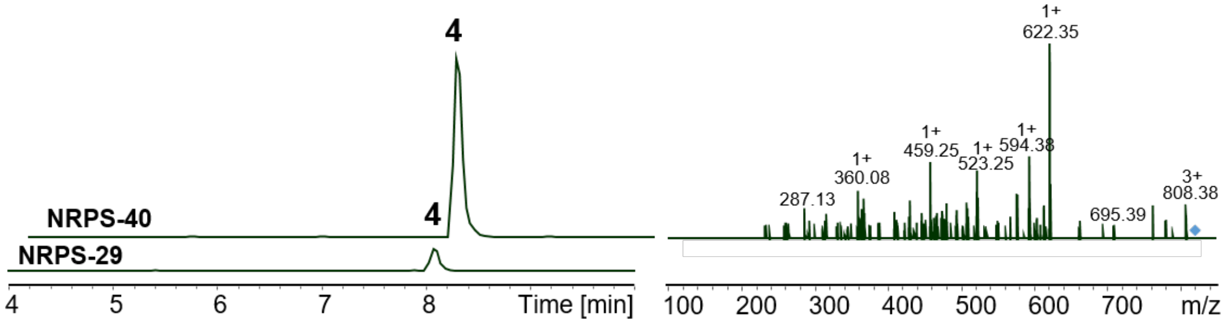

**Figure S11. HPLC/MS data (Figure 5) of compound 4 produced in *E. coli* DH10B::*mtaA*.** EIC/MS$^2$ of 4 (m/z $[M+H]^+$ = 826.45) produced by NRPS-38. EIC of 4 produced by NRPS-28.

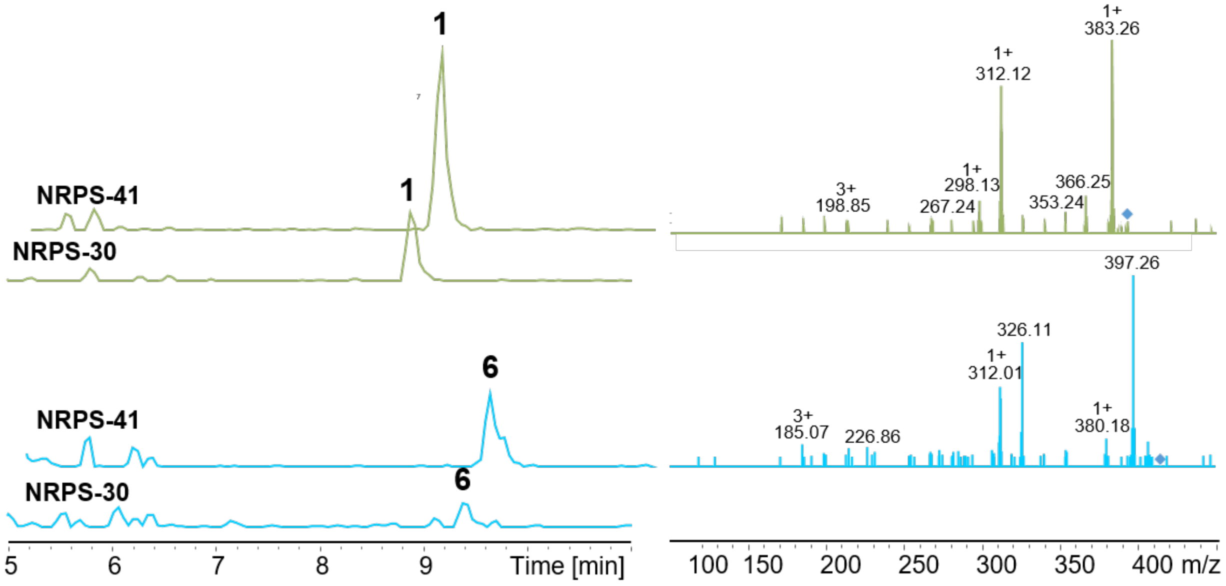

**Figure S12. HPLC/MS data (Figure 5) of compounds 1 and 6 produced in E. coli DH10B::mtaA.** EIC/MS$^2$ of 1 (m/z $[M+H]^+$ = 411.29) and 6 (m/z $[M+H]^+$ = 425.31) produced by NRPS-41. EIC of 1 and 6 produced by NRPS-30.

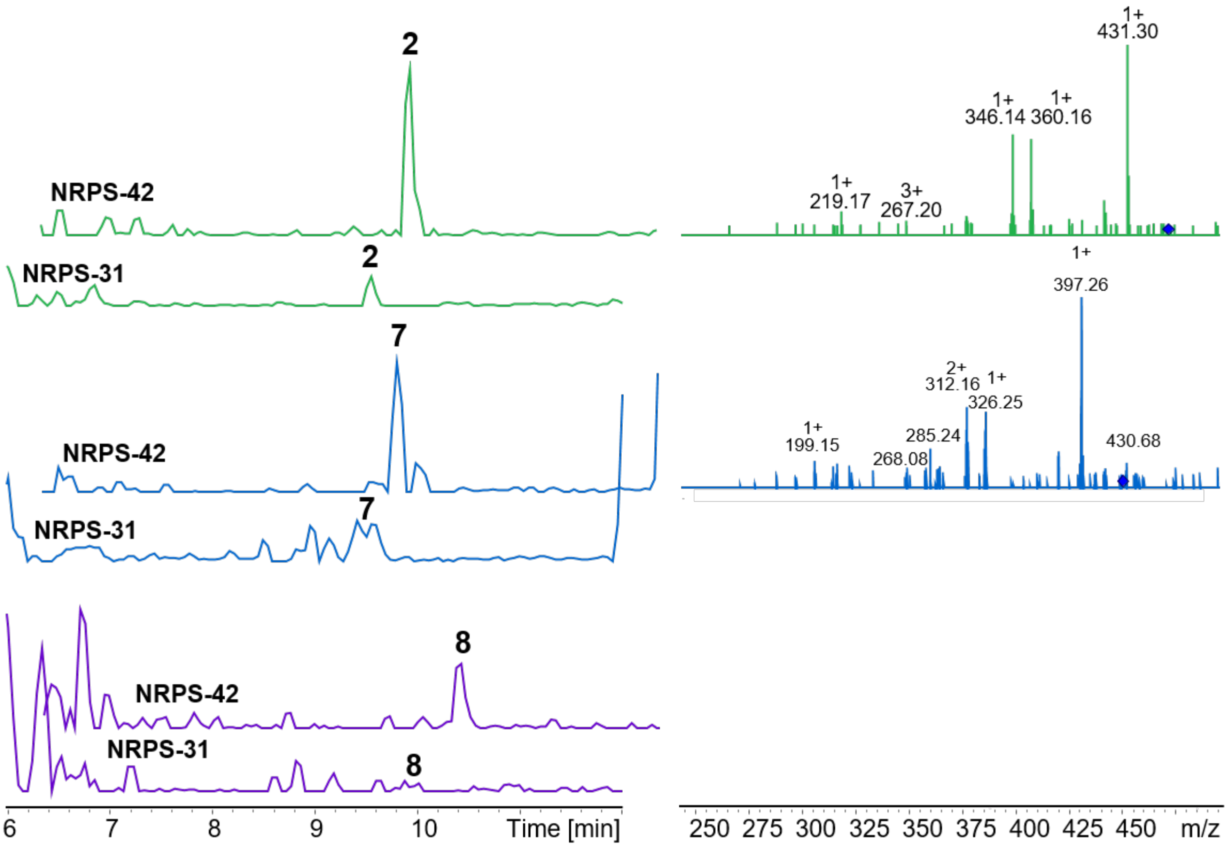

**Figure S13. HPLC/MS data (Figure 5) of compounds 2, 7 and 8 produced in *E. coli* DH10B::*mtaA*.** EIC/MS$^2$ of 2 (m/z $[M+H]^+$ = 459.30), 7 (m/z $[M+H]^+$ = 425.31) and 8 m/z $[M+H]^+$ = 472.31) produced by NRPS-42. EIC of 2, 7 and 8 produced by NRPS-31.

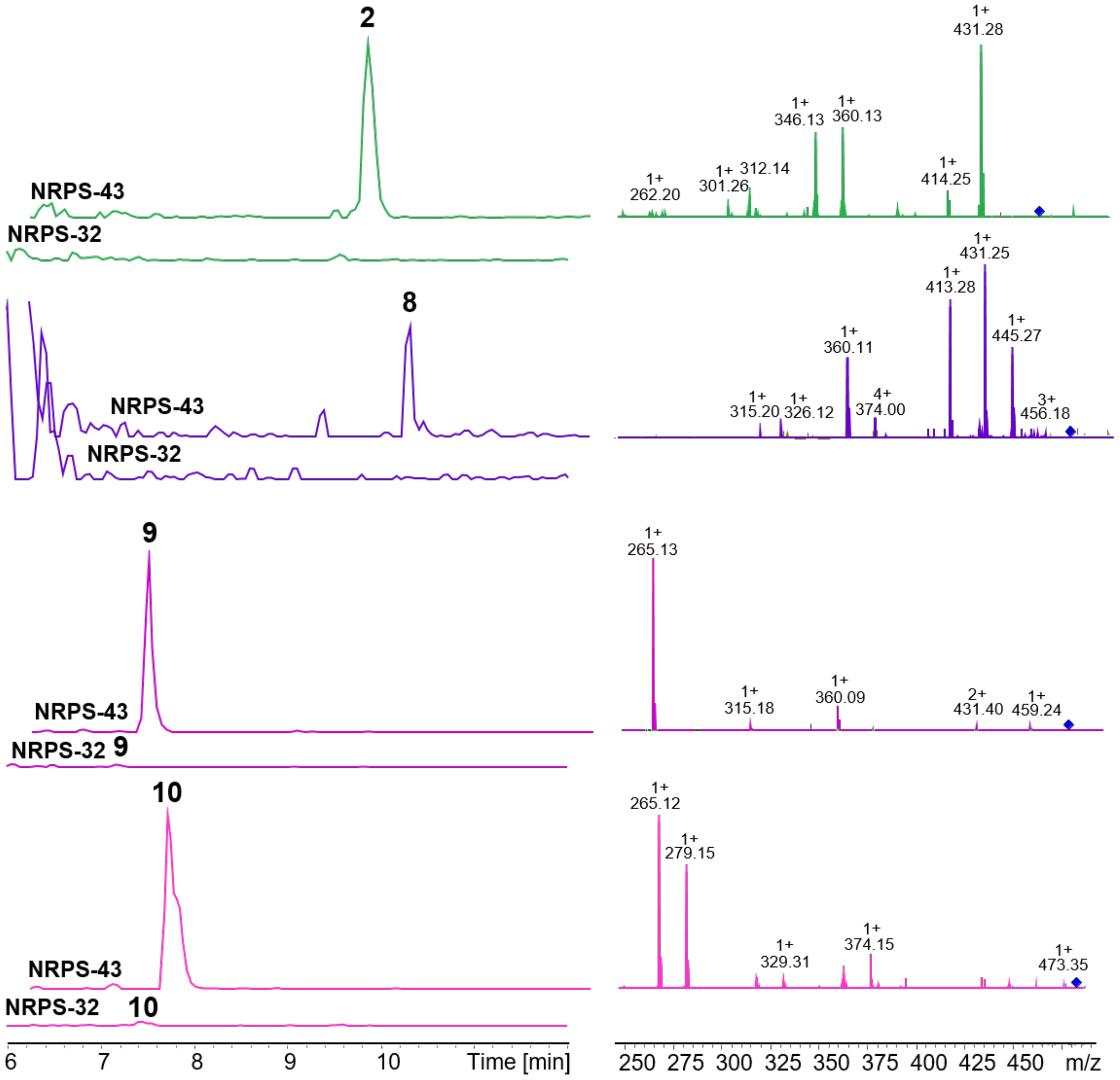

**Figure S14. HPLC/MS data (Figure 5) of compounds 2, 8, 9 and 10 produced in *E. coli* DH10B::*mtaA*.** EIC/MS$^2$ of 2 (m/z [M+H]$^+$ = 459.30), 8 (m/z [M+H]$^+$ = 472.31), 9 (m/z [M+H]$^+$ = 476.62) and 10 m/z [M+H]$^+$ = 490.65) produced by NRPS-43. EIC of 2, 8, 9 and 10 produced by NRPS-32.

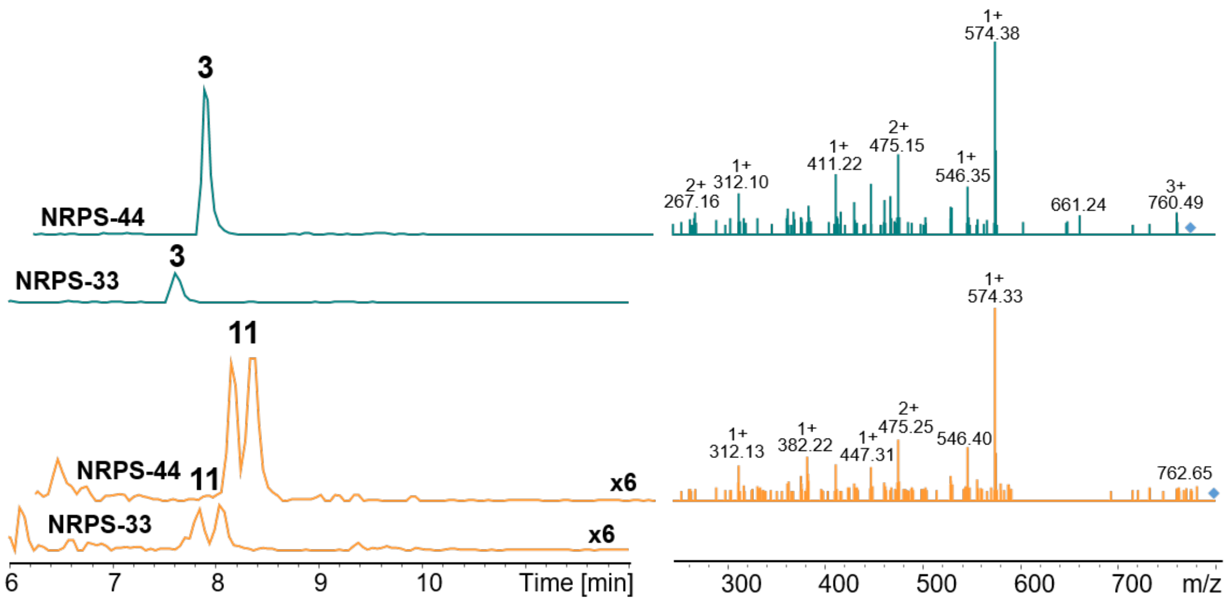

**Figure S15. HPLC/MS data (Figure 5) of compounds 3 and 11 produced in E. coli DH10B::mtaA.** EIC/MS$^2$ of 3 (m/z $[M+H]^+$ = 778.45) and 11 (m/z $[M+H]^+$ = 792.47) produced by NRPS-44. EIC of 3 and 11 produced by NRPS-33.

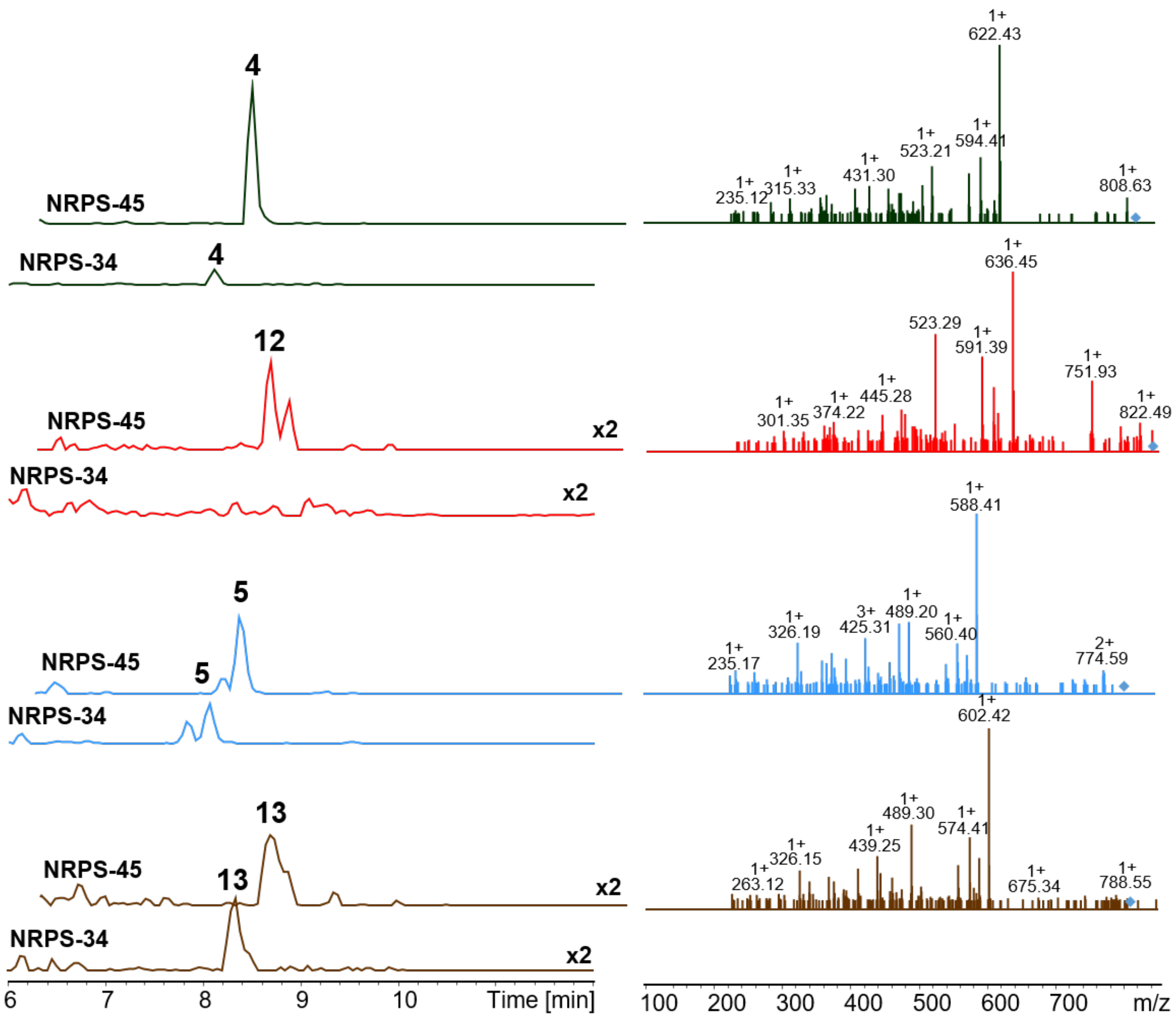

4
NRPS-45
NRPS-34
4
12
NRPS-45
x2
NRPS-34
x2
5
NRPS-45
5
NRPS-34
13
NRPS-45
13
x2
NRPS-34
x2
6
7
8
9
10
Time [min]
1+
622.43
1+
594.41
1+
523.21
1+
431.30
1+
315.33
1+
235.12
1+
808.63
1+
636.45
523.29
1+
591.39
1+
751.93
1+
445.28
1+
374.22
1+
301.35
1+
822.49
1+
588.41
1+
489.20
3+
425.31
1+
326.19
1+
560.40
1+
235.17
2+
774.59
1+
602.42
1+
489.30
1+
574.41
1+
439.25
1+
326.15
1+
263.12
1+
675.34
1+
788.55
100
200
300
400
500
600
700
m/z

**Figure S16. HPLC/MS data (Figure 5) of compounds 4, 12, 5 and 13 produced in *E. coli* DH10B::*mtaA*.** EIC/MS$^2$ of 4 (m/z $[M+H]^+$ = 826.45), 12 (m/z $[M+H]^+$ = 840.47), 5 (m/z $[M+H]^+$ = 792.47) and 13 m/z $[M+H]^+$ = 806.48) produced by NRPS-45. EIC of 2, 12, 5 and 13 produced by NRPS-34.

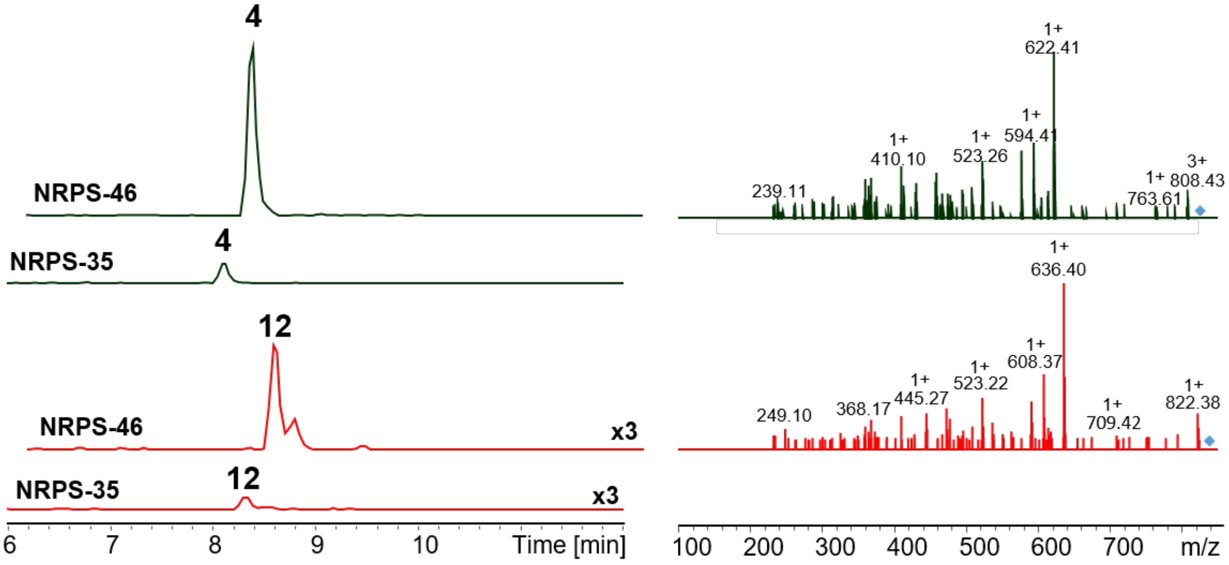

**Figure S17**. **HPLC/MS data (Figure 5) of compounds 4 and 12 produced in *E. coli* DH10B::*mtaA*.** EIC/MS$^2$ of 4 (m/z $[M+H]^+$ = 826.45) and 12 (m/z $[M+H]^+$ = 840.47) produced by NRPS-46. EIC of 4 and 12 produced by NRPS-35.

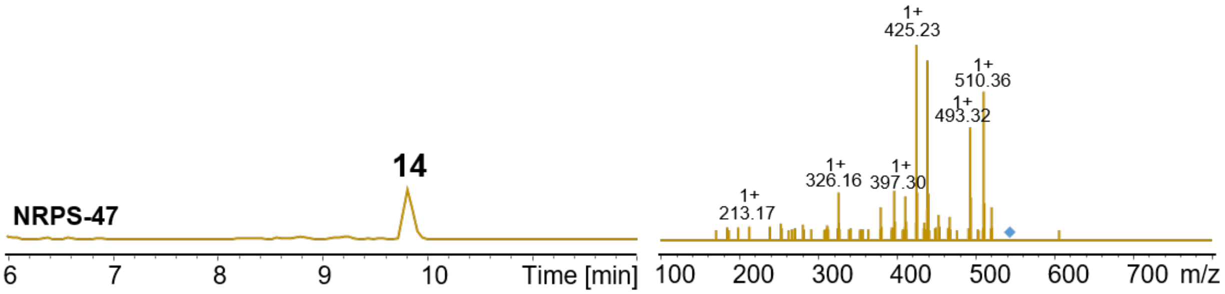

**Figure S18. HPLC/MS data (Figure 6) of compound 14 produced in *E. coli* DH10B::*mtaA*.** EIC/MS$^2$ of 14 (m/z $[M+H]^+$ = 538.40) produced by NRPS-47.

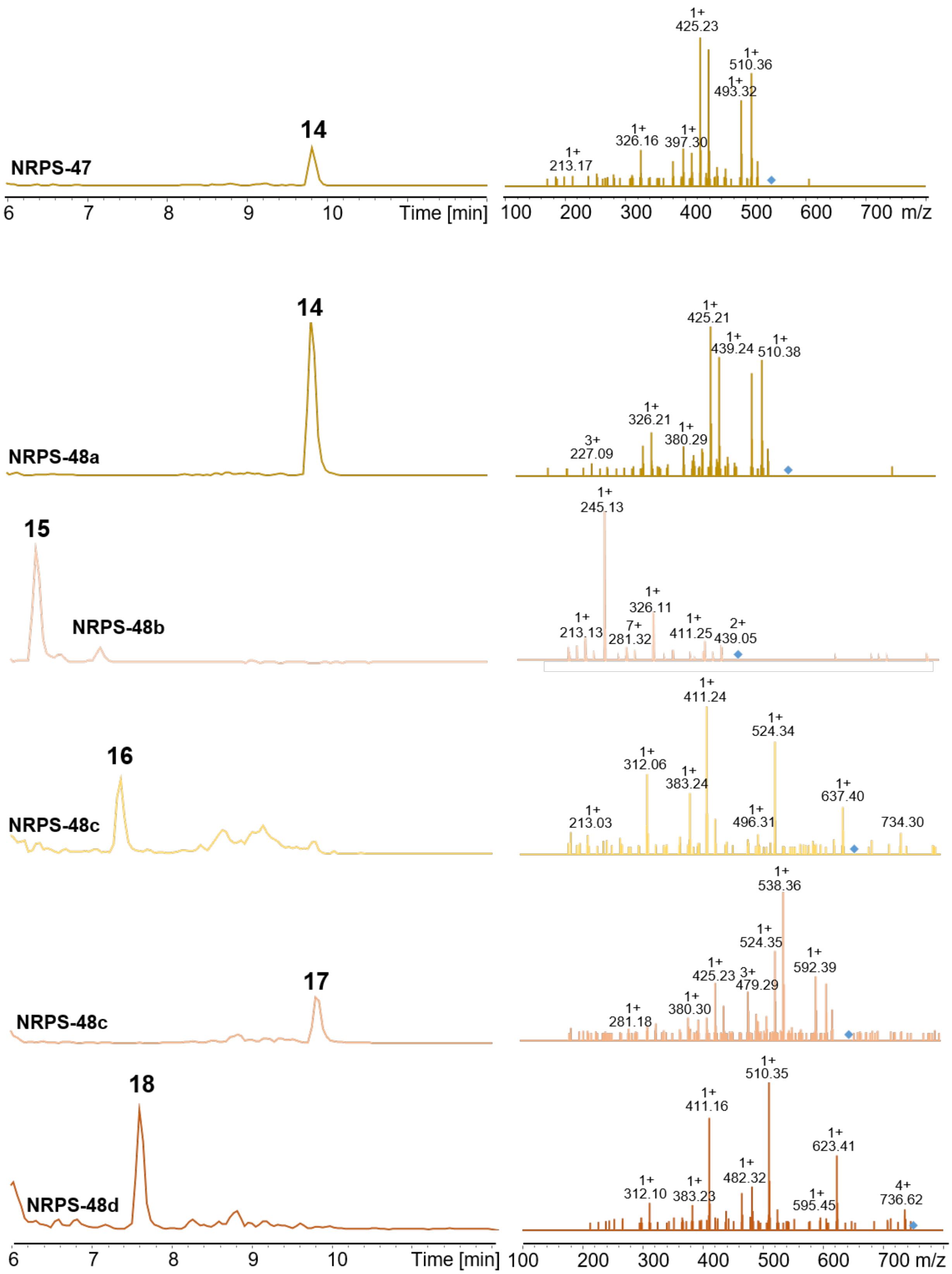
NRPS-47
14
NRPS-48a
14
15
NRPS-48b
16
NRPS-48c
17
NRPS-48c
18
NRPS-48d
Time [min]
m/z
1+ 425.23
1+ 510.36
1+ 493.32
1+ 326.16
1+ 397.30
1+ 213.17
1+ 425.21
1+ 439.24
1+ 510.38
1+ 326.21
1+ 380.29
3+ 227.09
1+ 245.13
1+ 326.11
1+ 213.13
7+ 281.32
1+ 411.25
2+ 439.05
1+ 411.24
1+ 524.34
1+ 312.06
1+ 383.24
1+ 637.40
1+ 213.03
1+ 496.31
734.30
1+ 538.36
1+ 524.35
1+ 425.23
3+ 479.29
1+ 592.39
1+ 380.30
1+ 281.18
1+ 510.35
1+ 411.16
1+ 623.41
1+ 482.32
1+ 312.10
1+ 383.23
1+ 595.45
4+ 736.62

**Figure S19. HPLC/MS data (Figure 5) of compounds 14, 15, 16, 17 and 18 produced in *E. coli* DH10B::*mtaA*.** EIC/MS$^2$ of 14 (m/z $[M+H]^+$ = 538.40) produced by NRPS-47 and -48a. EIC/MS$^2$ of 15 (m/z $[M+H]^+$ = 470.35) produced by NRPS-47. EIC/MS$^2$ of 16 (m/z $[M+H]^+$ = 655.47) produced by NRPS-48c. EIC/MS$^2$ of 17 (m/z $[M+7]^+$ = 637.46) produced by NRPS-48c. EIC/MS$^2$ of 18 m/z $[M+H]^+$ = 754.54) produced by NRPS-48d.

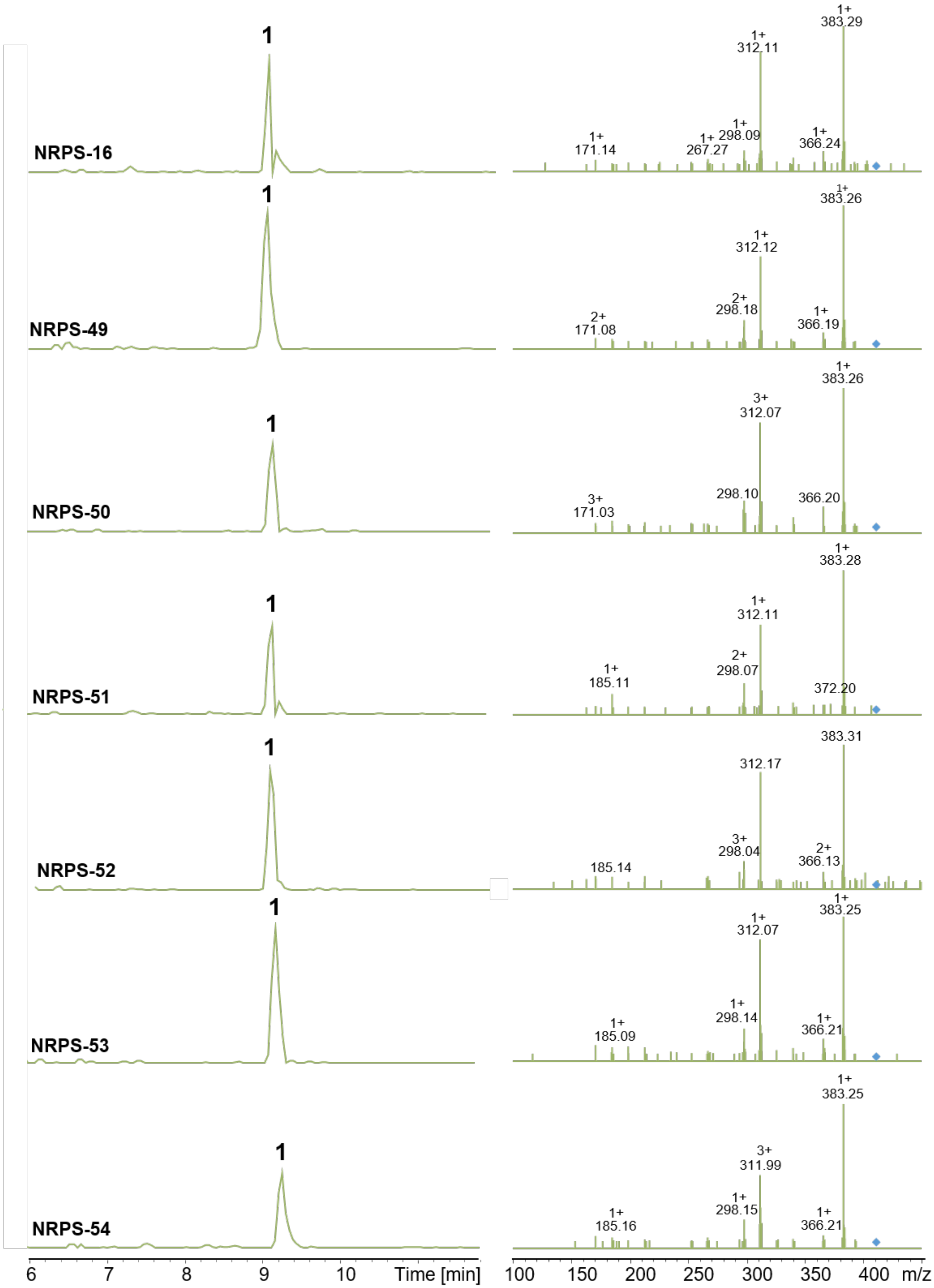

NRPS-16
NRPS-49
NRPS-50
NRPS-51
NRPS-52
NRPS-53
NRPS-54
1
1+ 383.29
1+ 312.11
1+ 298.09
1+ 267.27
1+ 171.14
1+ 366.24
1+ 383.26
1+ 312.12
2+ 298.18
2+ 171.08
1+ 366.19
1+ 383.26
3+ 312.07
298.10
3+ 171.03
366.20
1+ 383.28
1+ 312.11
2+ 298.07
1+ 185.11
372.20
383.31
312.17
3+ 298.04
185.14
2+ 366.13
1+ 383.25
1+ 312.07
1+ 298.14
1+ 185.09
1+ 366.21
1+ 383.25
3+ 311.99
1+ 298.15
1+ 185.16
1+ 366.21
6
7
8
9
10
Time [min]
100
150
200
250
300
350
400
m/z

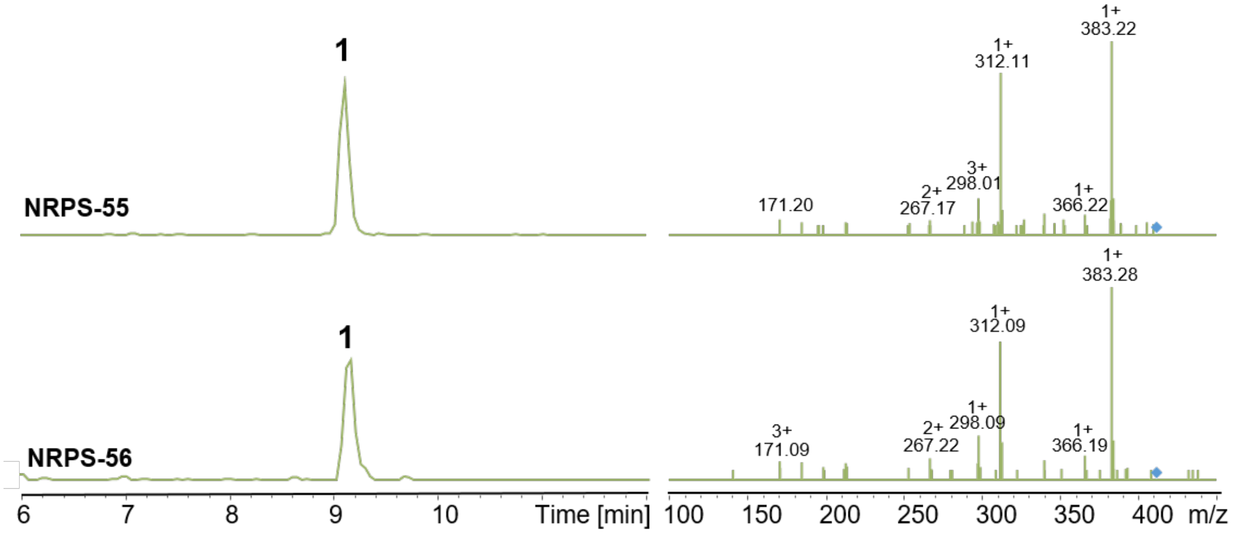

**Figure S20. HPLC/MS data (Figure S25) of compound 1 produced in *E. coli* DH10B::*mtaA*.** EIC/MS$^2$ of 1 (m/z $[M+H]^+$ = 411.29) produced by NRPS-16 and NRPS-49 to -56.

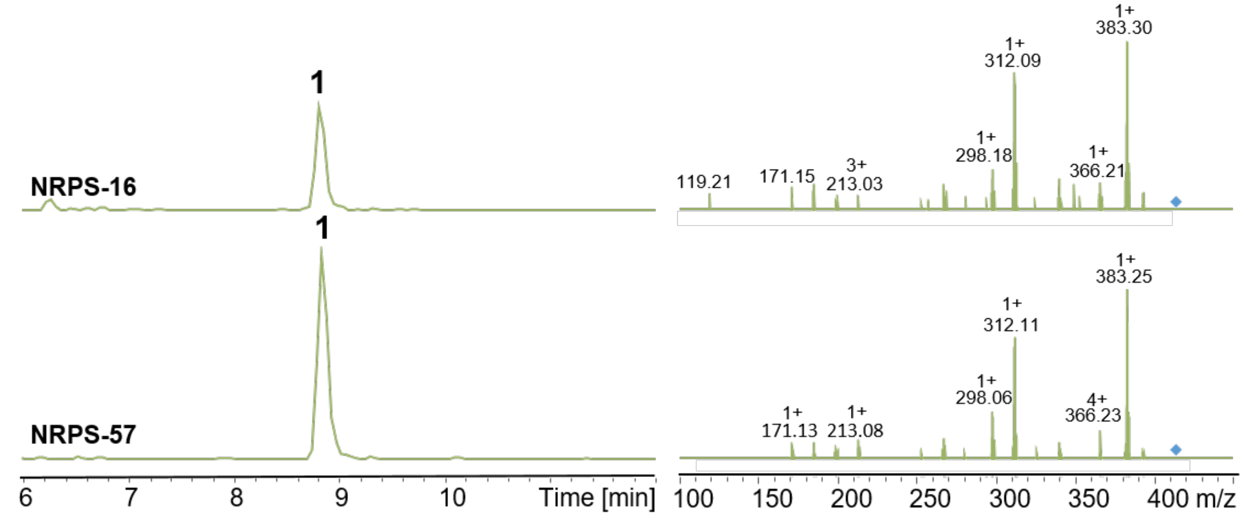

**Figure S21. HPLC/MS data (Figure S26) of compound 1 produced in E. coli DH10B::mtaA.** EIC/MS$^2$ of 1 (m/z $[M+H]^+$ = 411.29) produced by NRPS-16 and NRPS-57.

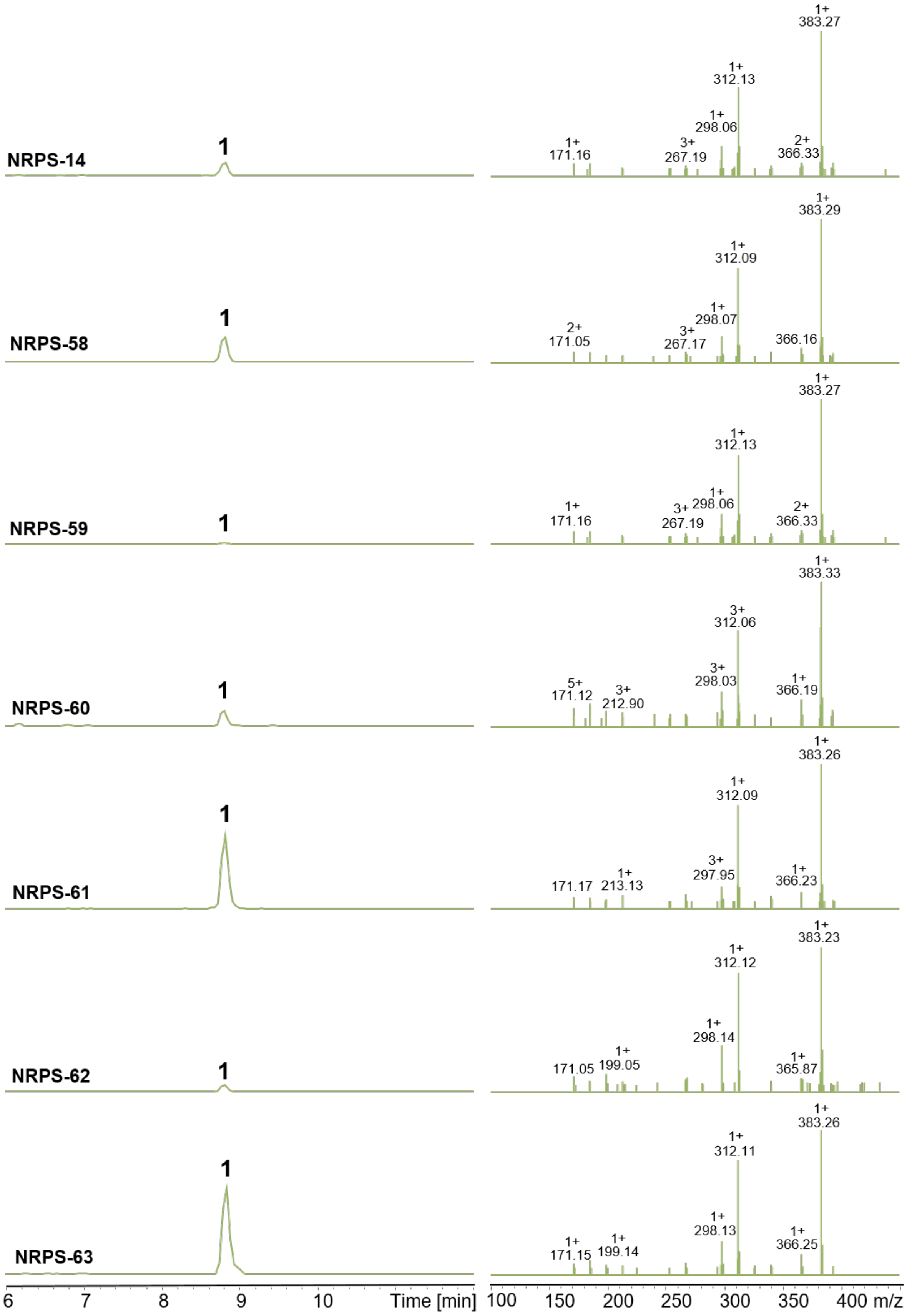
NRPS-14
NRPS-58
NRPS-59
NRPS-60
NRPS-61
NRPS-62
NRPS-63
1
Time [min]
m/z
1+ 383.27
1+ 312.13
1+ 298.06
3+ 267.19
1+ 171.16
2+ 366.33
1+ 383.29
1+ 312.09
1+ 298.07
3+ 267.17
2+ 171.05
366.16
1+ 383.27
1+ 312.13
1+ 298.06
3+ 267.19
1+ 171.16
2+ 366.33
1+ 383.33
3+ 312.06
3+ 298.03
5+ 171.12
3+ 212.90
1+ 366.19
1+ 383.26
1+ 312.09
3+ 297.95
171.17
1+ 213.13
1+ 366.23
1+ 383.23
1+ 312.12
1+ 298.14
171.05
1+ 199.05
1+ 365.87
1+ 383.26
1+ 312.11
1+ 298.13
1+ 171.15
1+ 199.14
1+ 366.25

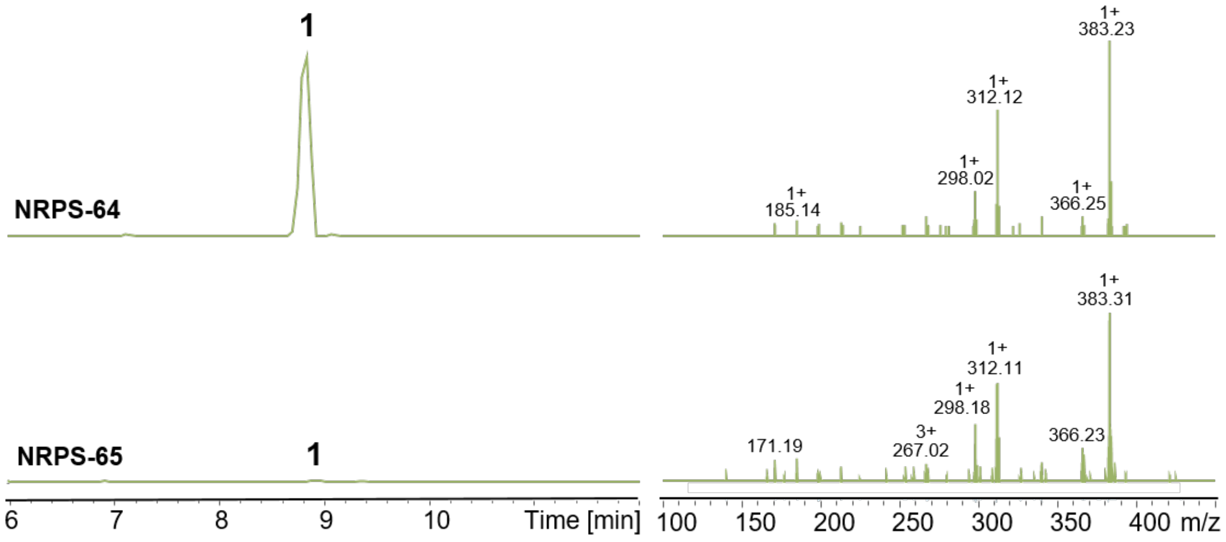

**Figure S22. HPLC/MS data (Figure S27) of compound 1 produced in *E. coli* DH10B::*mtaA*.** EIC/MS$^2$ of 1 (m/z $[M+H]^+$ = 411.29) produced by NRPS-14 and NRPS-58 to NRPS-65.

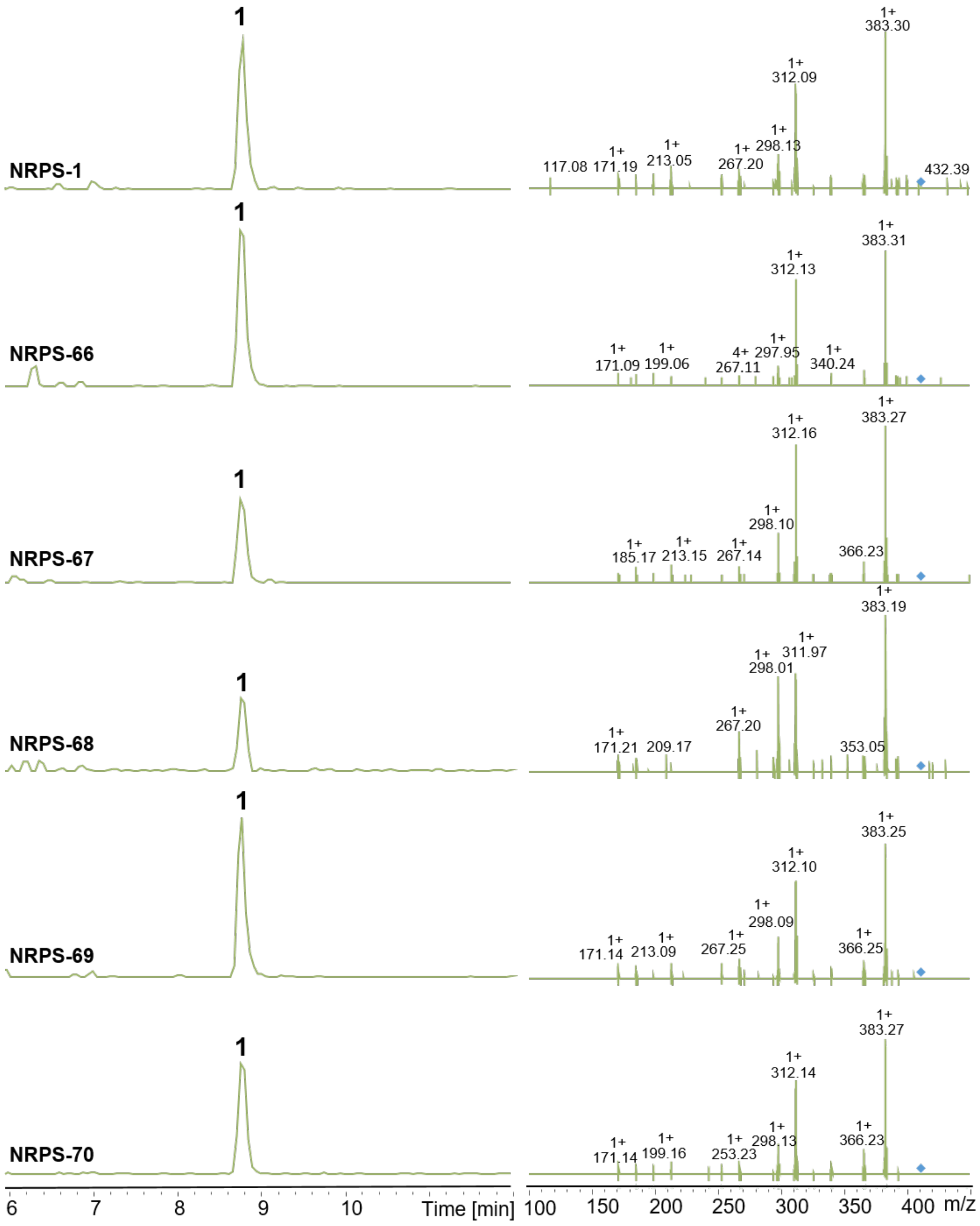

NRPS-1
NRPS-66
NRPS-67
NRPS-68
NRPS-69
NRPS-70
1
6
7
8
9
10
Time [min]
100
150
200
250
300
350
400
m/z
383.30
312.09
298.13
267.20
213.05
171.19
117.08
432.39
383.31
312.13
297.95
267.11
199.06
171.09
340.24
383.27
312.16
298.10
267.14
213.15
185.17
366.23
383.19
311.97
298.01
267.20
209.17
171.21
353.05
383.25
312.10
298.09
267.25
213.09
171.14
366.25
383.27
312.14
298.13
253.23
199.16
171.14
366.23

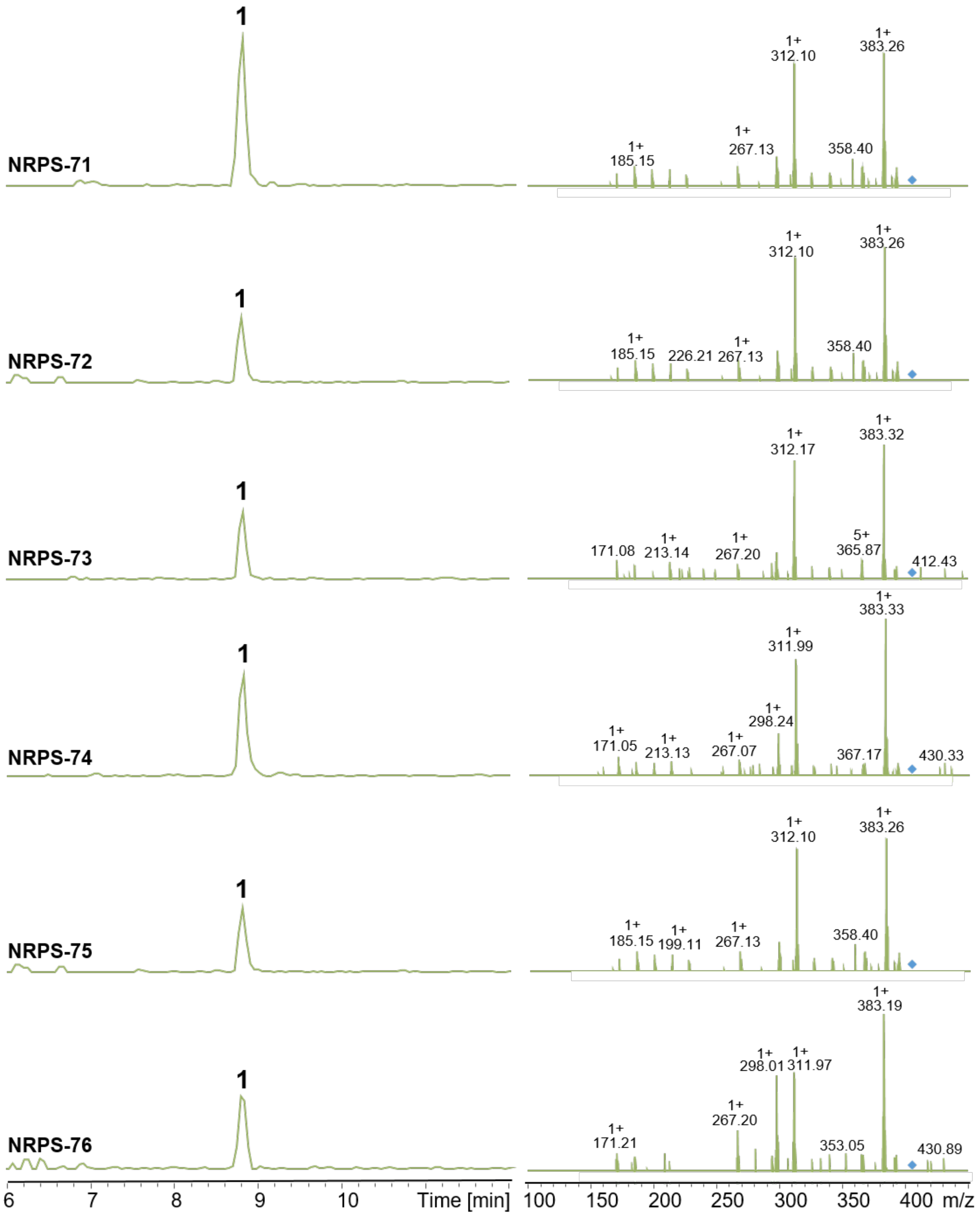
NRPS-71
1
1+ 185.15
1+ 267.13
1+ 312.10
358.40
1+ 383.26
NRPS-72
1
1+ 185.15
226.21
1+ 267.13
1+ 312.10
358.40
1+ 383.26
NRPS-73
1
171.08
1+ 213.14
1+ 267.20
1+ 312.17
5+ 365.87
1+ 383.32
412.43
NRPS-74
1
1+ 171.05
1+ 213.13
1+ 267.07
1+ 298.24
1+ 311.99
367.17
1+ 383.33
430.33
NRPS-75
1
1+ 185.15
1+ 199.11
1+ 267.13
1+ 312.10
358.40
1+ 383.26
NRPS-76
1
1+ 171.21
1+ 267.20
1+ 298.01
1+ 311.97
353.05
1+ 383.19
430.89
6 7 8 9 10 Time [min]
100 150 200 250 300 350 400 m/z

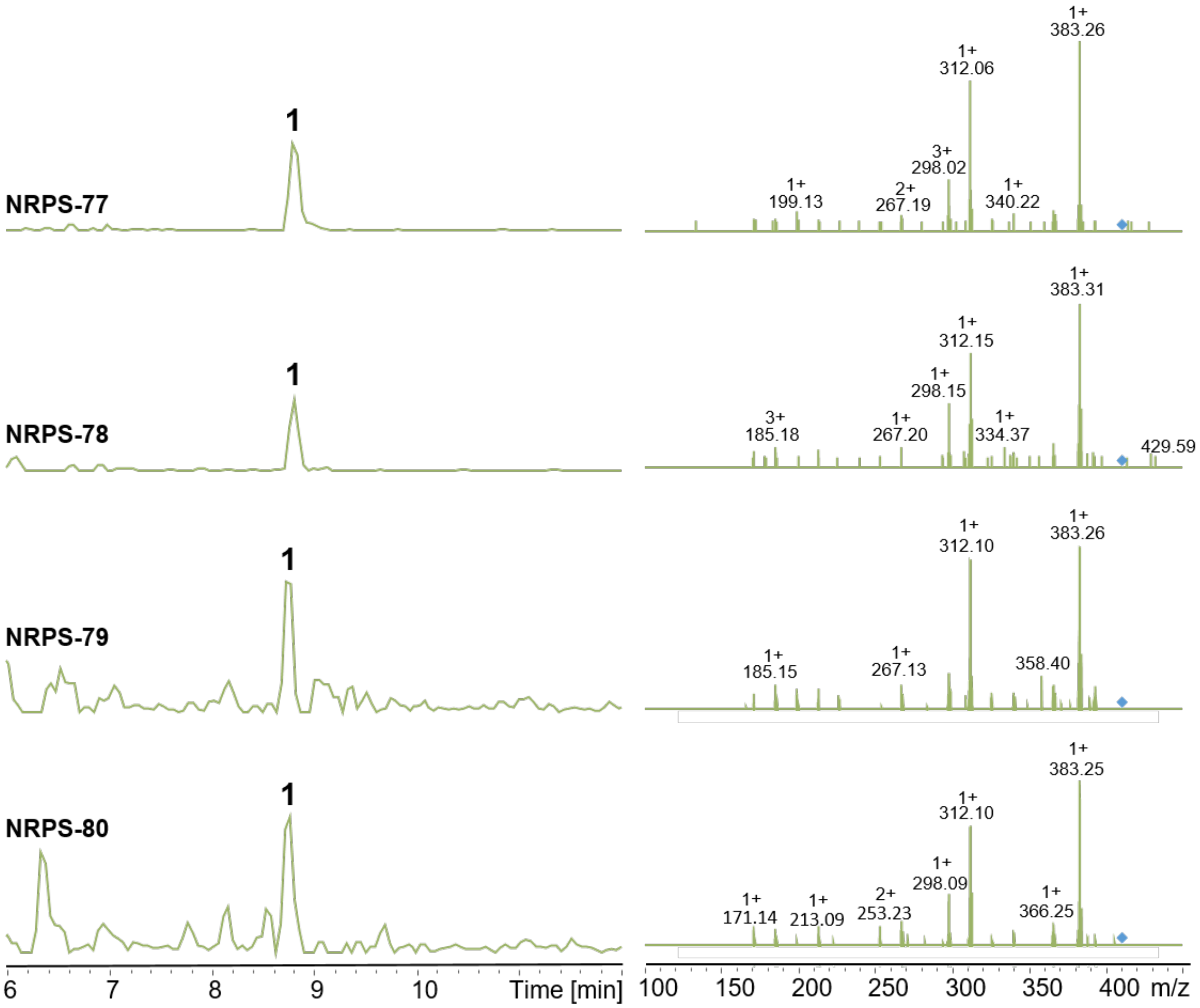

**Figure S23. HPLC/MS data (Figure S28) of compound 1 produced in *E. coli* DH10B::*mtaA*.** EIC/MS$^2$ of 1 (m/z $[M+H]^+$ = 411.29) produced by NRPS-1 and NRPS-66 to NRPS-80.

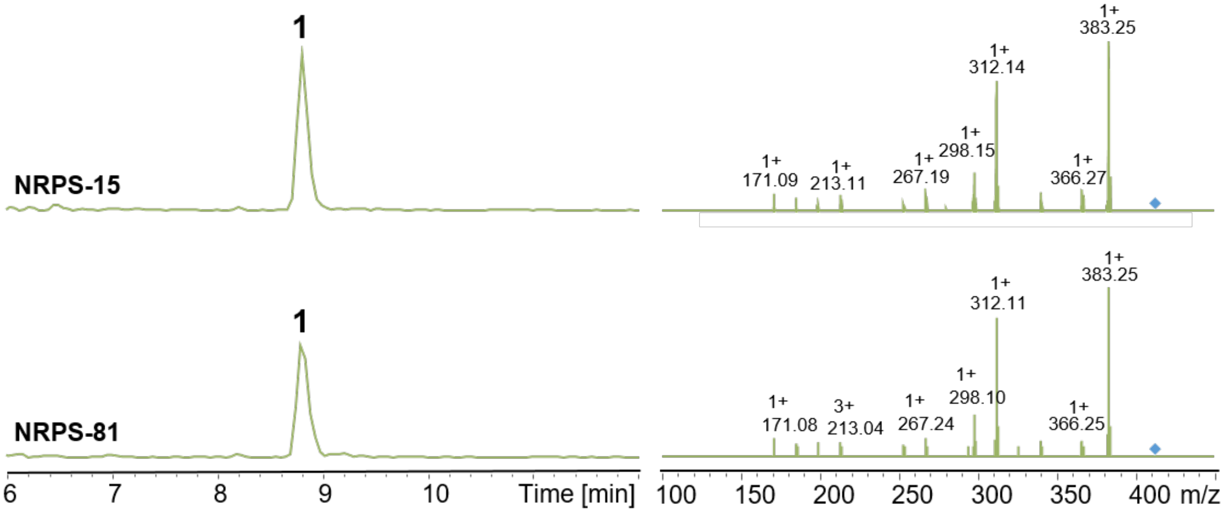

**Figure S24. HPLC/MS data (Figure S29) of compound 1 produced in *E. coli* DH10B::*mtaA*.** EIC/MS$^2$ of 1 (m/z $[M+H]^+$ = 411.29) produced by NRPS-15 and NRPS-81.

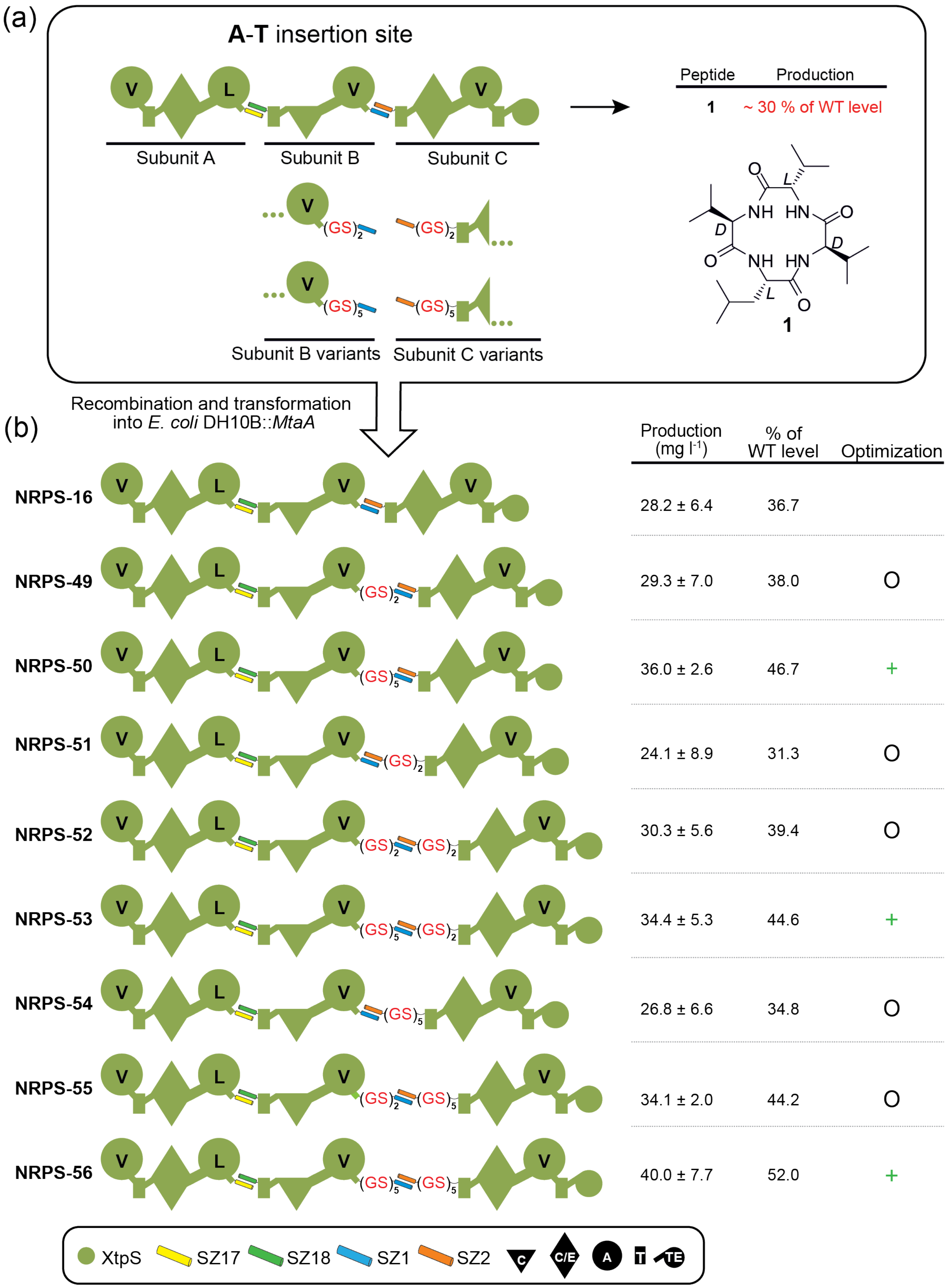
(a)
A-T insertion site
V
L
V
V
Subunit A
Subunit B
Subunit C
Subunit B variants
Subunit C variants
(GS)2
(GS)5
Peptide
Production
1
~ 30 % of WT level
NH
HN
O
L
D
1
Recombination and transformation into E. coli DH10B::MtaA
(b)
Production (mg l-1)
% of WT level
Optimization
NRPS-16
28.2 ± 6.4
36.7
NRPS-49
29.3 ± 7.0
38.0
O
NRPS-50
36.0 ± 2.6
46.7
+
NRPS-51
24.1 ± 8.9
31.3
O
NRPS-52
30.3 ± 5.6
39.4
O
NRPS-53
34.4 ± 5.3
44.6
+
NRPS-54
26.8 ± 6.6
34.8
O
NRPS-55
34.1 ± 2.0
44.2
O
NRPS-56
40.0 ± 7.7
52.0
+
XtpS
SZ17
SZ18
SZ1
SZ2
C
C/E
A
T
TE

**Figure S25. More GS-optimized chimeric tri-partite XtpS NRPSs split at the A-T position.** (a) Between each experimental approach, the production of non-optimized NRPS-16 varies, but is on average at ~30% of WT level. A set of modified subunit B and C variants were constructed by inserting GS stretches of varying length (4 AAs or 10 AAs) between subunit 2 and SZ1 and subunit 3 and SZ2. (b) Generated modified subunits were re-combined with non-modified subunits and transformed into *E. coli* DH10B::*MtaA* to obtain NRPS-49 to -56. Production titres of NRPS-49 to -56 were compared with each other and rated with from –, – –, – – – to O, +, ++, +++. Corresponding peptide yields (mg/L) and standard deviations are obtained from biological triplicate experiments. For domain assignment, the following symbols are used: (A, large circles), (T, rectangle), (C, triangle), (C/E, diamond), (TE, small circle); substrate specificities are assigned for all A domains and indicated by capital letters.

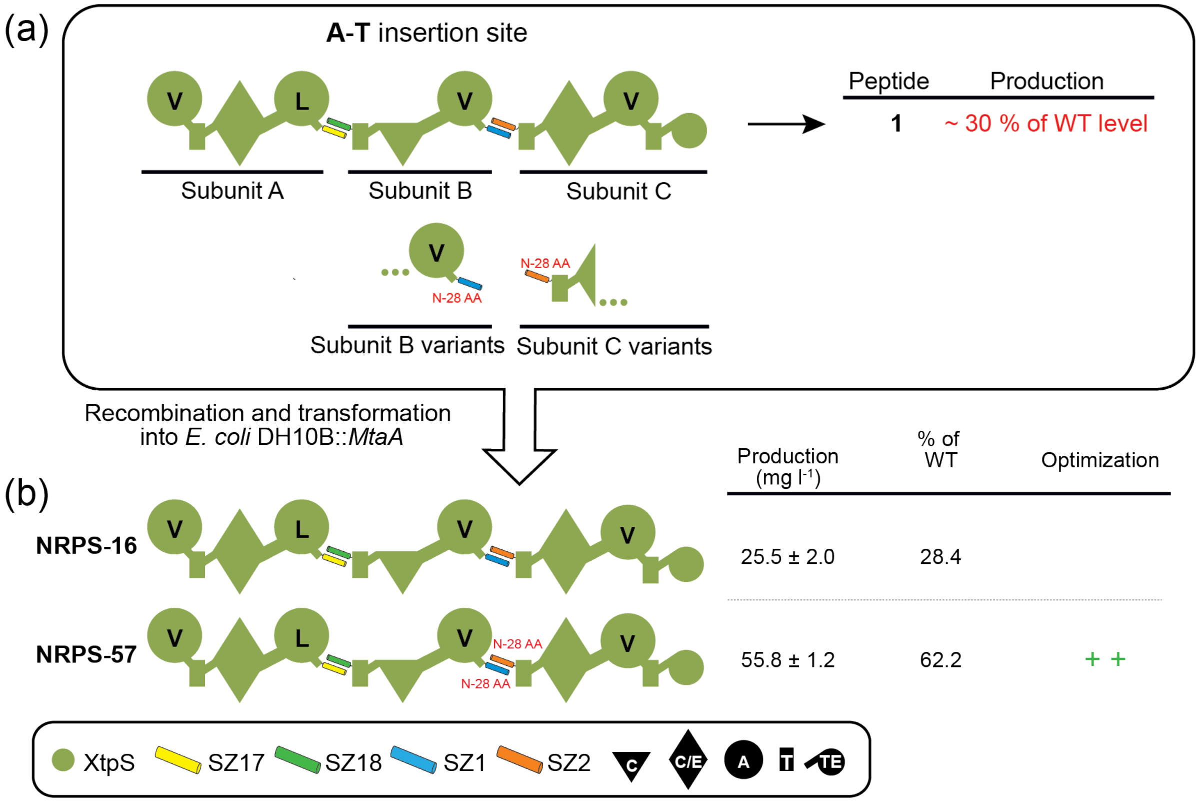

**Figure S26. SZ1:2 truncation of chimeric di-partite XtpS NRPSs split at the A-T position.** (a) Between each experimental approach, the production of non-optimized NRPS-16 varies, but is on average at ~30% of WT level. Subunit B and C variants were N-terminally truncated by 28 AA, respectively. (b) Generated modified subunits were re-combined with non-modified subunits and transformed into *E. coli* DH10B::*MtaA* to obtain NRPS-16 and -57.Corresponding peptide yields (mg/L) and standard deviations are obtained from biological triplicate experiments. Rating of production titres and domain assignment is as described before.

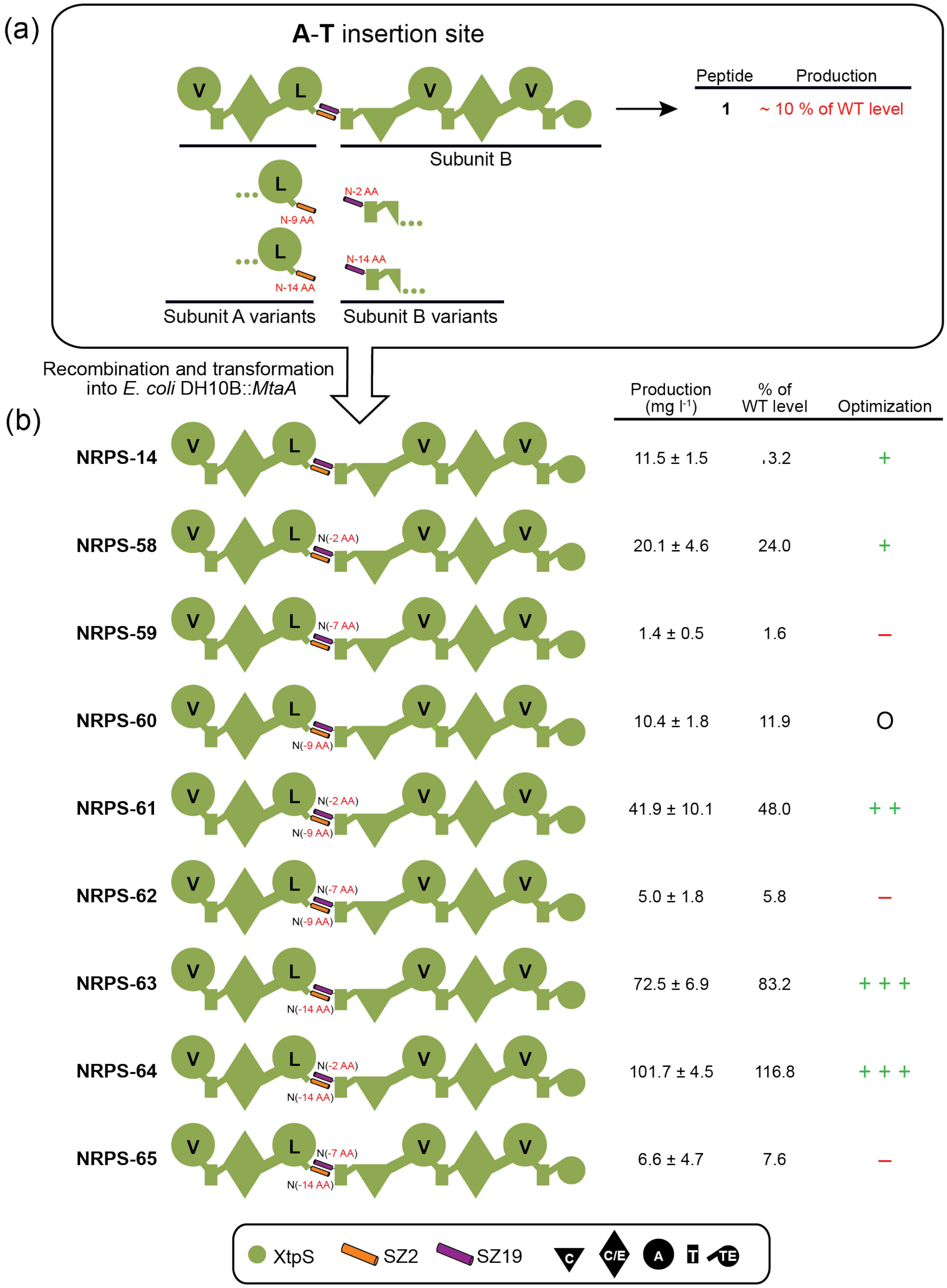
(a)
A-T insertion site
V
L
V
V
Subunit B
Peptide
Production
1
~ 10 % of WT level
L
N-9 AA
N-2 AA
L
N-14 AA
N-14 AA
Subunit A variants
Subunit B variants
Recombination and transformation
into E. coli DH10B::MtaA
(b)
Production (mg l-1)
% of WT level
Optimization
NRPS-14 11.5 ± 1.5 13.2 +
NRPS-58 N(-2 AA) 20.1 ± 4.6 24.0 +
NRPS-59 N(-7 AA) 1.4 ± 0.5 1.6 –
NRPS-60 N(-9 AA) 10.4 ± 1.8 11.9 O
NRPS-61 N(-2 AA) N(-9 AA) 41.9 ± 10.1 48.0 + +
NRPS-62 N(-7 AA) N(-9 AA) 5.0 ± 1.8 5.8 –
NRPS-63 N(-14 AA) 72.5 ± 6.9 83.2 + + +
NRPS-64 N(-2 AA) N(-14 AA) 101.7 ± 4.5 116.8 + + +
NRPS-65 N(-7 AA) N(-14 AA) 6.6 ± 4.7 7.6 –
XtpS
SZ2
SZ19
C
C/E
A
T
TE

**Figure S27. SZ2:19 truncation of chimeric di-partite XtpS NRPSs split at the A-T position**. (a) Between each experimental approach, the production of non-optimized NRPS-14 varies, but is on average at ~10% of WT level. A set of modified subunit A and B variants were constructed by N-terminally truncating SZ2 by 9 AAs and 14 AAs and SZ19 by 2 AAs and 7 AAs, respectively. (b) Generated modified subunits were re-combined with non-modified subunits and transformed into E. coli DH10B::MtaA to obtain NRPS-14, -59 to -66. Corresponding peptide yields (mg/L) and standard deviations are obtained from biological triplicate experiments. Rating of production titres and domain assignment is as described before.

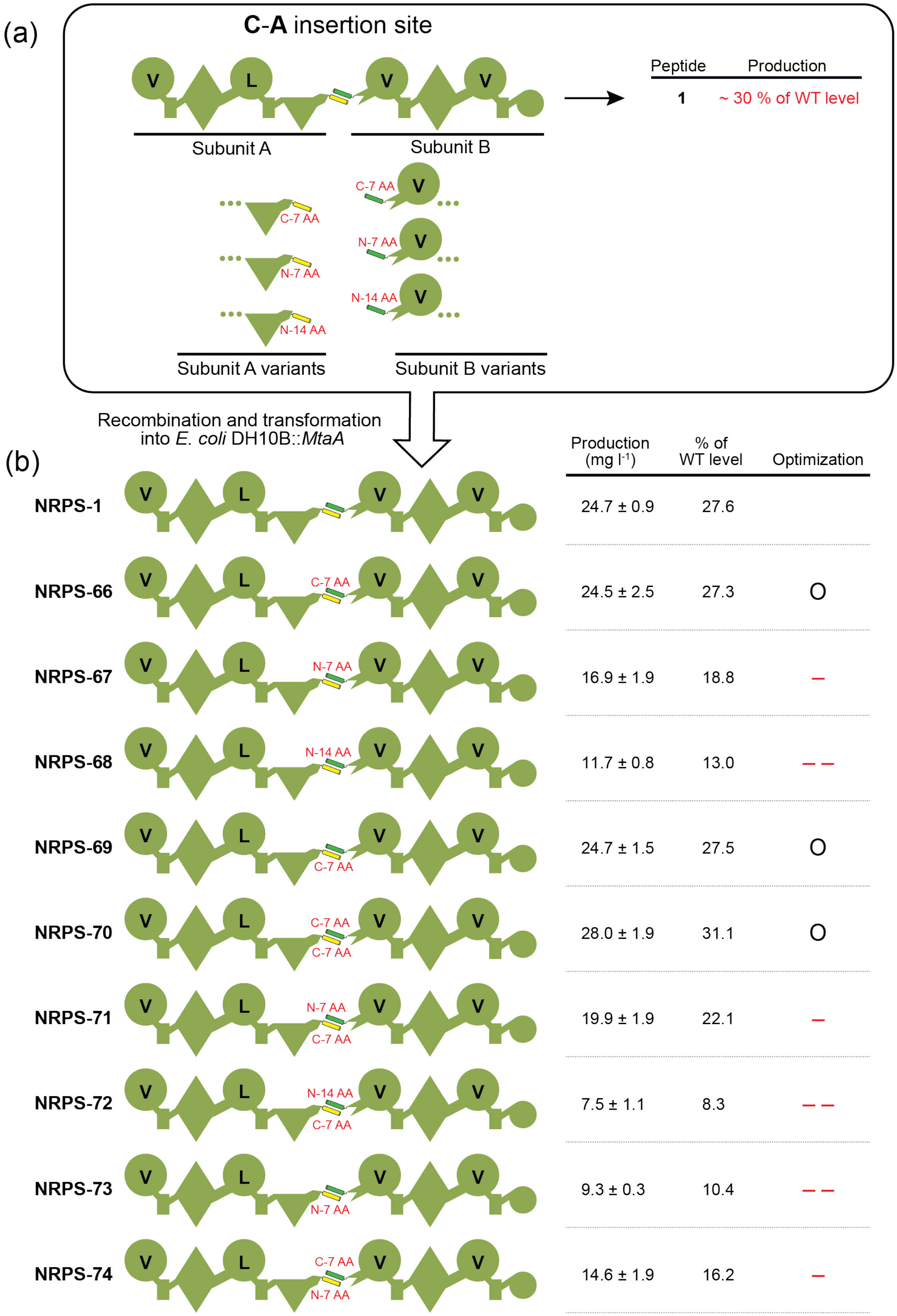

| | Production (mg l$^{-1}$) | % of WT level | Optimization |
|---|---|---|---|
| NRPS-1 | 24.7 ± 0.9 | 27.6 | |
| NRPS-66 | 24.5 ± 2.5 | 27.3 | O |
| NRPS-67 | 16.9 ± 1.9 | 18.8 | – |
| NRPS-68 | 11.7 ± 0.8 | 13.0 | – – |
| NRPS-69 | 24.7 ± 1.5 | 27.5 | O |
| NRPS-70 | 28.0 ± 1.9 | 31.1 | O |
| NRPS-71 | 19.9 ± 1.9 | 22.1 | – |
| NRPS-72 | 7.5 ± 1.1 | 8.3 | – – |
| NRPS-73 | 9.3 ± 0.3 | 10.4 | – – |
| NRPS-74 | 14.6 ± 1.9 | 16.2 | – |

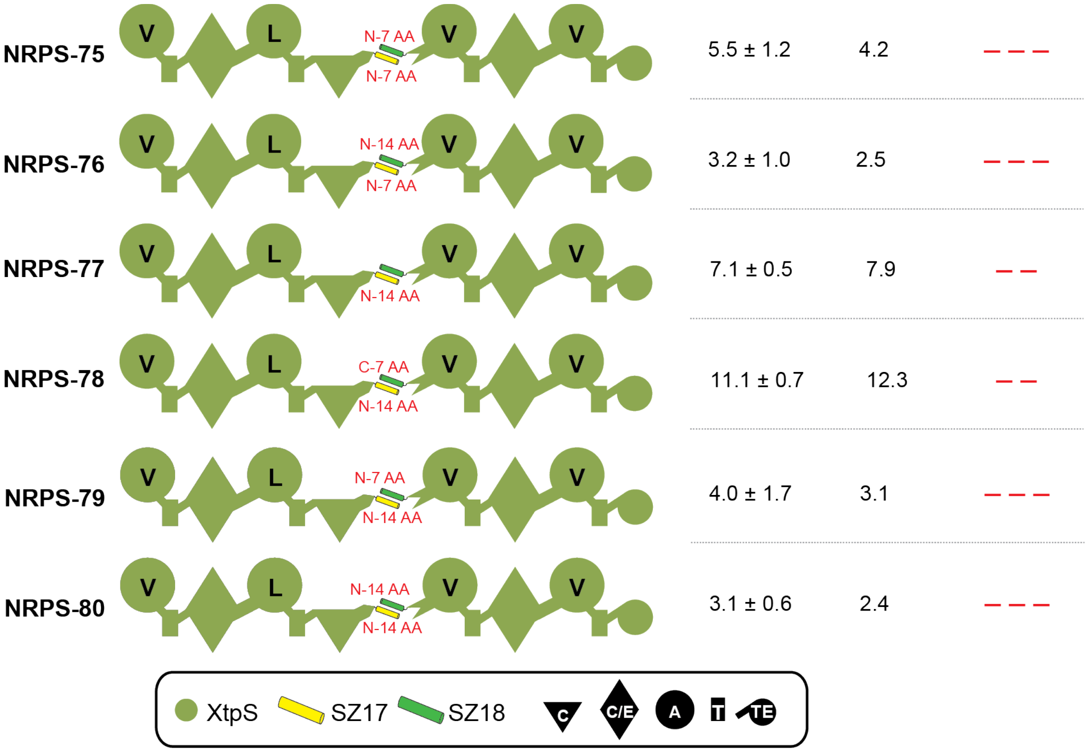

**Figure S28. SZ17:17 truncation of chimeric di-partite XtpS NRPSs split at the C-A position**. (a) Between each experimental approach, the production of non-optimized NRPS-1 varies, but is on average at ~30% of WT level. A set of modified subunit A and B variants were constructed by N- terminally truncating SZ17 and SZ18 by 7 AAs and 14 AAs, respectively and C-terminally truncating SZ17 and SZ18 by 7 AAs. (b) Generated modified subunits were re-combined with non-modified subunits and transformed into E. coli DH10B::MtaA to obtain NRPS-1, -67 to -81. Corresponding peptide yields (mg/L) and standard deviations are obtained from biological triplicate experiments. Rating of production titres and domain assignment is as described before.

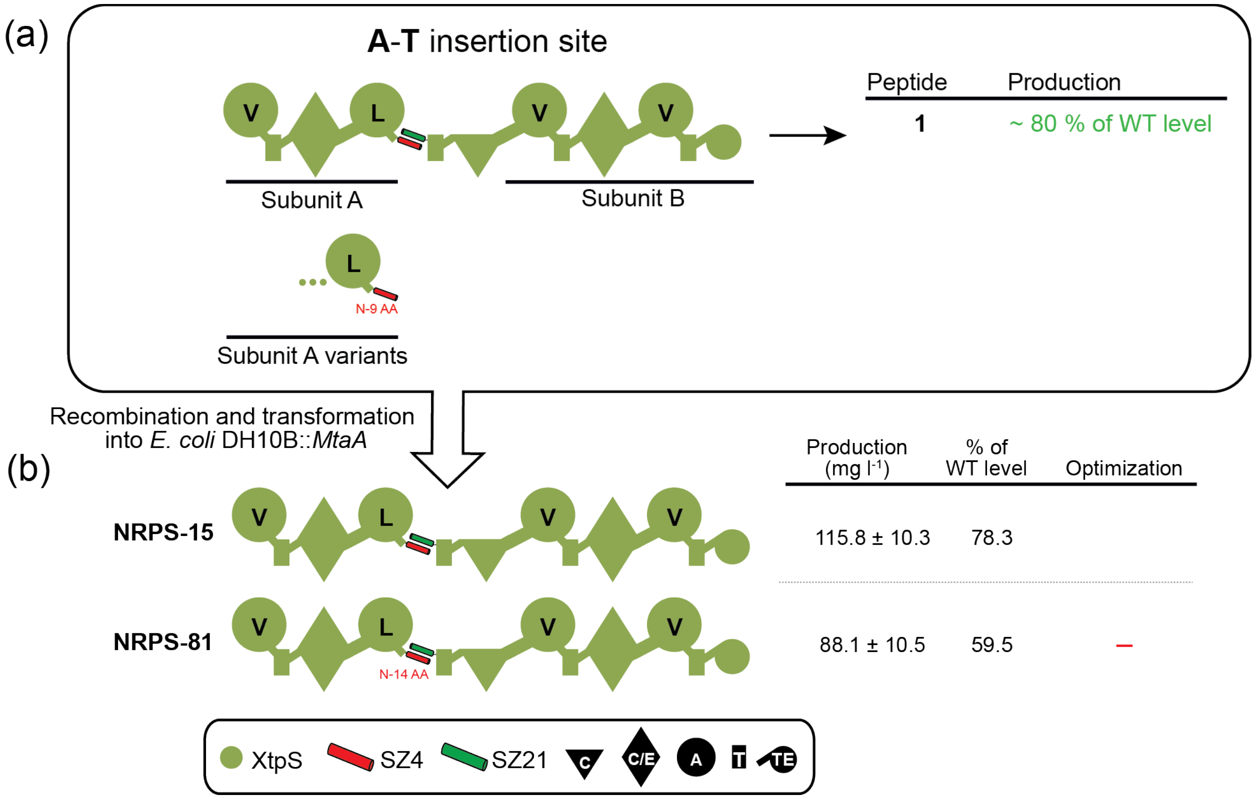

**Figure S29. SZ4:21 truncation of chimeric di-partite XtpS NRPSs split at the A-T position.** (a) Between each experimental approach, the production of non-optimized NRPS-15 varies, but is on average at ~30% of WT level. Modified subunit A was constructed by N- terminally truncating SZ4 by 14 AAs. (b) Generated modified subunits were re-combined with non-modified subunits and transformed into E. coli DH10B::MtaA to obtain NRPS-1, -67 to -81. Corresponding peptide yields (mg/L) and standard deviations are obtained from biological triplicate experiments. Rating of production titres and domain assignment is as described before.

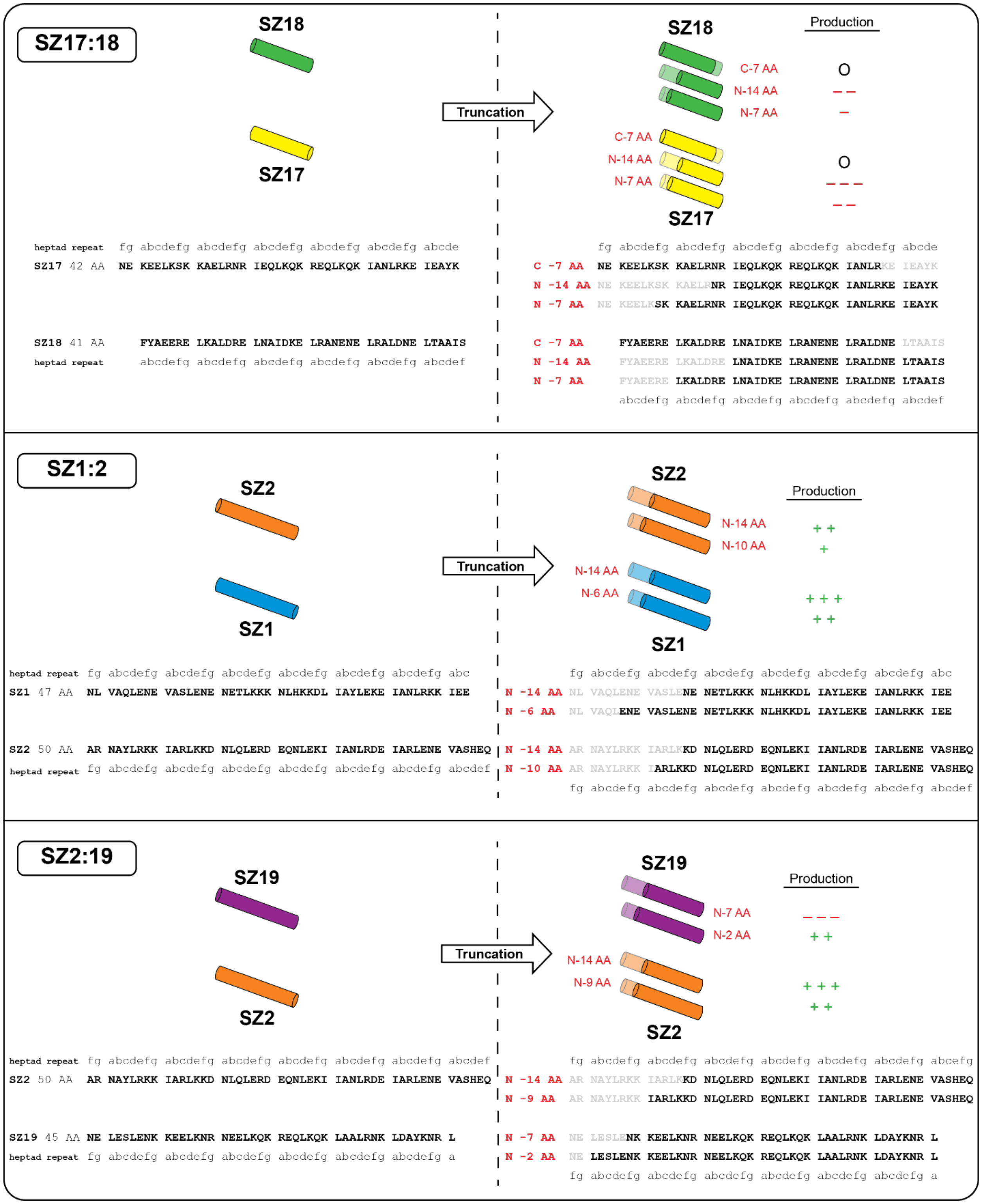
SZ17:18
SZ18
SZ17
Truncation
Production
C-7 AA
N-14 AA
N-7 AA
O
heptad repeat fg abcdefg abcdefg abcdefg abcdefg abcdefg abcde
SZ17 42 AA NE KEELKSK KAELRNR IEQLKQK REQLKQK IANLRKE IEAYK
C -7 AA NE KEELKSK KAELRNR IEQLKQK REQLKQK IANLRKE IEAYK
N -14 AA NE KEELKSK KAELRNR IEQLKQK REQLKQK IANLRKE IEAYK
N -7 AA NE KEELKSK KAELRNR IEQLKQK REQLKQK IANLRKE IEAYK
SZ18 41 AA FYAEERE LKALDRE LNAIDKE LRANENE LRALDNE LTAAIS
heptad repeat abcdefg abcdefg abcdefg abcdefg abcdefg abcdef
C -7 AA FYAEERE LKALDRE LNAIDKE LRANENE LRALDNE LTAAIS
N -14 AA FYAEERE LKALDRE LNAIDKE LRANENE LRALDNE LTAAIS
N -7 AA FYAEERE LKALDRE LNAIDKE LRANENE LRALDNE LTAAIS
abcdefg abcdefg abcdefg abcdefg abcdefg abcdef
SZ1:2
SZ2
SZ1
Truncation
Production
N-14 AA
N-10 AA
N-6 AA
+ +
+
+ + +
heptad repeat fg abcdefg abcdefg abcdefg abcdefg abcdefg abcdefg abc
SZ1 47 AA NL VAQLENE VASLENE NETLKKK NLHKKDL IAYLEKE IANLRKK IEE
N -14 AA NL VAQLENE VASLENE NETLKKK NLHKKDL IAYLEKE IANLRKK IEE
N -6 AA NL VAQLENE VASLENE NETLKKK NLHKKDL IAYLEKE IANLRKK IEE
SZ2 50 AA AR NAYLRKK IARLKKD NLQLERD EQNLEKI IANLRDE IARLENE VASHEQ
heptad repeat fg abcdefg abcdefg abcdefg abcdefg abcdefg abcdefg abcdef
N -14 AA AR NAYLRKK IARLKKD NLQLERD EQNLEKI IANLRDE IARLENE VASHEQ
N -10 AA AR NAYLRKK IARLKKD NLQLERD EQNLEKI IANLRDE IARLENE VASHEQ
fg abcdefg abcdefg abcdefg abcdefg abcdefg abcdefg abcdef
SZ2:19
SZ19
SZ2
Truncation
Production
N-7 AA
N-2 AA
N-14 AA
N-9 AA
+ +
+ + +
heptad repeat fg abcdefg abcdefg abcdefg abcdefg abcdefg abcdefg abcdef
SZ2 50 AA AR NAYLRKK IARLKKD NLQLERD EQNLEKI IANLRDE IARLENE VASHEQ
fg abcdefg abcdefg abcdefg abcdefg abcdefg abcdefg abcefg
N -14 AA AR NAYLRKK IARLKKD NLQLERD EQNLEKI IANLRDE IARLENE VASHEQ
N -9 AA AR NAYLRKK IARLKKD NLQLERD EQNLEKI IANLRDE IARLENE VASHEQ
SZ19 45 AA NE LESLENK KEELKNR NEELKQK REQLKQK LAALRNK LDAYKNR L
heptad repeat fg abcdefg abcdefg abcdefg abcdefg abcdefg abcdefg a
N -7 AA NE LESLENK KEELKNR NEELKQK REQLKQK LAALRNK LDAYKNR L
N -2 AA NE LESLENK KEELKNR NEELKQK REQLKQK LAALRNK LDAYKNR L
fg abcdefg abcdefg abcdefg abcdefg abcdefg abcdefg a

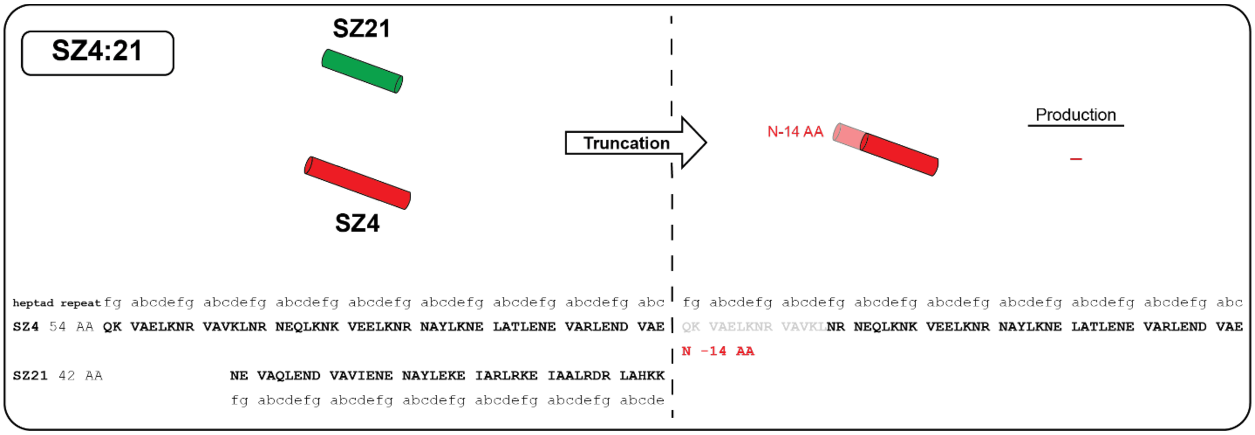

**Figure S30. Overview of truncated SZs.** Left row: schematic representation of SZs and their corresponding AAs sequences. Right row. Truncated SZs variants and corresponding AAs sequence. Impact of truncated SZs on type S NRPSs is indicated and rated from –, – –, – – – to O, +, ++, +++.
